# Transposon end recognition and excision mechanisms of type I-F CRISPR-associated transposases

**DOI:** 10.64898/2026.05.05.722991

**Authors:** Mateusz Walter, Giada Finocchio, Seraina Oberli, Iana C. Hammerschmid, George D. Lampe, Julia Karan, Thomas Swartjes, Samuel H. Sternberg, Martin Jinek, Irma Querques

## Abstract

CRISPR-associated transposons (CASTs) are Tn7-like elements that have co-opted RNA-guided CRISPR effectors for targeted DNA insertion. CASTs have been adapted as genome editing tools for programmable, site-specific integration. Among them, the type I-F system from *Ps*e*udoalteromonas* (*Pse*CAST) shows uniquely robust activity in human cells, yet its mechanistic basis remains poorly understood. Here, we present structural and biochemical analysis of the *Pse*CAST transposase TnsAB. Biochemical reconstitution of transposon DNA excision defines key characteristics of the transposition mechanism. Cryogenic electron microscopy (cryo-EM) structures of *Pse*TnsAB paired-end complexes reveal molecular determinants of transpososome assembly, transposon end recognition and cleavage. We validate these findings using biochemical and *in vivo* assays of structure-based transposase mutants, and provide mechanistic insights into the enhanced activity of a laboratory-evolved TnsAB variant. Together, our studies highlight molecular features underlying the efficiency of natural and engineered type I-F transposases and establish a mechanistic framework for their continued rational optimization.

## Introduction

Transposable elements are mobile DNA segments that reshape genomes across all domains of life. Their transposition activity drives genomic plasticity, remodeling, and horizontal gene transfer^1^. DNA transposons are typically demarcated by end sequences containing inverted repeat motifs that are recognized and cleaved by transposase enzymes during the transposition process. Most DNA transposons move via a cut-and-paste mechanism, where the transposase synapses the transposon ends into a paired-end complex (PEC), excises the element from donor DNA, and integrates it into a target locus through target capture and strand transfer^2^.

The Tn7 transposon from *Escherichia coli* (*Ec*Tn7) is a model for targeted cut-and-paste DNA transposition, relying on the transposase subunits TnsA and TnsB, the AAA+ ATPase TnsC, and the target-selector proteins TnsD and TnsE^3,4^. Complete excision requires the complementary activities of the type II restriction endonuclease-like TnsA and the DDE-family transposase TnsB, which generate the 5′ and 3′ termini, respectively, of the excised left (LE) and right end (RE). TnsB catalyzes strand transfer to complete integration in the target DNA^5-7^ (**Fig. 1a**). TnsC bridges the transposase to the target selector, mediating target capture and regulating transpososome activity by ATP-dependent oligomerization^8-10^. TnsD, a member of the TniQ protein family, directs insertions into the chromosomal *att*Tn7 site^8^, whereas TnsE biases integration into conjugative plasmids, promoting horizontal gene transfer^4,11-13^.

**Fig. 1.**
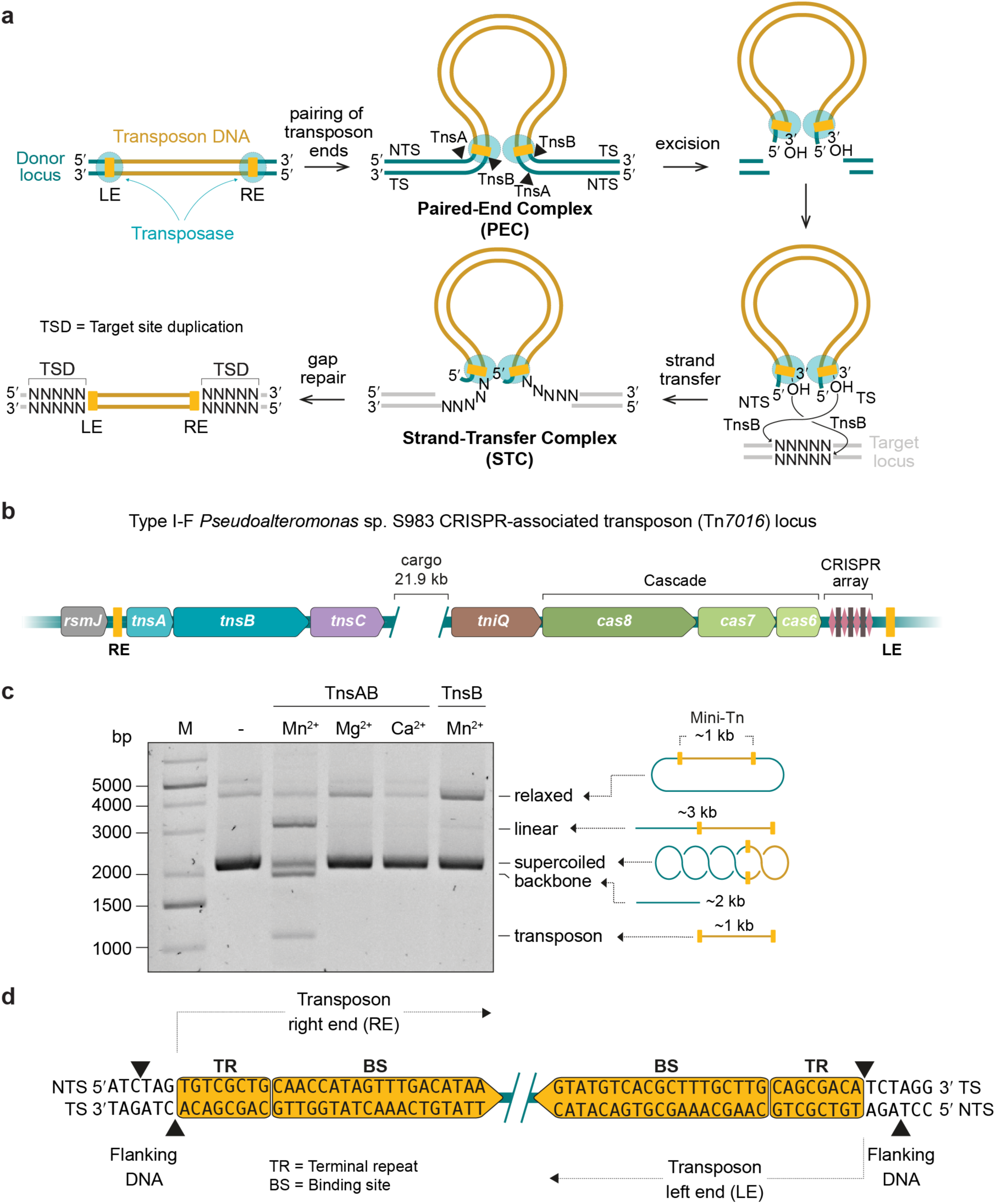
Biochemical analysis of the type I-F CRISPR-associated transposase *Pse*TnsAB. (**a**) Model of cut-and-paste transposition mediated by TnsA-TnsB transposases in Tn7-like elements. Transposases recognize inverted repeats at transposon ends, assemble a paired-end complex (PEC), fully excise the element via nucleophilic attacks on both strands, and integrate it into a target site via direct 3′-OH strand transfer. Gap repair generates target site duplications (TSDs). LE/RE, left/right end; NTS, non-transferred strand; TS, transferred strand. (**b**) Genomic organization of the type I-F CRISPR-associated Tn*7016* transposon system from *Pseudoalteromonas* sp. S983 (*Pse*CAST). The CRISPR array consists of alternating repeats (diamonds), and spacers (squares). The element is inserted downstream of the *rsmJ* gene. (**c**) *In vitro* excision assay monitoring DNA cleavage activity of *Pse*TnsAB in the presence of divalent metal cations. DNA species are resolved by agarose gel electrophoresis. Results of a representative experiment are shown. Quantification of three independent experiments is provided in **Supplementary Fig. 2b**. -, no-protein control. Mini-Tn, mini-transposon. M, marker. (**d**) Schematic representation of cleavage positions inferred from next-generation sequencing (NGS) results. TR, terminal repeat. BS, TnsB-binding site. Black triangles indicate predominant cleavage positions located near the transposon ends, as determined by sequencing of nested PCR products. NGS data are shown in **Supplementary Fig. 2e**.

CRISPR-associated transposons (CASTs) are Tn7-like elements that use nuclease-less or catalytically inactive CRISPR effectors for target DNA selection^14-16^ (**Fig. 1b**). CRISPR array-encoded RNA guides (crRNAs) direct DNA recognition by the CRISPR-Cas machinery, which recruits transposase components to the target DNA via the zinc-finger protein TniQ, thereby coupling RNA-guided recognition with transposon insertion downstream of the target recognition site^17-20^. CASTs have been repurposed as genome editing tools, enabling programmable, site-specific integration of large DNA payloads^21-29^.

The two major types of CAST systems differ in their CRISPR effectors and transpososome architectures. Whereas type I CASTs use a crRNA-guided multi-subunit Cascade-like complex for targeting^17,19^, type V-K CASTs rely on the pseudonuclease Cas12k, loaded with a crRNA-tracrRNA (trans-activating CRISPR RNA) dual guide, along with the host ribosomal protein S15^18,30^. In type I CASTs, transposition is catalyzed by TnsA and TnsB, which can be encoded either as separate proteins or a naturally fused single polypeptide, and is thought to proceed via a cut-and-paste mechanism. In contrast, type V-K systems contain TnsB and lack TnsA, which dictates a copy-and-paste mechanism that predominantly generates cointegrate products comprising duplicated transposons flanking the donor backbone^31,32^. CASTs are further subdivided into subtypes, including I-F^17^, I-B (I-B1 and I-B2)^19^, I-C^33^, I-D^20^, I-E^34^, and V-K^18^, with both their CRISPR and Tns(A)BC machineries being highly divergent^33,35^. Functional insights into CAST activity have so far been obtained mainly from insertion assays in heterologous hosts, which show that type V-K systems generally have limited specificity^18,23,36,37^ and generate heterogeneous transposition products dominated by cointegrates^24,32^. By contrast, type I-F CASTs consistently achieve high specificity and fidelity, producing precise simple integrations with markedly greater efficiency than type I-B, I-D, or V-K systems^17-20,26,36^.

Structural studies of the type V-K CAST system from *Scytonema hofmannii* (*Sh*CAST) revealed molecular details of Cas12k-mediated target recognition^38,39^ and TniQ- and S15-dependent TnsC oligomerization^30,40^, and captured TnsB in strand-transfer complex (STC) states, either in isolation^41,42^ or within megadalton holocomplexes including Cas12k, TniQ, TnsC, and S15^43,44^. In turn, structural information for type I-F and I-B2 CASTs includes TniQ-Cascade and TnsC assemblies^45-52^ as well as a recently reported transpososome structure of a type I-B2 CAST from *Peltigera membranacea cyanobiont* (*Pmc*CAST) revealing the mechanism of TnsD-dependent recruitment of the transposase to the *att* site^53^. Beyond CASTs, the structure of *Ec*Tn7 TnsB bound to repetitive sites in the transposon end DNA has revealed oligomerization-dependent mechanism of transposon end recognition prior to PEC formation^54^.

The lack of structural information on transposase architecture has limited our mechanistic understanding of type I-F CASTs. Among these, the Tn*7016* system from *Pseudoalteromonas* sp. S983^36^ (*Pse*CAST) (**Fig. 1b**) shows uniquely robust insertion activity in human cells^26,27^. Despite weak Cascade-mediated targeting, it integrates more efficiently than other systems, suggesting that its transposase has particularly high activity^26,52^. Directed evolution of *Pse*CAST components in *E. coli* has further increased integration rates in the human genome up to 10-30% without compromising product purity, although off-target activity was detected^27^. Key mutations map to TnsB, establishing the transposase as the primary driver of efficiency, fidelity and specificity.

Here, we provide mechanistic insights into the *Pse*CAST transposase. Using cryogenic electron microscopy (cryo-EM), we determined a set of structures of the heteromeric TnsA-TnsB transposase assembled on DNAs mimicking cleaved transposon ends, thus representing the post-excision product state. Our structural and biochemical data support a model in which *Pse*CAST transposition proceeds through highly specific recognition of transposon ends, pairing driven by transposase oligomerization, and coordinated DNA cleavage by TnsA and TnsB. Overall, these findings establish the molecular basis of type I-F CAST activity and reveal molecular features that explain the unique performance of both natural and engineered variants.

## Results

### Biochemical analysis of *Pse*TnsAB

To investigate the biochemical activity and molecular assembly of the type I-F transposase TnsA-TnsB, we recombinantly expressed and purified it as a single artificial fusion protein (*Pse*TnsAB) (**Supplementary Fig. 1a**), previously shown to retain full activity *in vivo*^26,32^. Size exclusion chromatography coupled with multi-angle light scattering (SEC-MALS) indicated that the protein is predominantly monomeric in isolation, displaying a molecular weight of 97.0 kDa (**Supplementary Fig. 1b**). To analyze the DNA cleavage activity of *Pse*TnsAB, we established an *in vitro* transposon excision assay (**Fig.1c**, **Supplementary Fig. 2a**). A supercoiled DNA plasmid containing a ∼1 kb cargo segment flanked by LE and the RE sequences was incubated with *Pse*TnsAB in the presence of different divalent metal cations. In reactions supplemented with Mn^2+^, known to relax the catalytic constraints of DDE/D enzymes *in vitro*^55^, we observed product fragments corresponding to a fully excised transposon and a ∼2 kb linearized plasmid backbone, along with linear intermediates corresponding to single-end cleavage events (**Fig. 1c**, **Supplementary Fig. 2b**). Such cleavage products were undetectable in the presence of Mg^2+^, the physiological catalytic ion, and with Ca^2+^, which is known to support assembly of cleavage-incompetent transposase complexes^56,57^. Mn^2+^-dependent activity is consistent with previous biochemical studies of the canonical *Ec*Tn7 transposon, where Mn^2+^ promotes cleavage by TnsA and TnsB alone, whereas Mg^2+^ supports catalysis only in the presence of TnsC and targeting factors^6,58^. In the absence of *Pse*TnsA, *Pse*TnsB (**Supplementary Fig. 1a**) exhibited processing activity that was reduced ∼3.8-fold compared to *Pse*TnsAB (**Fig. 1c**, **Supplementary Fig. 2b**), indicating that the TnsA subunit is required to support efficient nucleolytic function of TnsB in type I-F CASTs, in analogy with EcTn^76,8,59^.

To determine the precise cleavage positions at both transposon ends, cleavage products were ligated to single-stranded DNA oligonucleotide adapters via their phosphorylated 5′ ends. Nested PCR with plasmid- and adapter-specific primers enabled amplification of product fragments of discrete length (**Supplementary Fig. 2c**), which were then analyzed by next-generation sequencing. The results showed that *Pse*TnsAB produces staggered double-stranded breaks at both transposon ends, with one strand (the non-transferred strand, NTS) cleaved three nucleotides outside the transposon 5′ boundary and the complementary strand (the transferred strand, TS) cut exactly at the 3′ end of the transposon (**Fig. 1d**, **Supplementary Fig. 2d**-**e**). This cleavage pattern is consistent with the combined activities of TnsA and TnsB in *Ec*Tn7^58^. These findings indicate that artificial fusion of the transposase subunits and the use of manganese as a cofactor do not alter the cleavage pattern of the type I-F CAST transposase. They further demonstrate that transposition follows a cut-and-paste mechanism, as previously inferred from sequencing analysis of *in vivo* insertion products^32^.

The transposon ends of *Pse*CAST are defined by the presence of imperfect inverted repeats comprising an 8-bp terminal repeat (TR) sequence and an array of internal 19-bp TnsB-binding sites (hereafter referred to as “binding sites”, BSs). In a minimal configuration required for transposition in *E. coli*^26^, the RE contains three binding sites separated by 2 bp and 3 bp, while the LE harbors three sites separated by 36 bp and 4 bp (**Supplementary Fig. 2d**). To investigate the functional impact of this transposon end asymmetry, we conducted *in vitro* excision assays using mutant plasmid substrates with either symmetrized or truncated ends (**Supplementary Fig. 2f**-**g**). All substrates supported excision to varying degrees, with wild-type ends showing the highest activity. Compared to the native configuration, symmetrized RE-RE and LE-LE substrates retained approximately 12% and 27% excision activity, respectively. This result stands in contrast with *Ec*Tn7, in which the TnsB-binding sites within the RE are partially overlapping, resulting in functional asymmetry of the RE and LE, whereby two REs can support transposition, while two LEs cannot^54,60^. Although the LE harbors a predicted binding site for the bacterial integration host factor (IHF) in *Pse*CAST^61^, IHF was not required for excision under *in vitro* conditions. Plasmid substrates lacking internal BSs retained ∼19% excision activity, while deletion of the conserved TR nearly abolished excision (**Supplementary Fig. 2f**-**g**), consistent with its role in defining the precise TnsA and TnsB cleavage sites. Together, these results indicate critical functional features underlying excision in type I-F CASTs.

### Cryo-EM structure of *Pse*TnsAB paired-end-complex

To gain insights into the molecular mechanisms underpinning transposon end recognition and cleavage in type I-F CASTs, we proceeded to determine a cryo-EM structure of a *Pse*TnsAB paired-end complex (PEC). We initially focused on the RE, as previous studies showed that the presence of a minimum of two closely spaced binding sites is required for complex formation^41,42,53,54^. To facilitate structural analysis and minimize sample heterogeneity, we used a double-stranded DNA mimicking the RE excision product, comprising BS1 and 15 base pairs of BS2. The final design was informed by the structure of the type I-B2 CAST *Pmc*TnsABCD transpososome, in which density was resolved primarily for BS1 and only a portion of BS2^53^. Based on our biochemical data, the DNA included a 5′-phosphorylated 3-nucleotide overhang on the NTS (**Fig. 2a**). The DNA, hereafter referred to as PEC RE DNA, was incubated with *Pse*TnsAB (**Fig. 2b**) in the presence of 10 mM MgCl_2_, and complex formation was assessed by SEC (**Supplementary Fig. 3a**). To analyze the oligomeric state and determine the stoichiometry of *Pse*TnsAB, we performed mass photometry on both the apo and DNA-bound forms (**Supplementary Fig. 3b**). In the absence of DNA, the majority of *Pse*TnsAB was observed to exist in the monomeric state (molar mass of ∼93 kDa), along with a minor fraction of dimers (molar mass of ∼178 kDa). Upon addition of PEC RE DNA, a single species with a molar mass of ∼341 kDa was observed, corresponding to a complex comprising four *Pse*TnsAB protomers. Similar higher-order oligomeric species were also observed upon DNA addition with *Pse*TnsB alone (**Supplementary Fig. 3b**). These data indicate that transposon end DNA binding induces tetramerization of *Pse*TnsAB, likely promoting transposon end synapsis, in analogy with other related transposase systems^41,54,62^.

**Fig. 2.**
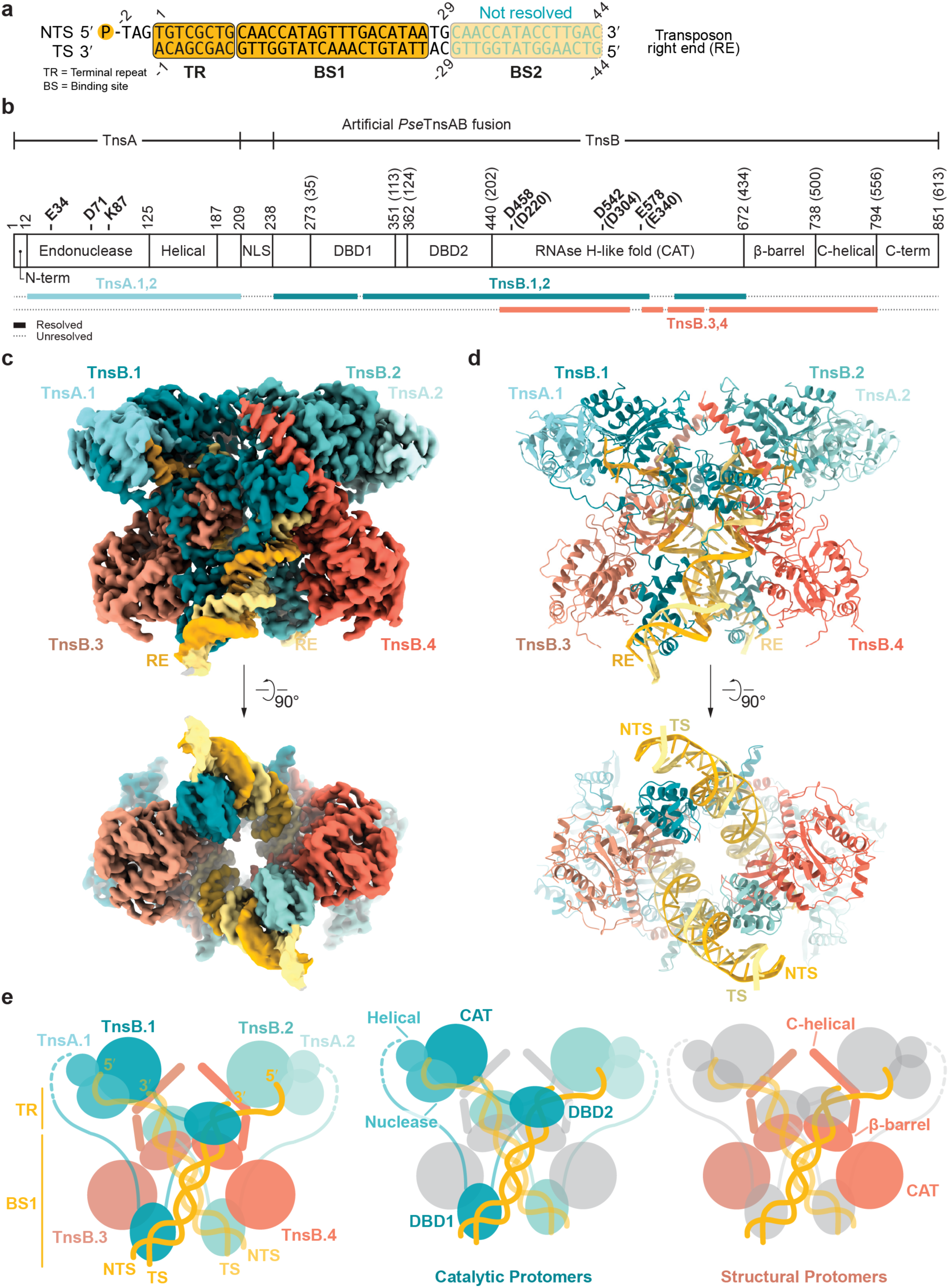
Cryo-EM structure of the *Pse*TnsAB paired-end complex on right end DNA. (**a**) Schematic of the transposon right end (RE) DNA used for paired-end complex (PEC) reconstitution, showing the TnsB-binding sites BS1 and BS2, and the terminal repeat (TR) sequences. NTS, non-transferred strand. TS, transferred strand. (**b**) Domain organization of *Pse*TnsAB artificial fusion protein. Catalytic residues are indicated, with numbering corresponding to the single TnsB protein shown in parentheses. Unresolved regions are indicated with dotted lines. NLS, nuclear localization signals. DBD, DNA binding domain. CAT, catalytic domain. N/C-term, N/C-terminal region. (**c**) Single-particle cryo-EM reconstruction (side and bottom views). Individual TnsAB are depicted in different colors, with structural protomers highlighted in shades of orange and catalytic protomers in shades of teal. The transposon right ends (RE) are colored yellow. (**d**) Atomic model (side and bottom views). Each TnsA and TnsB protomer is numbered. (**e**) Schematic representation of complex architecture (left) and domain organization of catalytic (middle) and structural (right) protomers.

We subsequently subjected the complex to single-particle analysis by cryo-EM and obtained a reconstruction, at an overall resolution of 2.78 Å (**Fig. 2c**, **Supplementary Fig. 4, Supplementary Table 1**), representing a tetrameric *Pse*TnsAB PEC assembled on DNA mimicking the cleaved transposon RE, i.e. in a post-excision, product-bound configuration (**Fig. 2d**). Overall, the structure exhibits 2-fold symmetry, revealing four copies of TnsB and two copies of TnsA assembled in an intertwined arrangement together with two copies of the RE excision product DNA (**Fig. 2d-e**). The full-length synthetic *Pse*TnsAB fusion protein consists of *Pse*TnsA and *Pse*TnsB fused via an artificial 29-residue spacer that includes two nuclear localization signals (NLSs)^26^. Each transposase subunit contains distinct domains connected by linker segments (**Fig. 2b**). *Pse*TnsA shares a comparable domain composition with TnsA proteins from *Ec*Tn7^63^ and the type I-B2 *Pmc*CAST^53^, comprising an N-terminal endonuclease followed by a helical subdomain. The endonuclease domain belongs to the type II restriction enzyme superfamily and harbors a conserved catalytic triad (Glu34^TnsA^, Asp71^TnsA^, Lys87^TnsA^). In turn, *Pse*TnsB consists of an N-terminal region subdivided into two DNA binding domains (DBD1 and DBD2). DBD1 contains a canonical helix-turn-helix (HTH) motif, while DBD2 features a winged helix-like (WH-like) fold. In *Ec*Tn7 TnsB and *Pmc*TnsB, DBD1 also includes an N-terminal SH3-like subdomain^53,54^; however, this subdomain is not conserved in *Pse*TnsB. Downstream of DBD2 is a catalytic domain (CAT) composed of an RNase H-like fold subdomain containing a catalytic triad (Asp220^TnsB^, Asp304^TnsB^, and Glu340^TnsB^) and a β-barrel subdomain, followed by a C-terminal helical region.

For each copy of the bound RE DNA, clear density is observed for a 28-bp duplex (NTS nucleotides dT-2–dG29 and TS nucleotides dA-1–dC-29) comprising the TR and BS1, as well as the 5′-phosphorylated TAG overhang on the NTS (**Fig. 2a**). Two TnsB subunits, hereafter referred to as TnsB.1 and TnsB.2, make the most extensive contacts with the transposon right end DNA and engage directly with TnsA.1 and TnsA.2, thereby functioning as the catalytic components of the complex. In both catalytic TnsB protomers, DBD1 and DBD2 are each associated with BS1, while the CAT domain is positioned *in trans* at the cleaved end of RE DNA copy (**Fig. 2e**). This *trans* configuration is a common feature of transpososomes, ensuring that DNA cleavage occurs only upon end recognition and PEC formation^2^. In contrast, the distal “structural” TnsB subunits, TnsB.3 and TnsB.4, contribute to complex assembly through extensive interactions with TnsB.1 and TnsB.2 (**Fig. 2d-e**). The DBDs of TnsB.3 and TnsB.4, along with the BS2 segments of the bound RE DNA, are not resolved, likely due to conformational flexibility.

The atomic model includes nearly the entire polypeptide sequence for both TnsA copies (residues Asn12–Pro209^TnsA^), while the artificial spacer connecting TnsA and TnsB is not resolved. In TnsB.1 and TnsB.2, residues Met1–Ser436^TnsB^, encompassing both DBDs and the catalytic domain, could be modeled, except for two flexible regions within the CAT domain (Lys79–Asp82^TnsB^ and Val348–Lys369^TnsB^). Residues Asp437–Glu613^TnsB^, including the β-barrel and the C-terminal helical region, of these subunits are not resolved in the reconstruction, likely due to conformational flexibility in the absence of additional DNA contacts (**Fig. 2b**). By contrast, in the type I-B2 *Pmc*TnsAB STC complex^53^, the corresponding regions are ordered in the catalytic protomers engaging the target DNA (**Supplementary Fig. 5**). TnsB.3 and TnsB.4 are clearly resolved from the CAT domains through the β-barrel and C-terminal helical region (residues Pro209–Ser557^TnsB^), except for a short flexible linker (Val330–Phe338^TnsB^) (**Fig. 2b**). While the type I-F and type I-B2 transposases share a similar overall architecture (**Supplementary Fig. 5**), their interaction interfaces differ substantially, as described below.

### Assembly of *Pse*TnsAB protomers

Complex assembly is primarily stabilized by interprotomer interactions between the catalytic and structural TnsB subunits, which adopt distinct conformations. The CAT domains of TnsB.3 and TnsB.4 are positioned above the DBD1 domains of TnsB.1 and TnsB.2, respectively, forming extensive interaction surfaces. A helix in the CAT domain, spanning residues Lys312–Ala321^TnsB^, interacts with the only helix in DBD1 that is not involved in DNA binding (**Fig. 3a**, **Supplementary Fig. 6a**). Within this interface, Glu315^TnsB^ and His316^TnsB^ in the CAT form a salt bridge and a T-shaped π–π interaction, respectively, with Lys52^TnsB^ and Tyr98^TnsB^ in DBD1. Although the Tyr-Arg/Lys motif in DBD1 is conserved in both *Ec*Tn7 TnsB^54^ and *Pmc*TnsB^53^, the corresponding helix in the CAT domain is not conserved, and the interactions it mediates are distinct (**Supplementary Fig. 5**). In addition to the CAT-DBD1 interaction, the β-barrel subdomains of TnsB.3 and TnsB.4 establish non-specific contacts with the protein backbone of DBD2 in TnsB.1 and TnsB.2, respectively (**Fig. 3b**, **Supplementary Fig. 6b**). These contacts are mediated by the charged residues Arg462^TnsB^ and Asp454^TnsB^, which are not conserved in *Ec*Tn7 TnsB or type I-B2 *Pmc*TnsB, underscoring distinct mechanisms of coupling transposon end recognition and oligomerization across TnsB orthologs.

**Fig. 3.**
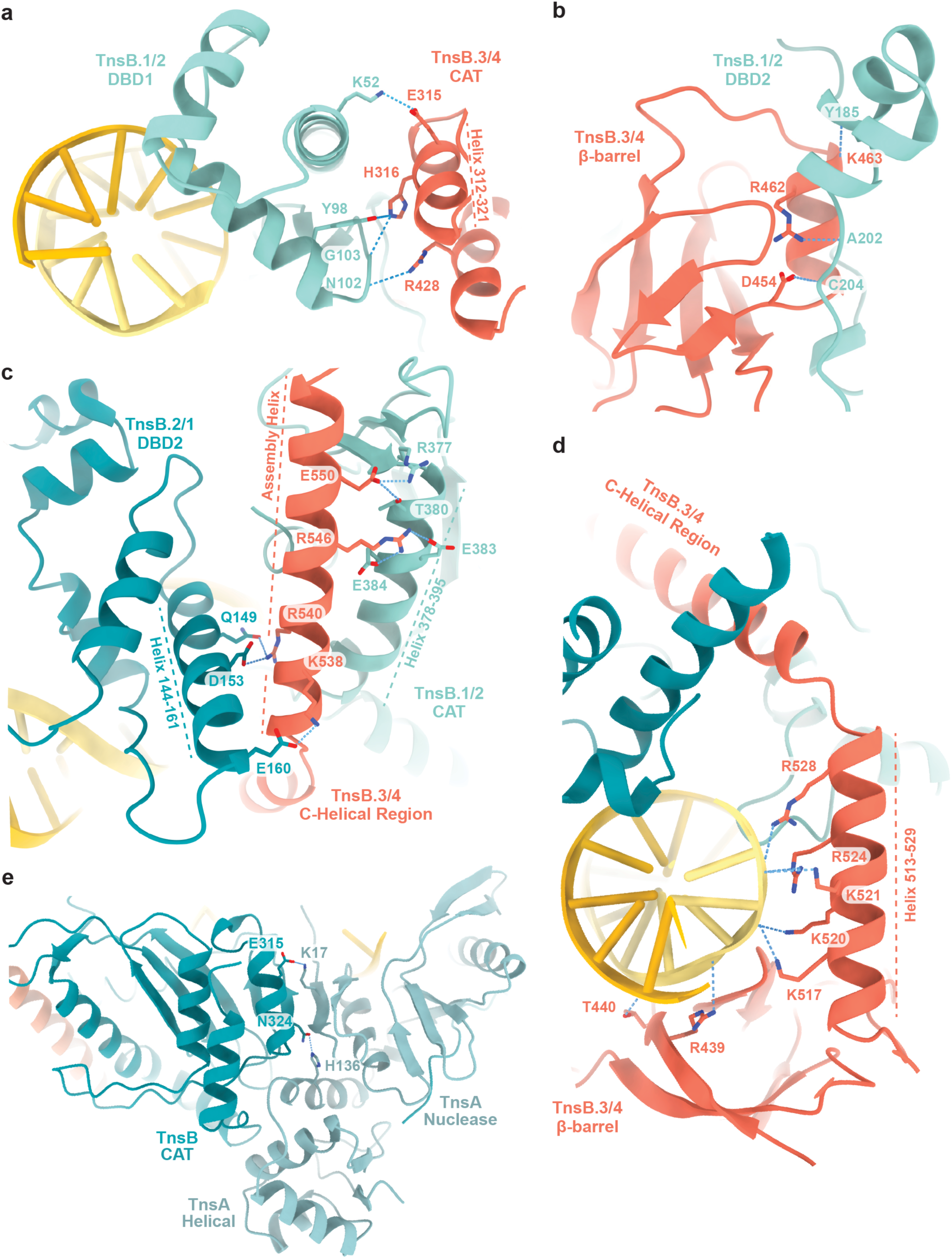
Interprotomer interfaces and TnsA-TnsB contacts. (**a**-**b**-**c**) Close-up views of interactions between TnsB protomers mediating tetramer formation. (**a**) Helix 312-321 in the CAT domain of each structural protomer packs against the non-DNA binding helix of DBD1 of each catalytic protomer. (**b**) The β-barrel of each structural protomer makes contacts with the DBD2 of each opposing catalytic protomer. (**c**) A C-terminal assembly helix in the structural protomer bridges the DBD2 of one catalytic protomer and the CAT of another, forming the main interprotomer interface. (**d**) The β-barrel and the first C-terminal helix in each structural protomer contact the DNA backbone, stabilizing the assembly helix and coupling DNA binding to oligomerization. (**e**) Detailed views of interactions between TnsA and TnsB. TnsA engages TnsB through contacts involving both its endonuclease and helical subdomains. Dashed lines represent interaction distances ≤ 3.5 Å.

The most extensive interprotomer interface is formed by the second helix (residues Ser533–Glu556^TnsB^) of the C-terminal helical region of TnsB.3 (or TnsB.4), which is sandwiched between the DBD2 of TnsB.2 (or TnsB.1) and the catalytic domain of TnsB.1 (or TnsB.2) (**Fig. 3c**, **Supplementary Fig. 6c**). The position of this helix, hereafter referred to as the “assembly helix”, is stabilized by the β-barrel domain and the first helix of the C-terminal region. These elements engage in nonspecific interactions with the deoxyribose-phosphate backbone through residues Arg439^TnsB^, Thr440^TnsB^, Lys517^TnsB^, Lys520^TnsB^, Lys521^TnsB^, Arg524^TnsB^, and Arg528^TnsB^ (**Fig. 3d**, **Supplementary Fig. 6d**), likely contributing to the structural coupling between transposon end binding and transposase oligomerization.

The structure further reveals how TnsA engages the catalytic TnsB protomer within the transpososome. The TnsA.1 (or TnsA.2) endonuclease domain interacts with TnsB.1 (or TnsB.2) by extending the five-stranded β-sheet of its RNase H-like fold via hydrogen bonding, resulting in a combined seven-stranded β-sheet, an interaction mode also observed in *Pmc*TnsAB^53^. A key contact involves a salt bridge between Lys17^TnsA^ and Glu315^TnsB^ in the CAT domain (**Fig. 3e**, **Supplementary Fig. 6e**). These residues are also conserved across other type I-F TnsB orthologs (**Supplementary Data 1, Supplementary Data 2**). In *Ec*Tn7, the interaction between TnsA and TnsB is mediated in part by TnsC, which forms extensive interfaces with both proteins and is required for transposase activity under physiological Mg^2+^ conditions^57,64^. The crystal structure of TnsA in complex with the C-terminal tail of TnsC reveals an interaction with a hydrophobic patch on the surface of TnsA, which enhances DNA binding and stimulates TnsAB-mediated excision^65^. In the type I-B2 *Pmc*CAST, the interaction between TnsA and TnsB is stabilized by the conserved Arg369-RxxxRD-Asp374 motif in the TnsC C-terminal tail, although the mode of interaction differs from that observed in *Ec*Tn7^53^. Notably, the TnsC C-tail sequence is not conserved in *Pse*TnsC or its orthologs, suggesting that TnsC is dispensable for functional TnsA-TnsB communication in type I-F CASTs. Consistent with this, addition of *Pse*TnsC had no effect on the efficiency of TnsAB-catalyzed transposon excision *in vitro* (**Supplementary Fig. 7a**-**c**).

The helical subdomain of TnsA also contacts the helical portion of the RNase H-like fold in TnsB via hydrogen bonding. A specific side-chain interaction is observed between His136^TnsA^ and Asn324^TnsB^ (**Fig. 3e**, **Supplementary Fig. 6e**). Finally, the N-terminal tail of TnsB further stabilizes the complex through a series of coordinated interactions: Phe4^TnsB^ is π-stacked between two consecutive histidines, His40^TnsA^ and His41^TnsA^; Glu10^TnsB^ forms a salt bridge with Arg98^TnsA^; and Glu7^TnsB^ forms a hydrogen bond with Thr119^TnsA^ (**Supplementary Fig. 7d**). These residues are overall conserved among type I-F orthologs (**Supplementary Data 1, Supplementary Data 2**) but not in type I-B2 *Pmc*TnsAB, indicating that inter-subunit contacts mediated by the TnsB N-terminus are a distinctive feature of *Pse*TnsAB and related type I-F systems.

### Transposon end binding by *Pse*TnsAB

Protein-DNA contacts are distributed throughout both catalytic and structural TnsB protomers, involving all major structural domains (**Fig. 4**, **Supplementary Fig. 8**). Multiple non-specific contacts with the DNA deoxyribose-phosphate backbone contribute to overall complex stabilization, whereas sequence-specific interactions are concentrated in the TnsB N-terminal domains, where both DBDs and the intervening linker participate in DNA recognition (**Fig. 4a**, **Supplementary Fig. 8a**-**c**). In DBD1, Arg89^TnsB^ and Arg93^TnsB^ form hydrogen bonds with dG-23 and dT-22, respectively. The linker connecting DBD1-DBD2 follows the minor groove and contributes further specificity through Met115^TnsB^ and Asn117^TnsB^, which contact dG17 and dT15, while Arg121^TnsB^ inserts between dG-13 and dT-14. In DBD2, Asn173^TnsB^ forms base-specific interactions with dT-10 and dG-9, extending the recognition interface.

**Fig. 4.**
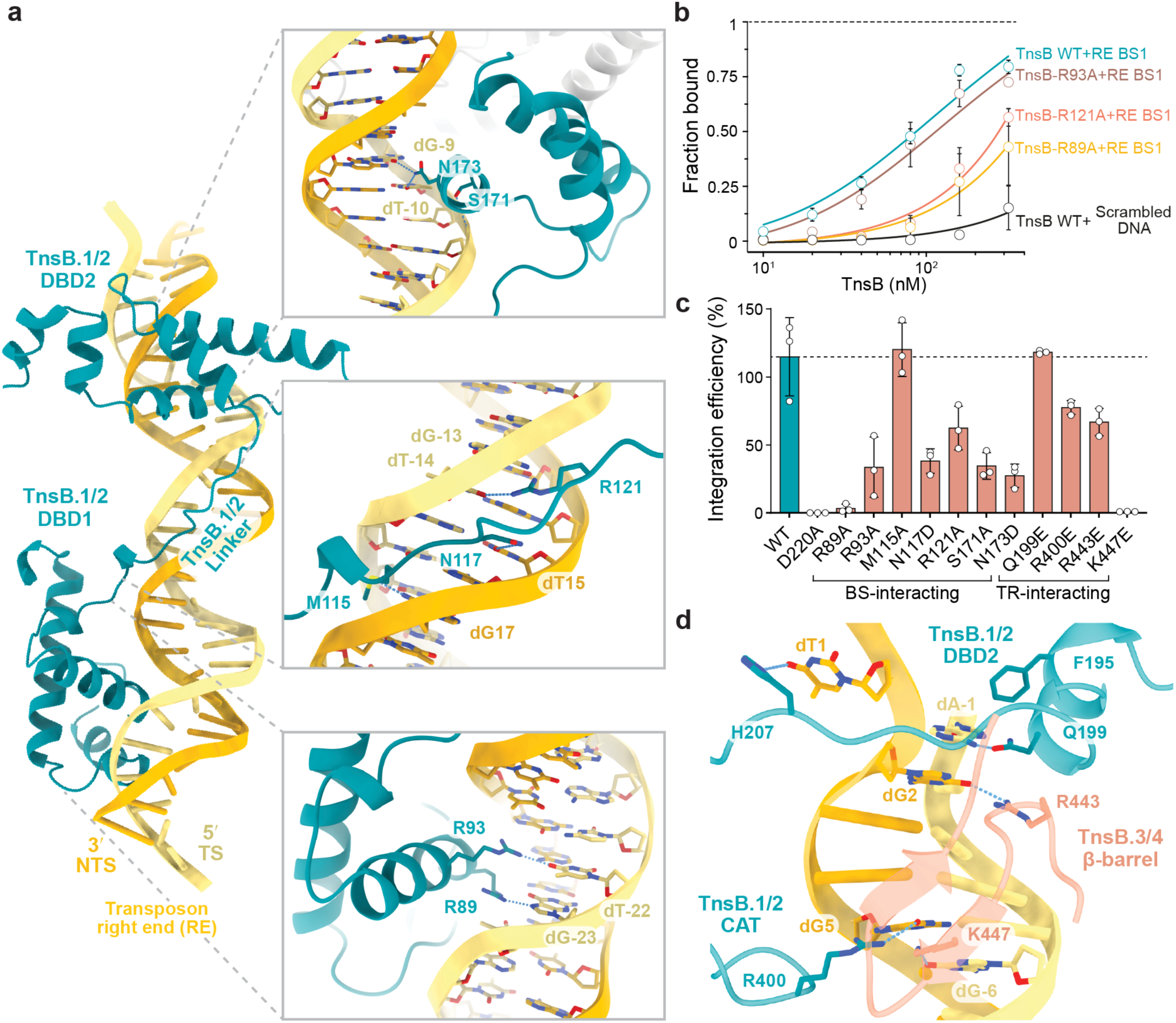
Transposon right end binding. (**a**) Detailed views of sequence-specific protein-right end (RE) DNA interactions mediated by the DBD2 (top), linker (middle) and DBD1 (bottom) of the TnsB catalytic protomers. Dashed lines represent atomic contact distances ≤ 3.5 Å. (**b**) Analysis of electromobility shift assay (EMSA) using fluorophore-labeled dsDNA. The DNAs were either RE BS1 DNA, containing the native BS1 (binding site 1) and the terminal repeat (TR) sequences of the transposon right end (RE), or a scrambled sequence, as indicated in **Supplementary Fig. 8d**. Reactions included either wild-type (WT) or mutant *Pse*TnsB proteins. Graph shows the fraction of bound DNA versus TnsB concentration (mean ± s.d., *n* = 3). Representative native PAGE gels for WT are shown in **Supplementary Fig. 8d**. (**c**) Site-specific integration activity in *E. coli* of *Pse*CAST systems containing structure-based mutations in DNA binding interface of TnsB, as determined by quantitative PCR (qPCR) analysis. (**d**) Detailed view of recognition of 8-bp TR, showing dT1 base flipping and sequence-specific contacts by catalytic protomer DBD2, linker, and CAT, reinforced by β-barrel contacts from the structural protomer.

To evaluate DNA binding specificity, we performed electrophoretic mobility shift assays (EMSAs) using *Pse*TnsB and fluorescently labeled RE DNA oligonucleotides containing BS1 and the 8-bp TR (**Fig. 4b**, **Supplementary Fig. 8d**). Clear shifts indicative of complex formation were observed only with the wild-type RE sequence and not with scrambled DNA, demonstrating sequence-specific recognition by *Pse*TnsB. To probe the importance of the interactions observed in the structure, we focused on arginine residues in the DBDs and the linker that contact conserved nucleotides in both right and left BSs (**Supplementary Fig. 2d**). Alanine substitutions of Arg89^TnsB^ and Arg121^TnsB^ markedly reduced DNA binding (**Fig. 4b, Supplementary Fig. 8f**), confirming their functional role in transposon end recognition. We subsequently tested the activity of a larger set of *Pse*TnsB variants containing mutations in DNA binding interface residues using transposition assays in *E. coli*. Except for the M115A variant, all substitutions substantially reduced *Pse*CAST activity *in vivo* (**Fig. 4c**).

In the structure, the 8-bp TR forms a continuous DNA duplex, but the terminal bases of nucleotides dA-1 (on TS) and dT1 (on NTS) are splayed and stabilized by specific interactions with the catalytic TnsB protomers (TnsB.1 and TnsB.2). This local unpairing is driven by a base-extrusion mechanism in which dT1 is flipped out of the duplex (**Fig. 4d**, **Supplementary Fig. 8c**) and base-stacks against His207^TnsB^, located in the linker connecting DBD2 and the CAT domain, while dA-1 is recognized by Gln199^TnsB^ and stacks against Phe195^TnsB^, which lies in the C-terminal helix of DBD2. Although the splayed terminal base pair and the TGT motif at the TR are conserved in type V-K *Sh*TnsB and type I-B2 *Pmc*TnsB, they are engaged in distinct ways (**Supplementary Fig. 8e**). Further downstream, a flexible loop within the CAT domain contributes to base-specific recognition, with Arg400^TnsB^ engaging in direct contact with dG5. In turn, the structural TnsB protomers (TnsB.3 and TnsB.4) engage the DNA through base-specific contacts, with Arg443^TnsB^ and Lys447^TnsB^ in the β-barrel domain interacting with dG2 and dG-6 (**Fig. 4d**, **Supplementary Fig. 8c**), positioning the DNA for cleavage in TnsB.1 and TnsB.2. To validate the structural observations, we generated mutant TnsAB variants containing alanine substitutions of Gln199, Arg443, and Lys447 and tested their activity in *in vitro* excision assays (**Supplementary Fig. 8f**-**g**). While the Q199A variant remained functional, the R443A and K447A substitutions led to markedly reduced activity. Corroborating these results, charge-reversal mutations of TR-interacting residues (R400E, R443E and K447E) substantially reduced or abolished *in vivo* transposition (**Fig. 4c**).

To further dissect the contribution of transposon end DNA sequences to transpososome assembly, we analyzed complex formation by SEC (**Supplementary Fig. 9a**) using RE DNAs identical to those used for cryo-EM, either with wild-type sequence or with scrambling mutations in BS2 or the TR (**Supplementary Fig. 9b**). Analytical SEC revealed two distinct nucleoprotein assemblies when using wild-type or BS2-mutated RE DNAs. In these cases, the two species were observed at comparable ratios. In contrast, TR-mutated DNAs predominantly yielded the lower-order assembly (**Supplementary Fig. 9c**). Mass photometry analysis confirmed these species as tetrameric TnsAB assembled on two RE DNA molecules and monomeric TnsAB bound to a single RE DNA molecule (**Supplementary Fig. 9d**). These experiments indicate that engagement of a second binding site at the right end is not required for tetramerization and end pairing, whereas an intact TR sequence is critical.

To investigate whether DNA binding interactions are conserved at the transposon left end, we next determined a structure of *Pse*TnsAB bound to a LE DNA mimicking the excision product, including BS1 and 17 base pairs between BS1 and BS2 (**Supplementary Fig. 10a**), at an overall resolution of 2.51 Å (**Supplementary Fig. 10b**-**f**, **Supplementary Table 1**). Consistent with our observations at the RE, a single TnsB-binding site supports PEC formation at the LE. The RE- and LE-bound complexes adopt highly similar architectures, superposing closely with minimal deviations (r.m.s.d. 0.332 Å over 607 pruned atom pairs). Clear density is observed for the TR-BS1 duplex, 4 bp of flanking DNA, and the 5′-phosphorylated AGA overhang on the NTS (**Supplementary Fig. 10a**). DNA binding residues adopt largely similar conformations in both structures (**Supplementary Fig. 10g**). Within the resolved region, the only sequence difference at a position contacted by TnsB occurs at position 15 of the NTS (dA in LE, dT in RE) (**Supplementary Fig. 10a**). Asn117^TnsB^ maintains base-specific interactions, engaging the N3 atom of dA15. Minor local differences are observed at the 5′ terminus of the NTS, where dT1 and the phosphate group of dG2 are shifted by ∼3.5 Å between the RE and LE structures. In the LE complex, this positions the dT1 base parallel to His207 ^TnsB^ and places the dG2 phosphate in proximity to Cys204^TnsB^ (**Supplementary Fig. 10g**).

Taken together, these results support a conserved structural framework for terminal binding site recognition at both transposon ends, and highlight the critical role of the TR sequence, together with the intertwined arrangement of structural and catalytic protomers, in mediating end synapsis and catalysis.

### Molecular activity of evolved *Pse*TnsAB

Building on our structural and biochemical reconstitution of the *Pse*TnsAB transposase, we next sought to elucidate the mechanistic basis of the enhanced activity of the evolved *Pse*TnsAB variant (evoTnsAB), generated via phage-assisted continuous evolution (PACE) for efficient genome integration in mammalian cells^27^. Mapping the mutations onto our structures allowed us to rationalize the potential contributions of two substitutions. F43S^TnsB^, situated in DBD1, may facilitate transposon end recognition, while P88T^TnsA^ located near the phosphate backbone of the cleaved NTS may stabilize the 3-nucleotide 5′ overhang (**Supplementary Fig. 11a**). The remaining substitutions are located within flexible or unresolved regions of the CAT, β-barrel, and C-terminal helices of TnsB, suggesting that they play roles in protein fold stabilization or interprotomer contacts. This is consistent with the AlphaFold3 models of the evoCAST TnsAB STC^27^ and PEC (**Supplementary Fig. 11b**), which closely match the experimental structure (r.m.s.d. 0.885 Å over 374 pruned atom pairs).

EMSA analysis revealed that evoTnsB does not exhibit a substantially increased binding affinity for RE DNA compared to the wild-type (**Supplementary Fig. 12a**-**b**). However, slowly migrating complexes, indicative of enhanced formation of higher-order nucleoprotein assemblies, appeared at lower protein-to-DNA ratios as compared to the wild-type *Pse*TnsAB (**Supplementary Fig. 12c**). To investigate this further, we performed mass photometry, which showed an increased propensity of evoTnsAB to oligomerize in both the apo state (31% dimers for evo versus 14% for wild-type) and DNA-bound state (28% complexes for evo versus 20% for wild-type). evoTnsB alone also exhibited an increased tendency to oligomerize (**Supplementary Fig. 12d**), suggesting that TnsB interprotomer interactions are enhanced in the evolved variant. Previous experiments in *E. coli* showed that evolved *Pse*CAST variants exhibit faster DNA insertion kinetics *in vivo*^27^. To test whether enhanced protein assembly influences transposon end cleavage, we carried out *in vitro* time-course excision assays at both 30 °C and 37 °C (**Fig. 5a**, **Supplementary Fig. 12e**-**f**-**g**), the latter approximating conditions used for genome editing in human cells. At 30 °C, evoTnsAB processed a greater proportion of supercoiled DNA than WT *Pse*TnsAB at later time points (60-240 min), whereas at 37 °C their supercoiled DNA processing activities were largely comparable (**Supplementary Fig. 12f**). Although the initial rates of plasmid linearization were similar at both temperatures, WT *Pse*TnsAB produced higher levels of the plasmid backbone product at the expense of the linear form at later time points (15-240 min), indicative of robust complete transposon excision. In contrast, evoTnsAB continued to accumulate the linear product with only minimal yield of fully excised product, consistent with predominantly single-end cleavage activity (**Fig. 5a**, **Supplementary Fig. 12e, g**). These findings suggest that laboratory evolution of *Pse*TnsAB introduced subtle changes in the higher-order assembly of the transposase that may favor nucleolytic processing at single transposon ends.

**Fig. 5.**
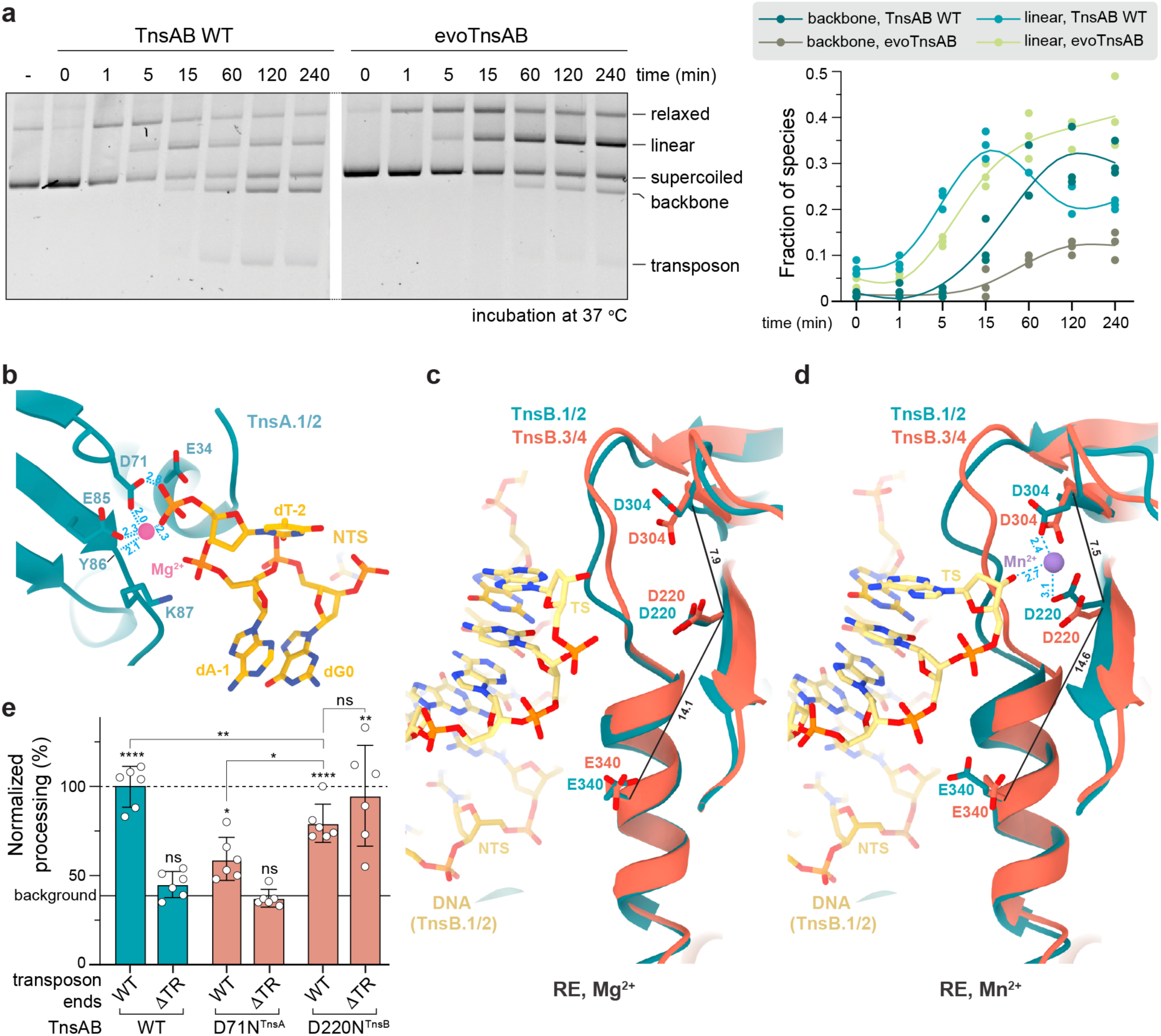
Transposon end cleavage. (**a**) *In vitro* time-course excision assays of wild-type (WT) and evolved (evo)*Pse*TnsAB variants at 37 °C. A representative agarose gel showing resolved DNA species (left) and associated densitometric quantification of plasmid backbone (∼2 kb-long excision product) and linearized plasmid (∼3 kb-long single-end cleaved product) (right) from three independent experiments are shown (see **Methods** for details). (**b**) Detailed view of the TnsA active site bound to transposon right end (RE) DNA in the presence of Mg^2+^. NTS, non-transferred strand. (**c-d**) Superposition of the active sites of the structural and catalytic TnsB protomers in RE-bound complexes assembled in the presence of Mg^2+^ (**c**) or Mn^2+^ (**d**). Inter-residue distances are shown as solid black lines. Coordination distances between catalytic residues and the divalent cation are indicated by blue dashed lines. Distances are given in Å. (**e**) *In vitro* excision assay with *Pse*TnsAB catalytic site mutants. Reactions included transposon ends in either wild-type (WT) configuration or lacking terminal repeats at both ends (ΔTR); -, no-protein control. Quantification from six independent experiments. Relative amounts of DNA species were determined by densitometry. Data show mean ± s.d. of all processed species normalized to the TnsAB WT on WT transposon ends. The solid horizontal line indicates background levels of processed plasmid in the no-protein control and using WT transposon ends. Statistical significance was assessed using unpaired two-tailed *t*-tests. P-values above each graph indicate comparisons with the corresponding negative control. P-values: *p < 0.05; **p < 0.01; ****p < 0.0001; ns, not significant (see **Methods** for details). Representative agarose gels are shown in **Supplementary Fig. 14h**.

### Transposon end cleavage catalysis

Finally, the *Pse*TnsAB structures visualize how the nuclease domains of *Pse*TnsA and *Pse*TnsB and their catalytic sites are positioned on the cleaved transposon termini (**Fig. 5b-c**, **Supplementary Fig. 13**). The active site cleft of TnsA accommodates the 3-nucleotide 5′ overhang (TAG) of the NTS, with the phosphate group precisely coordinated by the catalytic triad (**Fig. 5b**), consistent with the staggered cleavage pattern observed *in vitro*. The bases of NTS nucleotides dA-1 and dG0 form a base stack together with the aromatic side chain of Tyr23^TnsB^ from the N-terminal tail of TnsB (**Supplementary Fig. 13a**) as they insert into the active site cleft of TnsA. TnsA is captured with the 5′-terminal phosphate group of the pre-cleaved NTS positioned within its active site, adjacent to the catalytic triad Glu34^TnsA^, Asp71^TnsA^, and Lys87^TnsA^ (**Fig. 5b**). The cryo-EM maps feature density consistent with a single magnesium ion coordinated by Asp71^TnsA^, Glu85^TnsA^ and the backbone carbonyl of Tyr86^TnsA^ (**Supplementary Fig. 13b**-**c**). Lys87^TnsA^ lies near the scissile phosphate, likely contributing to DNA positioning for catalysis and stabilizing the non-bridging oxygen in the transition state (**Fig. 5b**, **Supplementary Fig. 13b**-**c**). The overall active site geometry and DNA backbone alignment indicate that the captured state corresponds to a catalytically competent conformation, despite the absence of one of the divalent cations.

TnsB catalyzes both 3′ end cleavage and subsequent strand transfer into the target DNA via its DDE catalytic triad (Asp220^TnsB^, Asp304^TnsB^, Glu340^TnsB^)^6^, which coordinates two divalent metal cations to activate a water molecule for nucleophilic attack on the DNA backbone^55^. In both RE and LE PEC structures determined in the presence of Mg^2+^, the CAT domains of all four *Pse*TnsB protomers are overall structurally very similar (r.m.s.d. 0.671 Å over 173 Cα for RE complex), and the active site residues of both catalytic and structural TnsB subunits adopt an inactive conformation (**Fig. 5c**, **Supplementary Fig. 13d**-**e**). The distances between Asp220^TnsB^ and Asp304^TnsB^ and the 3′-hydroxyl of the cleaved TS indicate they are not poised for catalysis. Furthermore, Glu340^TnsB^ is positioned at a distance of 14.071 Å (Cα distance) from Asp220^TnsB^. As a result, the TnsB catalytic triad is not properly oriented to enable the binding of the Mg^+2^ ion. To further investigate the organization of the TnsB active site, we determined a structure of an RE-assembled PEC in the presence of Mn^2+^ at an overall resolution of 2.86 Å (**Fig. 5d**, **Supplementary Fig. 14a**-**e**, **Supplementary Table 1**). Additional density is consistent with a single Mn^2+^ ion coordinated by Asp220^TnsB^ and Asp304^TnsB^ of the catalytic protomers (**Fig. 5d**, **Supplementary Fig. 13f**). The catalytic triad adopts a similar conformation to that observed in the Mg^2+^-bound structures and remains incompatible with catalysis, with Glu340^TnsB^ positioned distally. These observations suggest that Mn^2+^ may bind with higher affinity than Mg^2+^, potentially contributing to enhanced *in vitro* activity, although further studies are required to determine how Mn^2+^ coordination promotes the formation of a catalytically competent active site conformation under these conditions.

Notably, the relative disposition of the TnsA and TnsB active sites in the complex is incompatible with the canonical B-form geometry of duplex DNA and would not permit simultaneous positioning of both scissile phosphates in the active sites of TnsA and TnsB without steric clashes (**Supplementary Fig. 14f**). This indicates that concurrent binding of the TS and NTS in their respective catalytic clefts requires structural remodeling of the bound transposon end DNA, likely driven by local strand separation induced by splaying of the terminal base pair in the TR by interactions with TnsB. As a result, NTS and TS cleavage events are likely interdependent and spatially coordinated between TnsA and TnsB. To further investigate this, we performed *in vitro* excision assays with *Pse*TnsAB variants carrying active site mutations in either subunit (**Fig. 5e**, **Supplementary Fig. 14g**-**h**). Catalytic inactivation of either TnsB (D220N^TnsB^) or TnsA (D71N^TnsA^) resulted in plasmid nicking above background levels, with the TnsB-inactive mutant exhibiting overall higher processing activity than the TnsA-inactive mutant. In the D220N^TnsB^ mutant, TnsA mediated additional unspecific nicking when a TR-mutated plasmid was used, indicating that TnsA may exhibit limited sequence specificity. These findings suggest that the efficiency and specificity of TS and NTS nicking by TnsB and TnsA are functionally interdependent. Overall, our data suggest that TnsA and TnsB activities are not only complementary but also coordinated within the PEC, with cleavage events likely occurring sequentially.

## Discussion

Our structural and functional analyses provide mechanistic insight into the type I-F CAST transposition pathway (**Fig. 6**) and identify molecular features that may underlie the robustness, fidelity, and specificity of *Pse*CAST as a programmable genome engineering tool.

**Fig. 6.**
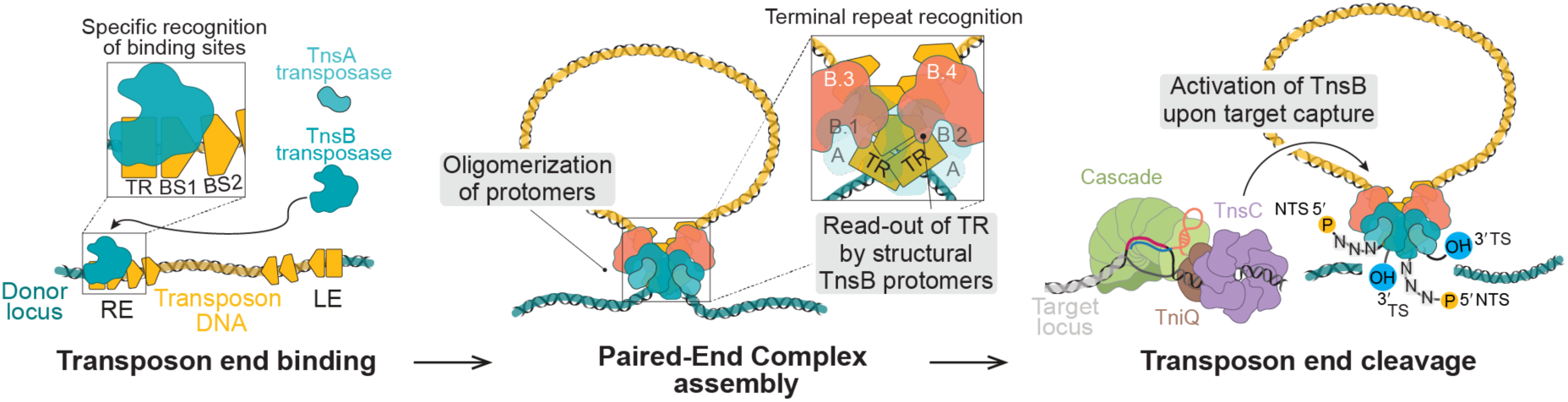
Mechanism of transposition in type I-F CASTs. Proposed mechanistic model for cut-and-paste transposition catalyzed by type I-F CRISPR-associated transposases. The TnsB transposase specifically recognizes its cognate binding sites within transposon right and left ends (RE and LE). DNA binding by TnsB induces oligomerization of transposase protomers, promoting the formation of the TnsAB paired-end complex (PEC). PEC assembly is further facilitated by recognition of terminal repeat (TR) sequences by the structural protomers of TnsB. TnsA cleaves the non-transferred strand (NTS) on both transposon ends 3 nucleotides outside the TR sequence. Following activation by TnsC assembled at the target site, TnsB cleaves the transferred strand (TS) at the transposon boundary, providing free 3′-OH used in the strand transfer reactions, ultimately leading to transposon integration at the target locus.

A key finding is that *Pse*TnsB recognizes transposon ends with high specificity, detectable even at single binding sites within both the right and left ends. This contrasts with type V-K^41,42,66^ and *Ec*Tn7 TnsB^54^, which bind promiscuously with modest sequence preference and few base-specific protein-DNA contacts, and with type I-B2 TnsB, which achieves high affinity only when two binding sites are present^53^. These differences are likely reflected in the distinct organization of the two final binding sites at the target-proximal end - non-overlapping in type I-F (RE) but overlapping in Tn7 (RE) and type I-B2 (LE)^31^, where such architecture may amplify weak specificity. Variation in DNA-contacting residues among type I-F TnsB orthologs likely explains their orthogonality, enabling closely related systems to coexist without cross-reactivity and supporting independent insertions without target immunity^36^, i.e. protection of nearby DNA from re-insertion mediated by transposase-dependent end binding. By contrast, *Ec*Tn7 is thought to rely on binding across extended DNA regions due to its weak sequence preference, conferring broad target immunity^67^ and causing strong inhibition upon TnsB overexpression (overproduction inhibition)^54,68^. In *Pse*CAST, however, transposition activity increases with higher transposase levels^26^, likely reflecting its highly specific recognition of transposon ends.

Our data indicate that the *Pse*CAST transposase assembles through protomer oligomerization upon DNA binding, similar to *Ec*Tn7^54^ and type V-K system^41,42^, and does not require TnsC, in contrast to type I-B2^53^. This protomer-driven pathway may also explain the absence of overproduction inhibition^26^, in contrast to dimeric transposases that hinder end synapsis by simultaneous occupation of both ends at high protein concentrations^69^. Although determined using minimal RE- and LE-derived DNAs and thus not capturing the fully physiological PEC configuration, the close agreement between symmetrized RE- and LE-bound structures indicates conserved structural features likely maintained in LE-RE complexes, stabilized by extensive TnsB-TnsB and TnsB-DNA contacts that support the intertwined *trans* architecture adopted upon end pairing. In this arrangement, the catalytic and structural TnsB protomers couple tetramerization to DNA binding and cleavage, preventing aberrant single-end cleavage. Notably, our data establish that terminal repeat (TR) recognition is essential for this process, as structural protomers engage the repeats to position catalytic protomers for cleavage. Disruption of TR binding, either by single point mutations in TnsB or by DNA sequence scrambling, substantially impairs excision and transposase tetramerization. These findings indicate that productive end pairing is likely initiated at the outermost terminal repeats, whose sequence serves as a trigger for PEC assembly, thereby imposing a fidelity checkpoint.

An additional feature we observe in *Pse*TnsAB is the apparent interdependency between TnsA and TnsB, which may extend to other type I-F systems. *In vitro* excision assays show that the cleavage pattern is dictated by the relative positions of their catalytic domains, similar to *Ec*Tn7^58^. Moreover, nicking by TnsA occurs more robustly, whereas TnsB nicking is comparatively weak, consistent with a model in which TnsB requires activation. Our structural data support a sequential cleavage model in which NTS cleavage may precede TS cleavage, since the geometry of pre-cleaved DNA prevents simultaneous positioning of both scissile phosphates and requires remodeling between events. Cutting the TS last would ensure that its 3′-terminal hydroxyl group is aligned within the TnsB active site for strand transfer. It is conceivable that productive excision under physiological conditions requires engagement of donor DNA by the transposase and binding to target DNA upon recruitment by ATP-bound TnsC oligomers. Such a mechanism may help ensure that full excision occurs only after target capture, while single-strand nicks remain innocuous because they are efficiently repaired *in vivo*.

*In vivo* data from type I-F CASTs align with this interdependency model. Transposase mutants with a disrupted TnsA catalytic pocket can still transpose via TnsB, but the pathway is converted into copy-and-paste with only ∼50% efficiency^32^. By contrast, type I-B2 CASTs are intrinsically prone to cointegrate formation^19,53^, indicating weaker coordination between NTS and TS cleavage. Collectively, these findings suggest that tight functional coupling between TnsA and TnsB is a defining property of type I-F CASTs and underlies their high-fidelity cut-and-paste mechanism.

Finally, our analysis of the evolved *Pse*TnsAB variant shows that laboratory-selected mutations result in an increased propensity for higher-order assembly, consistent with strengthened TnsB inter-subunit interactions. Although recognition of single binding sites was not improved *in vitro*, a threefold increase in the engagement of complete transposon right ends has been reported *in vivo*^27^, likely explained by cooperative binding. Yet, enhanced oligomerization does not translate into more efficient PEC formation as evoTnsAB preferentially accumulates single-end products *in vitro*. Unlike in natural transposons, this misregulation might not be relevant in a genome editing context, where excision proceeds from donor plasmids and can be circumvented by using linearized donor DNA^27^. Of note, none of the substitutions present in evoTnsAB correspond to TnsC-bypassing mutations identified in *Ec*Tn7^64^, suggesting that the observed hyperactivity arises through a distinct mechanism. Consistently, evoTnsAB-mediated off-target insertions in human cells remain dependent on TnsC^27^. Overall, our experiments suggest that the laboratory-evolved mutations enhance integration rather than excision, consistent with the selection pipeline that favored variants with elevated insertion rates.

Taken together, our results define key molecular principles of type I-F CAST transposition, where sequence-specific recognition, transpososome assembly, and regulated nuclease activity ensure a precise cut-and-paste mechanism (**Fig. 6**). These insights provide rationale for the unique robustness and fidelity of *Pse*CAST and establish a mechanistic foundation for engineering CRISPR-associated transposases as versatile genetic tools in biotechnology and synthetic biology.

## Acknowledgments

We thank the Next Generation Sequencing Facility at Vienna BioCenter Core Facilities (VBCF) for performing Illumina sequencing. We are grateful to the UZH Center for Microscopy and Image Analysis for providing electron microscope access and technical support. We further thank M. Klaußner for help with assay optimization, R. H. Budiman for assistance with mass photometry measurements, S. Falk for assistance with SEC-MALS measurements, T.M. Smith for laboratory support, and M. Schmitz for initial experiments at the start of the project. We also thank the members of the laboratories of Sebastian Falk and Thomas Leonard for helpful discussions about the project. This work was supported by the Horizon Europe European Research Council (ERC) Starting Grant BROADCAST (grant number 101115765) awarded to I.Q, ERC Consolidator Grant CRISPR2.0 (number 820152) and Swiss National Science Foundation Project Grant (number 320030-228089) awarded to M.J. I.Q. received funding from The Branco Weiss Fellowship – Society in Science, administered by ETH Zürich, as well as a Vallee Scholar Award from the Vallee Foundation. M.J. received funding from the Swiss National Center for Competence in Research (NCCR) RNA & Disease. S.H.S. was supported by NIH grants DP2HG011650, RM1HG009490, R01EB027793, and R01EB031935; a Pew Biomedical Scholarship, an Irma T. Hirschl Career Scientist Award, the Howard Hughes Medical Institute Investigator Program, and a generous startup package from the Columbia University Irving Medical Center Dean’s Office and the Vagelos Precision Medicine Fund. M.W. and I.C.H. are members of the Vienna BioCenter PhD Program, a Doctoral School of the University of Vienna and the Medical University of Vienna. G.F. and S.O. are members of the Biomolecular Structure and Mechanism doctoral program of the Life Science Zurich Graduate School. M.W. was supported by a DOC Fellowship from the Austrian Academy of Sciences.

## Author contributions

I.Q. and M.J. conceived the study. I.Q., M.J., M.W., G.F. and S.O. developed the experimental design. M.W. performed mass photometry experiments, *in vitro* excision assays, and determined the TnsAB cleavage positions by preliminary Sanger sequencing. G.F. and S.O. performed cryo-EM sample preparation, data collection and processing, and built the atomic models of the *Pse*TnsAB PEC. M.W. and I.C.H. performed protein purification and SEC-MALS measurements. G.D.L. designed and performed *in vivo* CAST transposition assays, with supervision from S.H.S.. J.K. carried out DNA binding experiments by EMSA. T.S. conducted next-generation sequencing experiments and wrote code to analyze the data. All authors contributed to data analysis. I.Q. and M.J. wrote the manuscript with input from all authors. M.W., G.F. and S.O. prepared figures. The manuscript was reviewed and approved by all authors.

## Materials & Correspondence

Correspondence and requests for materials should be addressed to Irma Querques.

## Methods

### Bacterial strains

For cloning purposes, *Escherichia coli* (*E. coli*) DH10B, OmniMAX and Mach1 T1 Phage-Resistant cells (ThermoFisher Scientific) were grown at 37 °C. For recombinant protein expression, *E. coli* BL21 Rosetta2 (DE3) cells (Novagen) were grown in Lysogeny Broth (LB) medium (Roth) at 37 °C until OD_600_ of 0.6 was reached, at which point they were induced with isopropyl β-D-1-thiogalactopyranoside (IPTG) for 18 h at 18 °C.

### DNA constructs

Plasmids containing human codon-optimized DNA sequences of TnsABf wild type (henceforth referred to as TnsAB WT), TnsC WT and evolved TnsAB (evoTnsAB) were obtained from Addgene (#200901, #200900 and #234729, respectively) and used to amplify the desired gene fragments with polymerase chain reaction (PCR) following the manufacturer’s instructions for Phusion polymerase (ThermoFisher Scientific). This allowed subsequent cloning into 1B_LIC (Addgene #29653; TnsC only) or 1C_LIC (Addgene #29654) vectors for protein expression using ligation-independent cloning (LIC)^70^, resulting in constructs carrying an N-terminal hexahistidine (His6) tag or a hexahistidine-maltose binding protein (His6-MBP) tag, each followed by a tobacco etch virus (TEV) protease cleavage site. The resulting plasmid pTnsAB_initial was additionally subjected to site-directed mutagenesis^71^ using the Q5 polymerase (NEB) protocol to remove a recombination hotspot, yielding the pTnsAB vector. Additional point mutations in TnsAB and TnsB sequences were introduced by site-directed mutagenesis with Q5 polymerase (NEB), using pTnsAB or pTnsB as templates.

*Pseudoalteromonas* Tn*7016* donor plasmid (pDonor_WT) containing minimal native transposon ends was obtained from Addgene (#200905) and used to generate pDonor plasmids containing mutated transposon ends by PCR using the Phusion polymerase and via Gibson assembly^72^. All constructs were cloned using either DH10B or OmniMAX *E. coli* strains (ThermoFisher Scientific). Plasmids were purified using the GeneJET Plasmid Miniprep kit (ThermoFisher Scientific) or MiniPEx kit (Vienna BioCenter Molecular Tool Development shop) and verified by Sanger sequencing (MicroSynth AG). pDonor plasmids were propagated in *E. coli* Mach1 cells (ThermoFisher Scientific), isolated using the MiniPEx kit, and subjected to whole-plasmid sequencing (Eurofins Genomics) before final purification by ethanol precipitation for use in *in vitro* excision assays. Resulting DNA pellets were resuspended in ultrapure water at a concentration of 200 ng µL^-1^. All primer and oligonucleotide sequences used for cloning, and the relevant plasmid sequences are provided in **Supplementary Table 2**.

### Protein expression and purification

For biochemical and biophysical characterization of TnsAB WT, TnsB WT and their mutants, constructs carrying N-terminal His6-MBP were expressed in *E. coli* BL21 Rosetta2 (DE3) cells (Novagen). Cell cultures were grown in LB medium (Roth) at 37 °C, shaking at 150 rpm until OD_600_ of 0.6 was reached, and protein expression was induced with 0.4 mM IPTG and continued for 18 h at 18 °C. Cell pellets were resuspended in 60 mL Ni-wash buffer (20 mM Tris-HCl pH 7.0, 500 mM NaCl, 5 mM imidazole, 5% [v/v] glycerol) supplemented with 1 μg mL^-1^ pepstatin, 200 μg ml^-1^ AEBSF, and 1 μg ml^-1^ DNAse. Cells were lysed by ultrasonication (using a Branson sonifier W-450D equipped with 102C (CE) tip, 1 s on/2 s off cycle, 30% amplitude, 24 min total) and centrifuged at 36 000g for 75 min at 4 °C (Hitachi CR22N centrifuge with R20A2 rotor) to remove cell debris. Cleared lysates were filtered through 0.45 µm pore size and applied onto one (all mutant constructs) or two connected in tandem (TnsAB WT and TnsB WT) HisTrap HP 5 mL column(s) (Cytiva) pre-equilibrated with Ni-wash buffer using an ÄKTA Pure system (Cytiva). Columns were washed using Ni-wash buffer, and bound proteins were eluted using an imidazole gradient prepared by mixing Ni-wash buffer with Ni-elution buffer (20 mM Tris-HCl, pH 7.0, 500 mM NaCl, 500 mM imidazole, 5% [v/v] glycerol). Eluted fractions containing the protein of interest were pooled and dialysed overnight at 4 °C using 12-14 kDa molecular weight cut-off (MWCO) Spectra/Por tubing (Roth) against 2 L dialysis buffer, containing 20 mM Tris-HCl pH 7.0, 200 mM NaCl, 5% glycerol [v/v], 1 mM DTT for the TnsAB WT and TnsB WT proteins, or 20 mM Tris-HCl pH 7.0, 500 mM NaCl, 5% glycerol [v/v], 1 mM DTT for the other TnsAB and TnsB constructs. Dialysis was performed in the presence of excess His6-tagged TEV protease to remove the His6-MBP tag. Soluble fractions of dialysed proteins were applied to HiTrap Heparin HP 5 mL columns (Cytiva): one column for TnsAB-D71N^TnsA^, TnsAB-K447A^TnsB^, TnsB-R93A, TnsB-R121A, evoTnsAB, and evoTnsB; two tandem-connected columns for TnsAB-D220N^TnsB^, TnsAB-Q199A^TnsB^, TnsAB-R443A^TnsB^, TnsB WT, and TnsB-R89A; and three tandem-connected columns for TnsAB WT. Columns were pre-equilibrated with buffer A, containing 20 mM Tris-HCl pH 7.0, 200 mM NaCl, 5% glycerol [v/v], 1 mM DTT for TnsAB WT and TnsB WT, or 20 mM Tris-HCl pH 7.0, 500 mM NaCl, 5% glycerol [v/v], 1 mM DTT for the other constructs. Bound proteins were eluted with a linear gradient of buffer B (20 mM Tris-HCl pH 7.0, 1 M NaCl, 5% glycerol [v/v], 1 mM DTT). Pooled fractions were concentrated using 30 000 MWCO centrifugal filters (Merck Millipore) and further purified by size exclusion chromatography (SEC) on a HiLoad Superdex 200 16/600 column (Cytiva) pre-equilibrated with SEC buffer (20 mM Tris-HCl pH 7.0, 500 mM NaCl, 1 mM DTT). Fractions corresponding to monomeric and dimeric TnsAB or TnsB were pooled separately, concentrated to 2.5-18.5 mg mL^-1^ using 30 000 MWCO centrifugal filters, flash-frozen in liquid nitrogen, and stored at -80 °C.

For cryo-EM experiments, TnsAB WT was expressed and purified essentially as described for the His6-MBP-tagged TnsAB mutants, with the following modifications. Sonication was performed using a Bandelin Sonopuls HD 3200 equipped with a VS 70 T probe (1 s on / 2 s off, 30% amplitude, 15 min, 4 °C). The lysate was clarified at 40 000g for 60 min without post-spin filtration. Throughout the purification the buffer pH was maintained at 7.5 rather than 7.0. The Ni-affinity step used a stepwise elution with 5%, 20%, and 100% of 500 mM imidazole, and the 20% and 100% fractions were pooled for subsequent steps. As in the mutants workflow, the heparin step employed two HiTrap Heparin HP 5 mL columns (Cytiva) connected in tandem, with elution by a linear gradient to 1 M NaCl over 150 mL. Size-exclusion chromatography was performed on a Superdex 200 Increase 10/300 GL (Cytiva).

Expression, lysis, and first Ni-NTA affinity purification step of the TnsC construct, tagged N-terminally with His6 followed by a TEV protease site, followed procedure analogous to that described for TnsAB and TnsB constructs, using two tandem-connected HisTrap HP 5 mL columns (Cytiva). The only differences were that both Ni-wash and Ni-elution buffers were adjusted to pH 7.5. Pooled fractions containing TnsC were dialysed overnight at 4 °C using 12-14 kDa MWCO Spectra/Por tubing (Roth) against 2 L dialysis buffer (20 mM Tris-HCl pH 7.5, 500 mM NaCl, 5% glycerol [v/v], 4 mM β-mercaptoethanol) in the presence of excess His6-tagged TEV protease to remove the His6 tag. The soluble fraction of the dialysed protein was applied onto two tandem-connected HisTrap HP 5 mL columns (Cytiva) pre-equilibrated with Reverse-Ni-wash buffer (20 mM Tris-HCl pH 7.5, 500 mM NaCl, 5% glycerol [v/v]) and subsequently eluted with Ni-elution buffer (composition as above, adjusted to pH 7.5). Eluted fractions were pooled, concentrated using 10 000 MWCO centrifugal filters (Merck Millipore), and applied onto a HiLoad Superdex 200 16/600 column (Cytiva) pre-equilibrated with SEC buffer (20 mM Tris-HCl pH 7.5, 500 mM NaCl, 1 mM DTT). Fractions corresponding to monomeric TnsC were pooled, concentrated to 4 mg mL^-1^ using 10 000 MWCO centrifugal filters, flash-frozen in liquid nitrogen, and stored at -80 °C. Protein samples were analyzed by SDS-PAGE using 4-20% gradient polyacrylamide gels (Bio-Rad) stained with Quick Coomassie Stain (Protein Ark).

### DNA oligonucleotides

Double-stranded DNA (dsDNA) constructs used in electrophoretic mobility shift assay (EMSA), mass photometry experiments, as well as analytical size exclusion chromatography (SEC) experiment shown in **Supplementary Fig. 9a** were prepared by annealing equimolar amounts of two complementary oligonucleotides (IDT) resuspended in TE50 buffer (10 mM Tris-HCl pH 8.0, 1 mM EDTA, 50 mM NaCl). For constructs used in all EMSA experiments and mass photometry experiments shown in **Supplementary Fig. 3b and 12d**, DNA was denatured at 98 °C for 3.5 min and gradually cooled to 20 °C (0.2 °C per 90 s) in a thermocycler. For constructs used in analytical (SEC) and mass photometry experiments shown in **Supplementary Fig. 9a and 9d**, DNA was denatured at 98 °C for 3.5 min and gradually cooled to 20 °C (0.2 °C per 10 s) in a thermocycler. The DNA duplexes used in analytical SEC experiment shown in **Supplementary Fig. 3a**, as well as in cryo-EM sample preparation (o577/o578 or oGF254/oGF255) were prepared by mixing complementary 100 µM oligonucleotides in ultrapure water (20 µL total) in a 1:1 ratio. The mixture was annealed at 95 °C for 5 min, cooled to 4 °C over 1 h, and diluted to 10 µM in ultrapure water. dsDNA constructs were stored at -20 °C. Oligonucleotide sequences are provided in **Supplementary Table 2**.

### Electrophoretic mobility shift assay

DNA binding assays were conducted using 40 nM dsDNA oligonucleotides obtained by mixing 20 nM of 5′-Cy5-labeled dsDNA and 20 nM of corresponding unlabelled dsDNA. Binding reactions (20 µL) contained TnsB at concentrations ranging from 10 to 320 nM in a reaction buffer containing 20 mM HEPES-NaOH pH 7.0, 5 mM Tris-HCl pH 7.0, 400 mM NaCl, 1% glycerol [v/v], 50 µg mL^-1^ BSA, 1.25 mM DTT (final concentrations). Reactions were incubated on ice for 30 min before being mixed with orange G loading dye (0.02% [w/v] orange G, 5% [v/v] glycerol). Protein-DNA complexes were resolved at 4 °C on 5% native 0.5x TBE polyacrylamide gels, pre-equilibrated by running at 200 V for 60 min. Upon sample loading, gels were run at 90 V for 70 min, followed by 75 V for 10 min. Gels were imaged in an Amersham Typhoon imager (GE Healthcare) using Cy5 fluorescence readout.

Quantification analysis was performed in Fiji^73^ by measuring absolute densities of bands corresponding to three species: free DNA, TnsB-DNA complex I, and TnsB-DNA complex II, and subsequently normalizing them between replicates to the density of free DNA of the no-protein control. Densitometry data were analysed in GraphPad Prism version 10.2.0 (GraphPad Software, Boston, Massachusetts USA, www.graphpad.com). Fraction bound was calculated as:

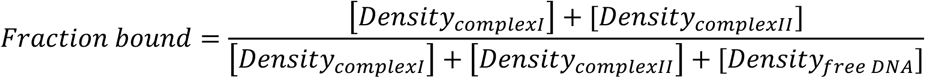

Sigmoidal binding curves were obtained using a one-site saturation binding model, with B_max_ imposed to 1.

### SEC-MALS analysis

The molecular mass of TnsAB WT was determined by size-exclusion chromatography coupled with multi-angle light scattering (SEC-MALS) at room temperature. Purified protein at a concentration of 2 mg mL^-1^ was loaded at 0.5 mL min^-1^ onto a Superdex 200 10/300 column (Cytiva) connected to a 1290 Infinity HPLC system (Agilent Technologies) equilibrated with buffer containing 20 mM HEPES pH 7.0, 400 mM NaCl, 5 mM CaCl_2_ and 1 mM DTT as mobile phase. Eluting particles passed through a miniDawn Treos II detector (Wyatt Technologies; laser emitting at 690 nm) and an RI-501 refractive index detector (Shodex). Data were analysed using the ASTRA 8 software package (Wyatt Technologies).

### Analytical size exclusion chromatography

For experiments shown in **Supplementary Fig. 3a**, TnsAB WT was diluted to 50 µM in 20 mM Tris-HCl pH 7.5, 500 mM NaCl, and 1 mM DTT. On ice, high-salt buffer (1050 mM KCl, 10 mM HEPES-KOH pH 7.5, 10 mM MgCl_2_, 1 mM DTT) was added first (11.4 µL for 80 µL final total volume), followed by either DNA (o577/o578, prepared as described above; sequences provided in **Supplementary Table 2**) from the 10 µM stock (to 2 µM final) or ultrapure water for DNA-minus controls, and then either TnsAB from the 50 µM stock (to 8 µM final) or an equivalent volume of its stock buffer for protein-minus controls. Mixtures were brought to 80 µL by slow addition of dilution buffer (28 mM HEPES-KOH pH 7.5, 18 mM MgCl_2_, 1.5 mM DTT, 140 mM KCl), with gentle mixing throughout. Samples were incubated at 4 °C for 30 min, centrifuged at 4 °C, 14 000g for 10 min, and the supernatant was transferred to a pre-chilled tube. For SEC, 75 µL of each sample (TnsAB apo, PEC DNA, and TnsAB+PEC DNA) was manually injected onto a Superdex 200 5/150 GL column (Cytiva) connected to an ÄKTA Micro system and eluted at 0.15 mL min^-1^ in SEC buffer (20 mM HEPES-KOH pH 7.5, 10 mM MgCl_2_, 1 mM DTT, 300 mM KCl). Complex formation and oligomerization were evaluated by comparing elution profiles of TnsAB in the absence and presence of the DNA.

Analytical SEC experiments shown in **Supplementary Fig. 9a** were performed using the following dsDNA oligonucletides: WT (o577/o578), BS2* (o808/o809) or TR* (o806/o807), prepared as described above (sequences provided in **Supplementary Table 2**). Samples were prepared in 25 µL reaction volumes on ice, and reactions contained 12.5 µL of 50 µM TnsAB WT dilution, 3.57 µL high salt buffer (25.3 mM HEPES-NaOH pH 7.0, 410 mM NaCl, 36.4 mM MgCl_2_, 0.7 mM DTT), and 4.2 µL of the respective dsDNA (at 25 µM), and were slowly diluted with 4.73 µL of low salt buffer (15.86 mM HEPES-NaOH pH 7.0, 26.43 mM MgCl_2_) to produce assembly buffer at a final composition of 8.3 mM HEPES-NaOH pH 7.0, 10 mM Tris-HCl pH 7.0, 317 mM NaCl, 10.2 mM MgCl_2_, 0.6 mM DTT, containing 25 µM of respective protein and 4.2 µM DNA oligonucleotides. Apo-protein reaction was assembled in an analogous way, using TE50 buffer instead of the DNA dilution, while DNA only reaction contained SEC buffer instead of the protein dilution. Reactions were incubated on ice for 30 min. For SEC, 25 µL of each sample was manually injected onto a Superdex 200 Increase 3.2/300 column (Cytiva) connected to an ÄKTA Micro system and eluted at 0.05 mL min^-1^ in SEC buffer (20 mM HEPES-NaOH pH 7.0, 10 mM MgCl_2_, 300 mM NaCl, 1 mM DTT). Data were evaluated by comparing elution profiles in the presence of different DNAs and calculating ratios of peak A260 mAU values for complexes II and I. Indicated fractions were additionally mixed with orange G loading dye (0.02% [w/v] orange G, 5% [v/v] glycerol), and protein-DNA complexes and free DNA were resolved at 4 °C on 6% native 0.5x TBE polyacrylamide gels, pre-equilibrated by running at 200 V for 60 min. Upon sample loading, gels were run at 90 V for 70 min, followed by 75 V for 10 min. Gels were stained with SYBR Gold (ThermoFisher Scientific).

### Mass photometry

Samples of TnsAB WT, evoTnsAB, TnsB WT and evoTnsB for mass photometry were prepared in 25 µL reaction volumes on ice, with or without the paired-end complex (PEC) DNA construct (generated by annealing oligonucleotides o577 and o578; sequences provided in **Supplementary Table 2**). Reactions contained 12.5 µL of 50 µM protein, 3.57 µL high salt buffer (25.3 mM HEPES-NaOH pH 7.0, 410 mM NaCl, 36.4 mM MgCl_2_, 0.7 mM DTT), and 4.2 µL PEC DNA (25 µM), and were slowly diluted with 4.73 µL of low salt buffer (15.86 mM HEPES-NaOH pH 7.0, 26.43 mM MgCl_2_) to produce assembly buffer at a final composition of 8.3 mM HEPES-NaOH pH 7.0, 10 mM Tris-HCl pH 7.0, 317 mM NaCl, 10.2 mM MgCl_2_, 0.6 mM DTT, containing 25 µM of respective protein and 4.2 µM PEC DNA. Apo-protein reactions were assembled in an analogous way, using TE50 buffer instead of PEC DNA. Reactions were incubated on ice for 30 min. Mass photometry measurements were performed on a Refeyn TwoMP instrument (Refeyn). Assembled reactions were diluted to 50 nM in mass photometry dilution buffer (18.3 mM HEPES-NaOH pH 7.0, 317 mM NaCl and 10.2 MgCl_2_), immediately prior to measurement. Calibration curve was generated using BSA (Sigma Aldrich; measurements of TnsAB WT and evoTnsAB) or MassFerence P1 Calibrant (Refeyn; measurements of TnsB WT and evoTnsB) diluted in the mass photometry dilution buffer. For mass photometry measurements of complexes eluting from analytical SEC (shown in **Supplementary Fig. 9d**), indicated fractions obtained from the TnsAB + DNA WT sample were kept on ice and analyzed in their native state. Complex I fraction was diluted 1:40, while complex II fraction was diluted 1:20, both using 1x PBS (Roth), immediately prior to measurement. Calibration curve for this experiment was generated using MassFerence P1 Calibrant resuspended in 1x PBS. All data were analysed using the DiscoverMP software (v2024 R2.1).

### *In vitro* excision assay

*In vitro* excision reactions (20 µL) were performed in a reaction buffer containing 20 mM HEPES-NaOH pH 7.0, 5 mM Tris-HCl pH 7.0, 400 mM NaCl, 15% DMSO [v/v], 1% glycerol [v/v], 50 µg mL^-1^ BSA, 1.25 mM DTT. The addition of DMSO was inspired by a previous report on type V-K TnsB^66^. TnsAB WT, TnsB WT or their mutants (400 nM) were pre-incubated with 5 nM of supercoiled pDonor plasmid for 30 min at room temperature. DNA cleavage reactions were initiated by adding 20 mM of divalent metal ions (MnCl_2_, MgCl_2_ or CaCl_2_, as indicated in the figures) and incubated at 30 °C for 240 min. The divalent ion concentrations were chosen based on prior biochemical studies of *Ec*Tn7^6,58^. Time-course excision assays using TnsAB WT and evoTnsAB were performed under the same conditions, and incubated at 30 °C or 37 °C (as indicated in the figures) and quenched at the indicated time points using 23.8 mM EDTA pH 8.0 (final concentration). All excision reactions were terminated by addition of 1 mg mL^-1^ proteinase K, followed by incubation at 37 °C for 60 min. DNA species were purified by ethanol precipitation and resuspended with 17.5 µL ultrapure water mixed with DNA loading dye (2 mM Tris-HCl pH 7.5, 10 mM EDTA, 10% glycerol, 0.005% bromophenol blue, 0.005% xylene cyanol FF final concentrations). Samples were resolved at room temperature on 1% agarose gels in 1x TAE buffer. DNA was visualised by staining with Safe DNA gel stain (APExBIO). Excision assays with TnsC were performed as described above, except that TnsC (at the indicated concentrations) was co-incubated with TnsAB and 5 nM wild type pDonor for 30 min at room temperature. Reactions were initiated by addition of 20 mM MnCl_2_ or MgCl_2_, and incubated at 30 °C for 240 min. For assays with mutant transposon end constructs, DNA pellets were resuspended in 6 µL ultrapure water with DNA loading dye and resolved at room temperature on 6% native 0.5x TBE polyacrylamide gels in 0.5x TBE. These gels were stained with SYBR Gold (ThermoFisher Scientific).

Band intensities were quantified in Fiji^73^. To calculate fraction of produced excision product or fraction of remaining supercoiled plasmid, the density of the corresponding band was divided by the total DNA signal in the lane. To obtain normalized processing efficiency values, the sum of raw densities for the relaxed, linear, backbone and transposon bands from within one lane were divided by the total DNA signal in that lane, and values were subsequently normalized by treating the mean processing efficiency by TnsAB WT in Mn^2+^ on WT transposon ends as 100% (without TnsC in **Supplementary Fig. 7b**-**c**). For assays with mutant transposon ends, values were normalized across replicates to the density of the linear DNA band (3 kb) obtained from reactions with WT transposon ends and TnsAB WT. Excision efficiency was then calculated by dividing the normalized density of the backbone DNA band (2 kb) resulting from excision on mutant transposon ends by the mean normalized density obtained on WT transposon ends. Data were plotted and statistical analyses performed in GraphPad Prism version 10.2.0 (GraphPad Software, Boston, Massachusetts USA, www.graphpad.com).

### Determination of TnsAB cleavage sites

Analysis of transposase-induced cleavage positions was adapted from a previously described protocol^74^. Excision reactions were prepared as described above, using 400 nM TnsAB WT and 5 nM pDonor_WT, and incubated for 240 min following the addition of 20 mM MnCl_2_ or MgCl_2_. Reactions were stopped by transferring samples to -20 °C without proteinase K treatment or ethanol precipitation. Reactions were subjected to adaptor ligation, where each sample was mixed with 20 pmol single-stranded DNA oligonucleotide (Link1/2), heated to 95 °C for 3 min, cooled on ice for 3 min, and supplemented with 1x T4 RNA ligase buffer, 1 mM ATP, 3.3% PEG8000 and T4 RNA ligase 1 (NEB) following the manufacturer’s instructions. Ligation reactions were incubated at room temperature overnight and inactivated at 65 °C for 15 min. To increase DNA recovery, five ligation reaction per condition were pooled and purified using a PCR cleanup kit (Machery-Nagel). To obtain a qualitative analysis of the cleavage products, DNA was used as a template for nested PCR (primer set including o158-o168 are listed in **Supplementary Table 2**). In the first amplification round, the program was 98 °C for 3 min; 30 cycles of 98 °C for 30 s, 55 °C for 30 s, and 72 °C for 15 s; followed by a final extension at 72 °C for 10 min. For the second round, 50-fold diluted products from the first PCR were used as templates. The program was 98 °C for 3 min; 30 cycles of 98 °C for 30 s, 68 °C for 30 s, and 72 °C for 20 s; followed by a final extension at 72 °C for 10 min. Resulting PCR products were purified using PCR cleanup kit (Machery-Nagel) and visualised in 1% agarose gel (**Supplementary Fig. 2c**).

Cleavage sites were mapped using Illumina sequencing. For library preparation, adaptor-ligated DNA was amplified in two PCR steps. PCR1 incorporated part of the sequencing primer binding site (using primers o681, o704, and o683-o685 listed in **Supplementary Table 2**). Reactions (25 µL) contained 250 pg template DNA, 0.5 µM of each primer, and 0.02 U µL^-1^ Q5 high-fidelity polymerase (NEB). Cycling conditions were: 72 °C for 1 min; 98 °C for 30 s; 24 cycles of 98 °C for 10 s, 65–67 °C for 20 s (65, 66, 67, and 67 °C for samples S1-S4, respectively), and 72 °C for 10 s; followed by a final extension at 72 °C for 1 min. PCR1 products were purified using the PCR purification protocol of the MiniPEx kit (Vienna BioCenter Molecular Tool Development shop). PCR2 incorporated the remaining primer binding sequences, TruSeq-style indexes, and flow-cell adapters (UD# primers listed in **Supplementary Table 2**). Reactions (20 µL) contained 0.5 pM PCR1 product, 1.25 µM of each primer, and 0.02 U µL^-1^ Q5 high-fidelity polymerase (NEB). Cycling conditions were: 72 °C for 5 min; 98 °C for 3 min; 25 cycles of 98 °C for 10 s, 67 °C for 30 s, and 72 °C for 30 s; followed by a final extension at 72 °C for 1 min. PCR2 products were purified with magnetic beads using the Select-a-Size DNA Clean & Concentrator MagBead kit (Zymo Research) at a 7:10 bead-to-sample ratio.

Final libraries were sequenced on a NovaSeq X 10B XP flow cell (Illumina) as 150 bp paired-end reads. FASTQ file quality was assessed with FastQC v0.11.9. Sequencing was carried out by the Next Generation Sequencing Facility at the Vienna BioCenter Core Facilities (VBCF). Sequencing reads were analyzed using in-house Bash and Python scripts (see **Code availability**). Briefly, reads were quality-filtered and merged with fastp^75^, then dereplicated. Reads lacking the ligated adaptor sequence or containing ≤15 additional bases were discarded. For the remaining reads, the adaptor sequence was trimmed, and reads were mapped to the pDonor plasmid with Bowtie2^76^. Mapped reads with mismatches in the first 10 nt were excluded to avoid misassignment of cleavage positions. For all retained reads, the DNA cleavage site was defined as between the most 5′ mapped nucleotide and the nucleotide immediately upstream.

### Cryo-EM sample preparation and data collection

A TnsAB WT stock at 75 µM was prepared in 20 mM Tris-HCl pH 7.5, 500 mM NaCl, and 1 mM DTT. The DNA constructs (o577/o578 for the RE samples, oGF254/oGF255 for the LE sample; sequences provided in **Supplementary Table 2**) were generated as described above and diluted to 50 µM in ultrapure water.

Complexes were assembled on ice by mixing 3.6 µL of high-salt buffer (1050 mM KCl, 10 mM HEPES, 10 mM MgCl_2_, or MnCl_2_, 1 or 0 mM DTT) with 2.1 µL DNA from the 50 µM stock. 8.3 µL of TnsAB from the 75 µM stock were then added, and the mixture was brought to 25 µL by slowly adding dilution buffer (20 mM HEPES, 20 mM MgCl_2_, or MnCl_2_, 1 or 0 mM DTT) while mixing gently. The final sample contained 25 µM TnsAB, 4.2 µM DNA, 317.0 mM total NaCl plus KCl, 18.3 mM HEPES-KOH plus Tris-HCl, 10.2 mM MgCl_2_, or MnCl_2_ and 0.6 or 0.0 mM DTT. DTT was omitted from the buffer for the Mn^2+^-containing sample and used only for the Mg^2+^-containing ones. In each case, the sample was incubated at 4 °C for 30 min, centrifuged at 4 °C, 14 000g for 10 min, and the supernatant was transferred to a pre-chilled tube. Based on analytical SEC and mass photometry experiments, we anticipated the presence of both monomeric and higher-order DNA-bound species under these conditions. For cryo-EM analysis, the assembled reaction was therefore used directly without preparative purification to preserve the full spectrum of assembly states. Homogeneous subsets of complexes were subsequently isolated by computational classification during image processing, as described below.

For vitrification, freshly glow-discharged Quantifoil R 1.2/1.3 Au 300-mesh grids (60 s glow discharge) were used. A 2.5 µL aliquot of the sample was applied, blotted for 2.5 s (TnsAB-LE-MgCl_2_ sample), 3 s (TnsAB-RE-MgCl_2_ and TnsAB-RE-MnCl_2_ samples), or 3.5 s (TnsAB-LE-MgCl_2_ and TnsAB-RE-MnCl_2_ samples) at 100% humidity, 4 °C, and plunge-frozen into an ethane/propane cryogen mixture using a Vitrobot Mark IV plunger (FEI) and stored in liquid nitrogen. Cryo-EM data were collected on an FEI Titan Krios G3i (University of Zurich, Switzerland) operating at 300 kV and equipped with a Gatan K3 direct electron detector in super-resolution counting mode. For TnsAB-RE-MgCl_2_, 3 449 movies were acquired at ×130000 magnification, yielding a super-resolution pixel size of 0.325 Å. Each movie consisted of 37 subframes with a cumulative exposure of 62.130 e^-^/Å^2^. EPU Automated Data Acquisition Software for Single Particle Analysis (Thermo Fisher Scientific) was used for data acquisition, with three shots per hole and a defocus range of -1.0 to -2.4 μm in 0.2-μm increments. For TnsAB-LE-MgCl_2_ and TnsAB-RE-MnCl_2_, 15 169 and 8 333 movies, respectively, were collected at ×130 000 magnification from two grids per sample that differed only in blotting time, yielding a super-resolution pixel size of 0.325 Å. Each movie consisted of 37 subframes with a cumulative exposure of 64.201 e-/Å^2^. EPU Automated Data Acquisition Software for Single Particle Analysis (Thermo Fisher Scientific) was used for data acquisition, with three shots per hole and a defocus range of -0.8 to -2.6 μm in 0.3-μm increments.

### TnsAB-RE-Mg: cryo-EM data processing and model building

Obtained cryo-EM data was processed using cryoSPARC^77^ (v4.7). 3 440 exposures were imported to cryoSPARC live for real-time processing, including patch motion and patch CTF correction. 1 095 968 particles were subsequently detected using the blob picker function (100-250 Å diameter), extracted in a box size of 480 pixels (Fourier-cropped to 160 pixels) and classified into 30 classes using 2D classification. Nine classes corresponding to 287 299 particles were selected and used in a 2-class ab-initio reconstruction yielding a junk class and a recognizable TnsAB volume. The respective 210 748 particles were re-extracted in a box size of 480 pixels in unbinned form (resulting in 208 483 particles) and fed into an initial non-uniform refinement calculation, applying C2 symmetry constraints. In parallel, contaminants were excluded from the re-extracted particle set employing the micrograph junk detector function and corresponding exposures were manually curated based on their NCC and power score. The final 2.78 Å (GSFSC resolution) reconstruction was obtained by non-uniform refinement (enforcing C2 symmetry) of the curated particle stack (203 418 particles), using the refined map obtained in the previous round of non-uniform refinement as initial volume. A detailed processing workflow is presented in **Supplementary Fig. 4a**-**e**.

An initial model of monomeric *Pse*TnsAB was generated using AlphaFold3^78^ and rigid body-fitted in the cryo-EM density map using UCSF ChimeraX^79^, followed by real-space refinement in Phenix^80^. The two identical DNA duplexes, consisting of the transferred (positions -29 to -1) and non-transferred strands (positions -3 to 29), were manually built in Coot^81^ with the corresponding density being well enough defined to confidently assign their register. The overall model was manually refined in Coot using secondary structure, side chain rotamer, Ramachandran, and LibG^82^ nucleic acid restraints, followed by a final refinement in Phenix. Key refinement statistics are reported in **Supplementary Table 1**.

### TnsAB-LE-Mg: cryo-EM data processing and model building

Obtained cryo-EM data was processed using cryoSPARC^77^ (v4.7). 15 223 exposures were imported to cryoSPARC live for real-time processing, including patch motion and patch CTF correction. 4 935 000 particles were detected using the blob picker function (100-250 Å diameter), extracted in a box size of 480 pixels (Fourier-cropped to 160 pixels) and classified into 50 classes using 2D classification. 11 classes corresponding to 1 192 445 particles were selected and used in a 2-class ab-initio reconstruction yielding a junk class and a recognizable TnsAB volume. The respective 866 938 particles were used in a 2-class heterogeneous refinement using the ab-initio TnsAB reconstruction as an input for both classes. The refinement yielded one poorly and one well defined TnsAB volume. The 584 938 particles corresponding to the best volume were re-extracted in a box size of 480 pixels (unbinned), resulting in 578 675 particles. Volume and particles were used for a non-uniform refinement job applying C2 symmetry constraints, which resulted in a final 2.51 Å (GSFSC resolution) reconstruction. A detailed processing workflow is presented in **Supplementary Fig. 10b**-**f**.

The structure of TnsAB-RE-MgCl_2_ was rigid body-fitted in the cryo-EM density map using UCSF ChimeraX^79^ and used as initial model. The sequence of the DNA duplexes was adjusted to match the LE DNA construct. The model was manually refined in Coot using secondary structure, side chain rotamer, Ramachandran, and LibG^82^ nucleic acid restraints, followed by a final refinement in Phenix. Key refinement statistics are reported in **Supplementary Table 1**.

### TnsAB-RE-Mn: cryo-EM data processing and model building

Obtained cryo-EM data was processed using cryoSPARC^77^ (v4.7). Initial processing (corrections, picking, 2D classification) was carried out separately for the two grids. 3 004 exposures from grid 1 (3.5 s blotting time) and 5 536 exposures from grid 2 (2.5 s blotting time) were imported to cryoSPARC live for real-time processing, including patch motion and patch CTF correction. 898 926 particles from grid 1 and 1 343 358 particles from grid 2 were detected using the blob picker function (100-250 Å diameter), extracted in a box size of 480 pixels (Fourier-cropped to 160 pixels) and classified into 50 classes using 2D classification. 4 classes from grid 1 and 4 classes from grid 2, corresponding to a total of 204 563 particles, were selected and used in a 2-class ab-initio reconstruction, which yielded a junk class and a recognizable TnsAB volume. The volume and its respective 161 236 particles were re-extracted in a box size of 480 pixels (resulting in a final 159 536 particles set) and used in a non-uniform refinement applying C2 symmetry. The refinement resulted in a final 2.86 Å (GSFSC resolution) reconstruction. A detailed processing workflow is presented in **Supplementary Fig. 14a**-**e**.

The structure of TnsAB-RE-MgCl2 was rigid body-fitted in the cryo-EM density map using UCSF ChimeraX^79^ and used as initial model. The structure was manually refined in Coot using secondary structure, side chain rotamer, Ramachandran, and LibG^82^ nucleic acid restraints, followed by a final refinement in Phenix. Key refinement statistics are reported in **Supplementary Table 1**.

### Transposition assays in *E. coli*

A *Pse*CAST pDonor_transposition_WT plasmid was first transformed into *E. coli* BL21 (DE3) cells (NEB). Single colonies were inoculated into liquid culture, and the resulting strains were made chemically competent using standard methods. *Pse*CAST pEffector plasmids encoding a crRNA targeting the lacZ locus (crRNA-4)^17^ and all CAST subunits were transformed into the resulting strain and plated on double antibiotic LB-agar plates containing 100 µg mL^-1^ carbenicillin and 100 µg mL^-1^ spectinomycin. After approximately 20 h of growth, surviving colonies were scraped from the plates, resuspended in fresh LB medium, and re-plated on double antibiotic LB-agar plates supplemented with 0.1 mM IPTG to induce *Pse*CAST protein expression. Cells were incubated for an additional 18 – 20 hours, scraped, and resuspended in LB medium. Cells were then lysed, and integration efficiencies were measured using qPCR as previously described^17^. Sequences of used plasmids, crRNA-4, and oligos used for qPCR are listed in **Supplementary Table 2**.

### Statistics and reproducibility

Sample sizes and statistical tests used are indicated in respective figure legends, and all normalization procedures are described in the **Methods** section. For biochemical experiments, measurements were performed in triplicates on independent biological samples, and are shown in quantification graphs, or as a representative gel. For *in vivo* transposition assays, integration efficiencies are presented normalized to the wild-type condition. The mean value, all biological replicates and ± standard deviation are plotted. No statistical method was used to predetermine sample size, and no data were excluded from the analyses. Data from all replicates showed similar results.

## Data availability

The cryo-EM maps and corresponding atomic models of the *Pse*TnsAB paired-end complexes have been deposited in the Electron Microscopy Data Bank (EMDB) and the Protein Data Bank (PDB) under accession codes EMD-55638 (TnsAB-RE-MgCl_2_), EMD-57834 (TnsAB-LE-MgCl_2_), EMD-57833 (TnsAB-RE-MnCl_2_), and 9T7L (TnsAB-RE-MgCl_2_), 30JW (TnsAB-LE-MgCl_2_), 30JV (TnsAB-RE-MnCl_2_), respectively. Source data are provided with this paper.

## Code availability

Data and code related to next-generation sequencing experiments are available at: Swartjes, T. (2025), “Transposon end recognition and excision mechanisms of type I-F CRISPR-associated transposases”, Mendeley Data, V1, https://doi.org/10.17632/r5ccxsc8wg.1. This entry can be accessed via the temporary link: https://data.mendeley.com/preview/r5ccxsc8wg?a=6b6c79d6-7423-4325-b1c4-19b14c6e55c6.

