## Supplementary Information for "Transposon end recognition and excision mechanisms of type I-F CRISPR-associated transposases"

#### Supplementary Figures

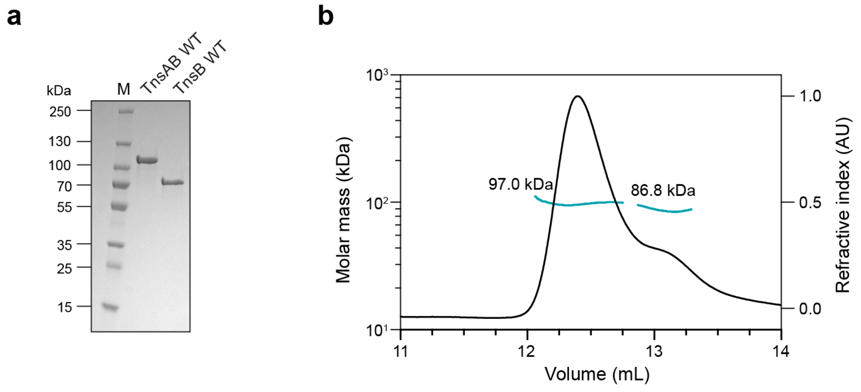


##### Supplementary Fig. 1 | Purification and characterization of *Pse*TnsAB

(**a**) SDS-PAGE analysis of the purified *Pse*TnsAB and *Pse*TnsB proteins. WT, (wild-type). M, marker. (**b**) Size exclusion chromatography coupled with multi-angle light scattering (SEC-MALS) of purified *Pse*TnsAB. The results align with the theoretical molecular weight of a monomer (97.65 kDa).

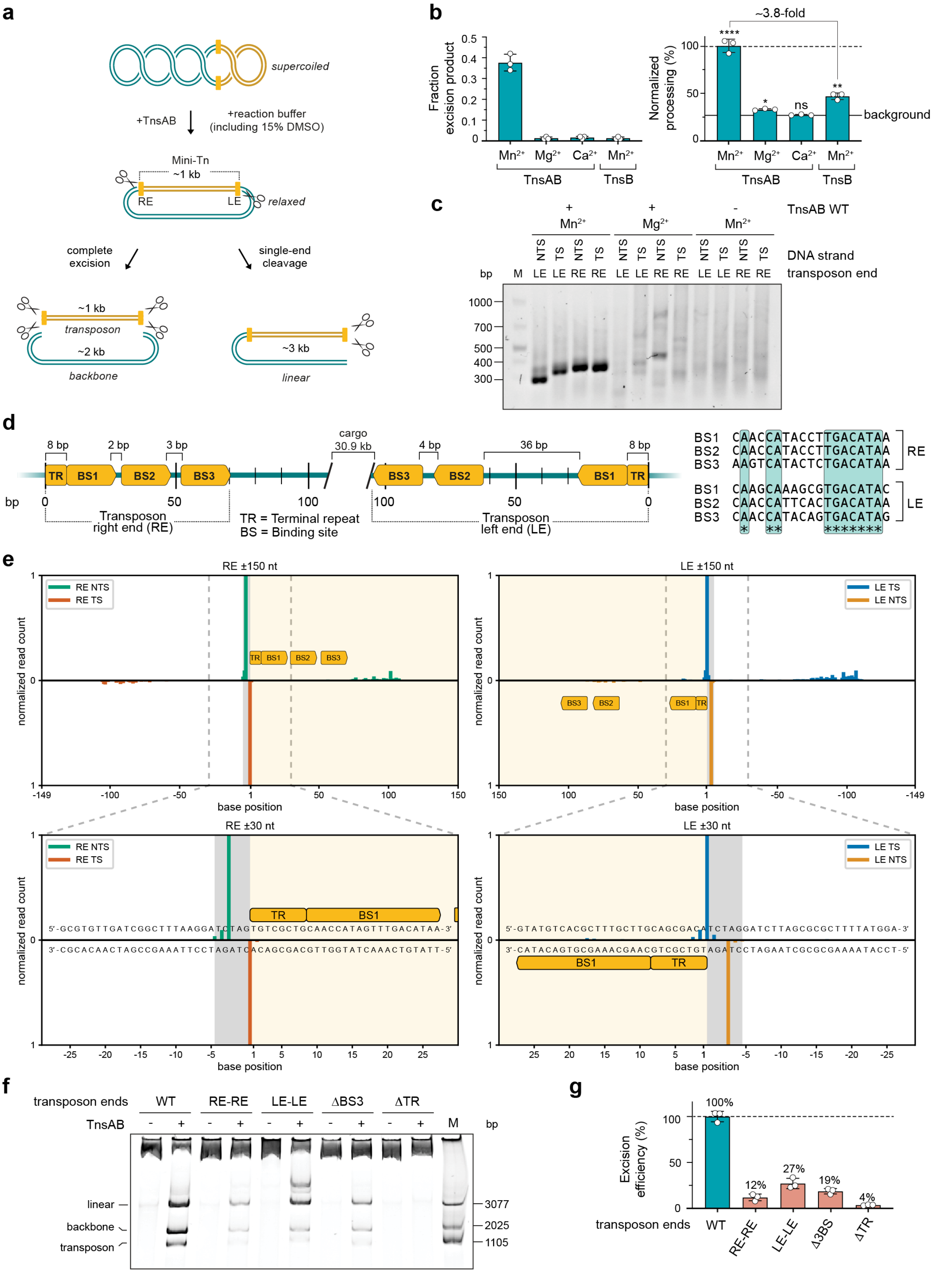


**Supplementary Fig. 2 | Reconstitution and analysis of *Pse*TnsAB cleavage activity *in vitro***

(**a**) Schematic of the *in vitro* excision assay. Mini-Tn, mini-transposon. LE/RE, left/right end. (**b**) Quantification of *in vitro* excision assay from three independent experiments (see **Methods** for details). Relative amounts of DNA species were determined by densitometry. Left: Quantification of excision. Data show mean ± s.d. of plasmid backbone (~2 kb-long excision product). Right: Quantification of total

DNA processing relative to WT TnsAB in the presence of $Mn^{2+}$. Data show mean ± s.d. of all processed DNA species. The solid horizontal line indicates the mean background level of processed plasmid in the no-protein control. Statistical significance was assessed using unpaired two-tailed *t*-tests. P-values above each graph indicate comparisons to the no-protein control. P-values: *p < 0.05; **p < 0.01; ****p < 0.0001; ns, not significant. (**c**) Nested PCR products resolved by agarose gel electrophoresis. TS, transferred strand; NTS, non-transferred strand. (**d**) Left: Arrangement of transposon ends. TR, terminal repeat; BS, binding sites. Minor ticks every 10 bp. Right: Alignment of RE and LE BS sequences, with conserved nucleotides marked by asterisks. (**e**) Cleavage positions inferred from Illumina sequencing. y-axis, normalized read counts per cut site. Bars above/below axis denote top/bottom strand cuts. Transposon end regions are shaded yellow; target site duplications (TSDs) grey. TRs and BSs are highlighted in bright yellow. Windows cover ≥99.999% of reads for each strand/end. (**f**) *In vitro* excision assay with mutant plasmids carrying wild-type (WT), symmetrized, or truncated ends. DNA species are resolved by native PAGE. Representative experiment is shown. ΔBS3, lacking BS3 at both ends; ΔTR, lacking terminal repeats at both ends; M, marker. (**g**) Quantification of *in vitro* excision assay with mutant plasmids. Mean ± s.d. of plasmid backbone (~2 kb-long excision product) normalized to WT from three independent experiments, as measured by densitometry (see **Methods** for details).

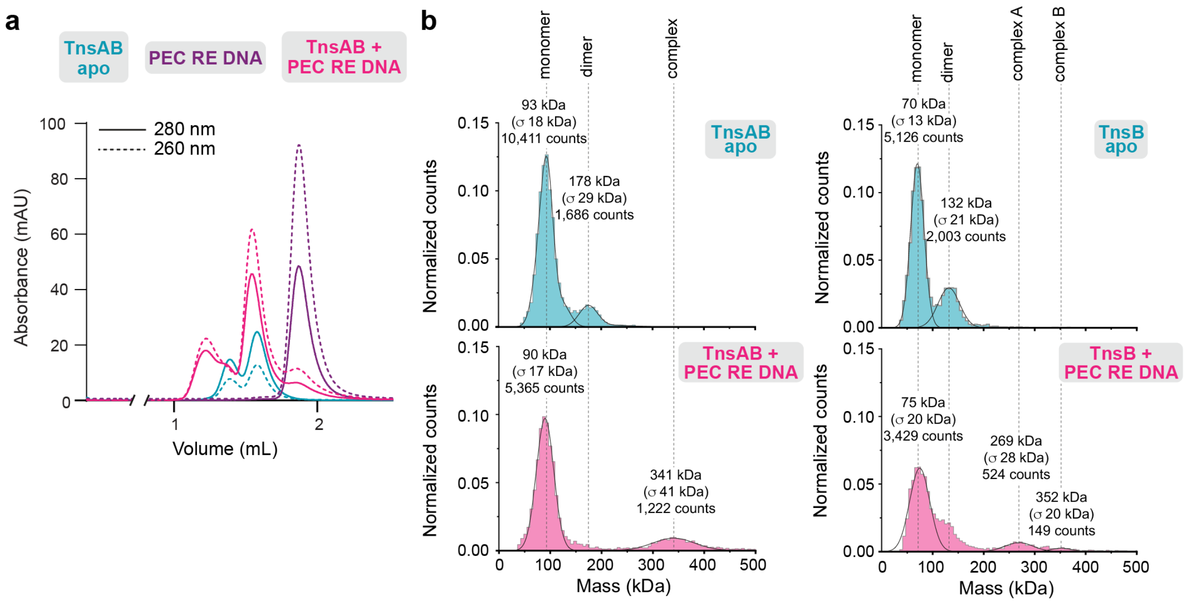


**Supplementary Fig. 3 | Assembly of the *Pse*TnsAB PEC on right end DNA**

(**a**) Size exclusion chromatography (SEC) profile of *Pse*TnsAB paired-end complex (PEC) reconstituted on transposon right end DNA (PEC RE DNA), used for structural analysis. (**b**) Mass photometry data of *Pse*TnsAB (left) and *Pse*TnsB (right) in its apo (top) and PEC (bottom) states, with masses and respective particle population counts indicated. The theoretical molecular weight of a TnsAB monomer is 97.65 kDa, and of TnsB is 71.29 kDa.

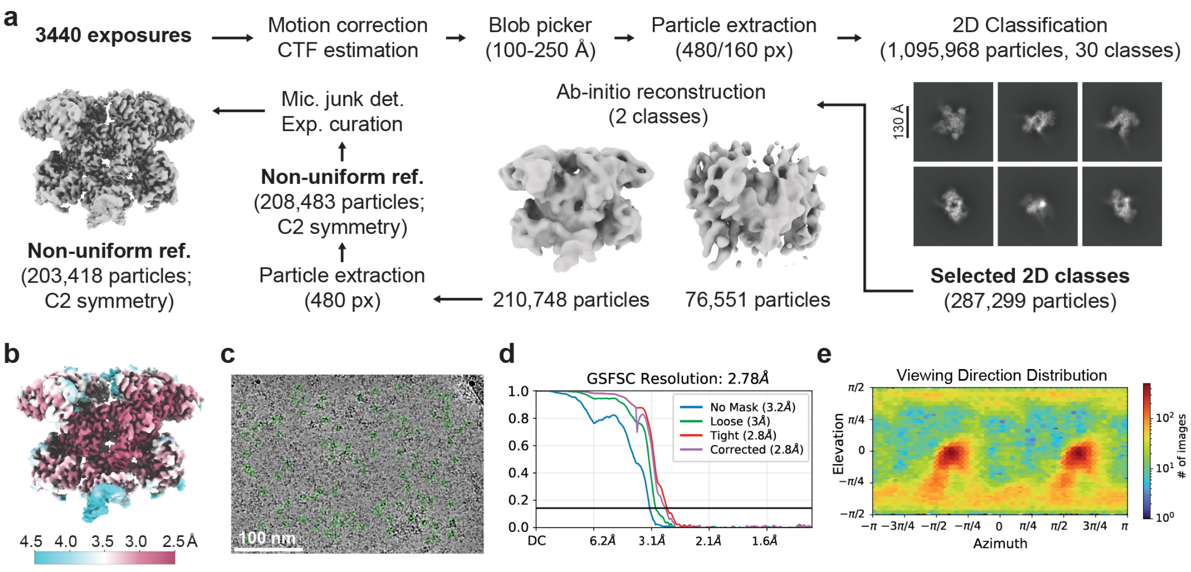


**Supplementary Fig. 4 | Cryo-EM data processing workflow for the *Pse*TnsAB PEC assembled on right end DNA in the presence of $Mg^{2+}$**

(**a**) Cryo-EM image processing workflow for the *Pse*TnsAB paired-end complex (PEC) reconstituted on transposon right end DNA. (**b**) Cryo-EM density maps colored by local resolution. (**c**) Representative cryo-EM micrograph. Scale bar, 100 nm. (**d**) Fourier Shell Correlation (FSC) of the reconstruction calculated from two independently refined half-maps. The gold-standard cutoff (FSC = 0.143) is marked with a horizontal black line. (**e**) Angular distribution showing the range of observed particle orientations.

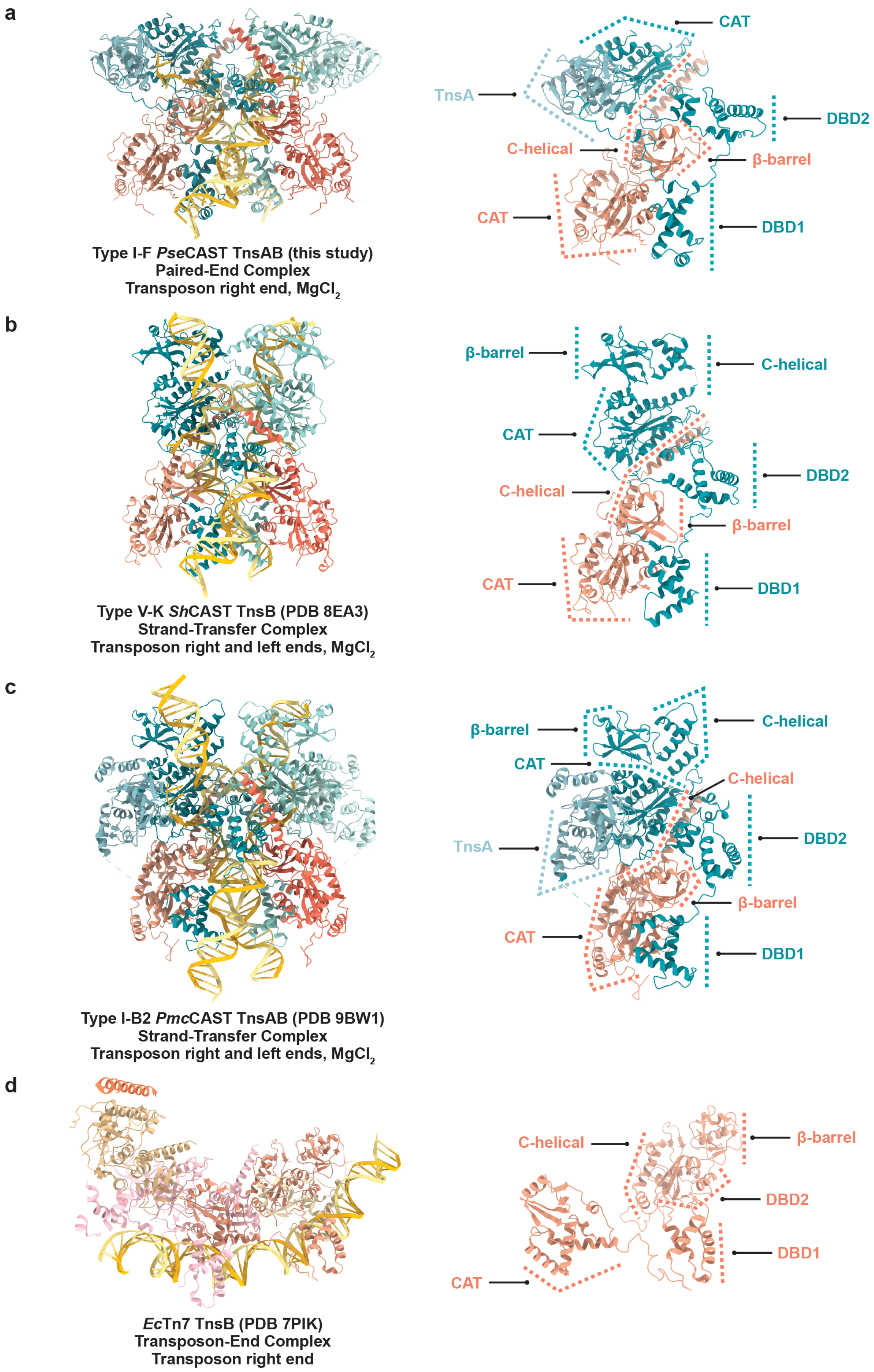
a
CAT
TnsA
DBD2
C-helical
β-barrel
CAT
DBD1
Type I-F *Pse*CAST TnsAB (this study)
Paired-End Complex
Transposon right end, $MgCl_2$
b
β-barrel
C-helical
CAT
DBD2
C-helical
β-barrel
CAT
DBD1
Type V-K *Sh*CAST TnsB (PDB 8EA3)
Strand-Transfer Complex
Transposon right and left ends, $MgCl_2$
c
β-barrel
C-helical
CAT
C-helical
TnsA
DBD2
β-barrel
CAT
DBD1
Type I-B2 *Pmc*CAST TnsAB (PDB 9BW1)
Strand-Transfer Complex
Transposon right and left ends, $MgCl_2$
d
C-helical
β-barrel
DBD2
DBD1
CAT
*Ec*Tn7 TnsB (PDB 7PIK)
Transposon-End Complex
Transposon right end

**Supplementary Fig. 5 | Structural comparison of *Pse*TnsAB PEC, type V-K *Sh*TnsB STC, type I-B2 *Pmc*TnsAB STC, and Tn7 TnsB transposon-end complex**

(**a-d**) Comparison of the *Pse*TnsAB paired-end complex (PEC) (**a**) with type V-K *Sh*TnsB[1] (**b**) and type I-B2 *Pmc*TnsAB[2] (**c**) strand-transfer complexes (STCs), and the *E. coli* Tn7 (*Ec*Tn7) TnsB transposon-end complex[3] (**d**). Left panels show overall architecture, right panels highlight domain differences. DBD, DNA binding domain; CAT, catalytic domain.

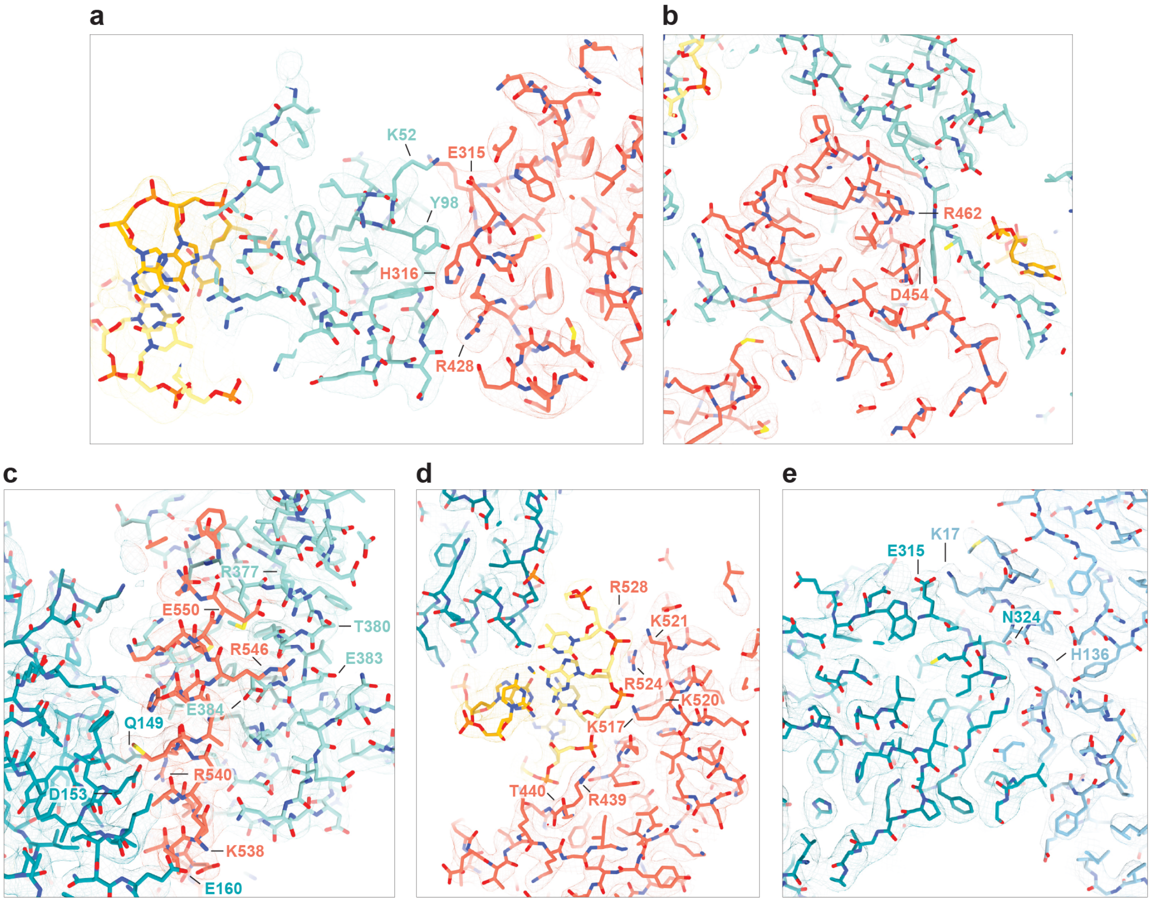


**Supplementary Fig. 6 | Local model-to-map fit for regions involved in TnsB interprotomer interactions and TnsA-TnsB interfaces**

Panels show a zoomed view of the structural regions corresponding to the indicated panels in **Fig. 3**, with the atomic model shown in stick representation overlaid with cryo-EM density displayed as a transparent mesh at the following contour levels: (**a**) 0.04, (**b**) 0.07, (**c**) 0.06, (**d**) 0.07, (**e**) 0.05. Residues highlighted in the corresponding panels of **Fig. 3** are labeled for reference, with consistent coloring. Panel correspondence: (**a**) **Fig. 3a**; (**b**) **Fig. 3b**; (**c**) **Fig. 3c**; (**d**) **Fig. 3d**; (**e**) **Fig. 3e**.

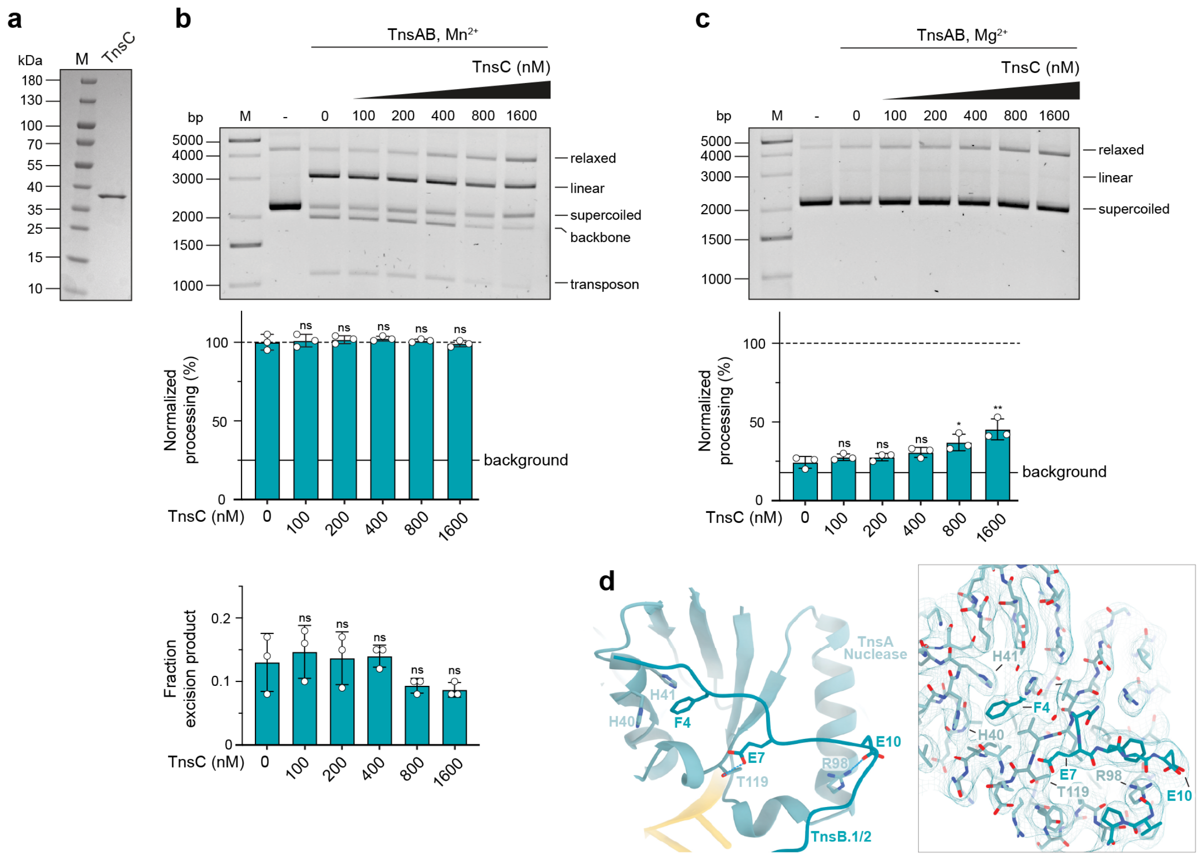


#### Supplementary Fig. 7 | *In vitro* cleavage activity in the presence of *Pse*TnsC and detailed view of TnsA-TnsB interactions

(**a**) SDS-PAGE analysis of purified *Pse*TnsC protein. M, marker. (**b-c**) *In vitro* excision assay with *Pse*TnsAB in the presence of increasing concentrations of *Pse*TnsC and either $Mn^{2+}$ (**b**) or $Mg^{2+}$ (**c**). Top: representative agarose gels showing resolved DNA species. -, no-protein control; M, marker. (**b**) Middle and bottom: Quantification of *in vitro* excision assay from three independent experiments (see **Methods** for details). Relative amounts of DNA species were determined by densitometry. Middle: Quantification of total DNA processing normalized to WT TnsAB in the presence of $Mn^{2+}$ without TnsC. Data show mean ± s.d. of all processed DNA species. The solid line indicates the mean background level of processed plasmid in the no-protein control. Bottom: Quantification of excision. Data show mean ± s.d. of plasmid backbone (~2 kb-long excision product). (**c**) Bottom: Quantification from three independent experiments, as in (**b**, middle panel), for $Mg^{2+}$ conditions. Total DNA processing is normalized to WT TnsAB in the presence of $Mg^{2+}$ without TnsC. Statistical analysis was conducted using unpaired two-tailed *t*-test. P-values above each graph indicate comparisons with the corresponding negative control. P-values: *p < 0.05; **p < 0.01; ns, not significant. (**d**) Left: Close-up view of the specific contacts between the *Pse*TnsB N-terminus and *Pse*TnsA. Dashed lines represent interaction distances ≤ 3.5 Å. Right: Corresponding atomic model shown in stick representation overlaid with cryo-EM density shown as a transparent mesh and at a contour level of 0.004.

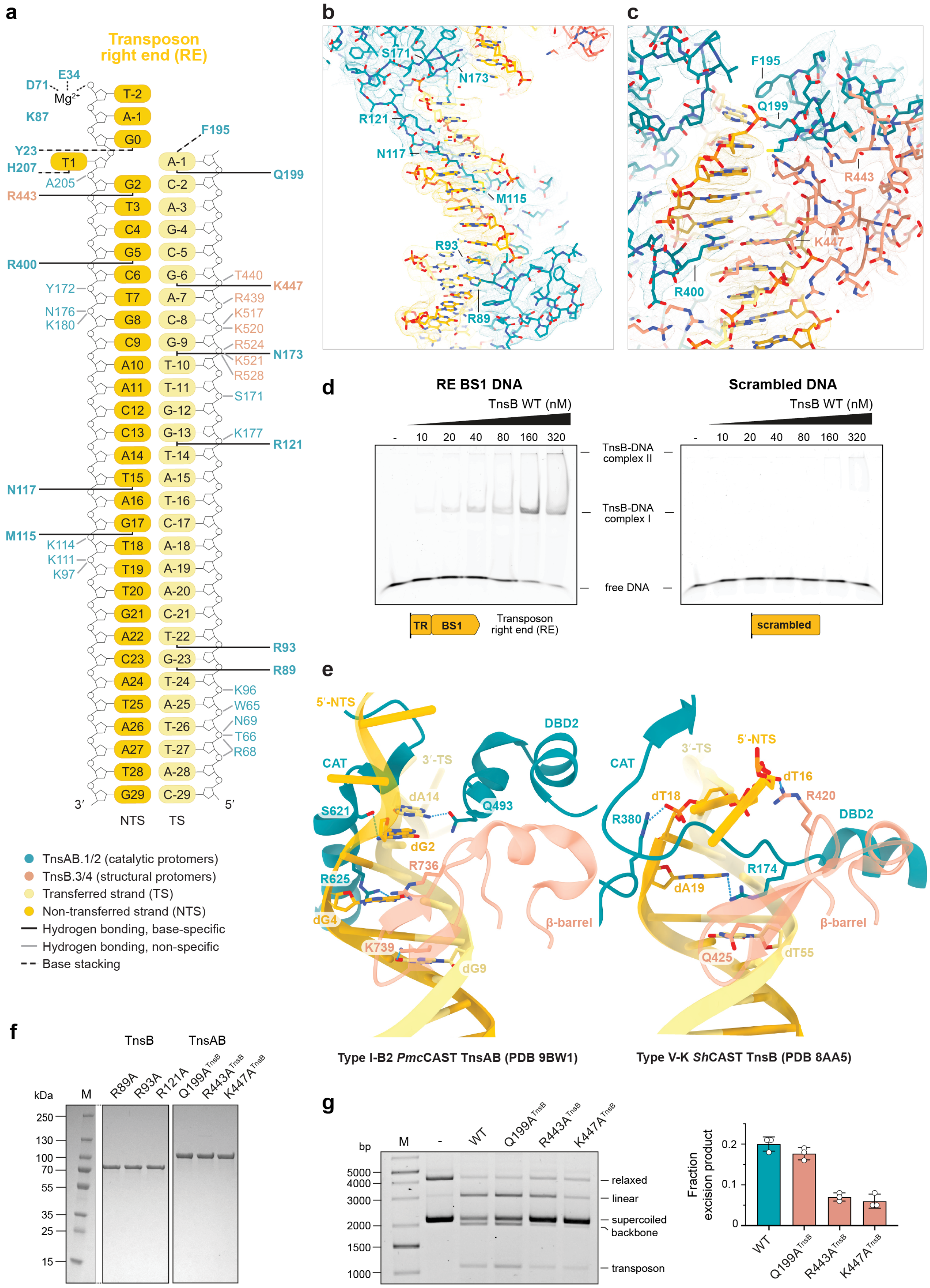
a
Transposon right end (RE)
D71
E34
Mg2+
K87
Y23
H207
A205
R443
R400
Y172
N176
K180
N117
M115
K114
K111
K97
T-2
A-1
G0
T1
G2
T3
C4
G5
C6
T7
G8
C9
A10
A11
C12
C13
A14
T15
A16
G17
T18
T19
T20
G21
A22
C23
A24
T25
A26
A27
T28
G29
A-1
C-2
A-3
G-4
C-5
G-6
A-7
C-8
G-9
T-10
T-11
G-12
G-13
T-14
A-15
T-16
C-17
A-18
A-19
A-20
C-21
T-22
G-23
T-24
A-25
T-26
T-27
A-28
C-29
F195
Q199
T440
K447
R439
K517
K520
R524
N173
K521
R528
S171
K177
R121
R93
R89
K96
W65
N69
T66
R68
3′
5′
NTS
TS
TnsAB.1/2 (catalytic protomers)
TnsB.3/4 (structural protomers)
Transferred strand (TS)
Non-transferred strand (NTS)
Hydrogen bonding, base-specific
Hydrogen bonding, non-specific
Base stacking
b
S171
N173
R121
N117
M115
R93
R89
c
F195
Q199
R443
K447
R400
d
RE BS1 DNA
TnsB WT (nM)
- 10 20 40 80 160 320
TnsB-DNA complex II
TnsB-DNA complex I
free DNA
TR
BS1
Transposon right end (RE)
Scrambled DNA
TnsB WT (nM)
- 10 20 40 80 160 320
scrambled
e
5′-NTS
DBD2
CAT
3′-TS
dA14
Q493
S621
dG2
R736
R625
dG4
β-barrel
K739
dG9
Type I-B2 PmcCAST TnsAB (PDB 9BW1)
CAT
3′-TS
5′-NTS
dT16
dT18
R420
R380
DBD2
R174
dA19
β-barrel
Q425
dT55
Type V-K ShCAST TnsB (PDB 8AA5)
f
TnsB
TnsAB
kDa
M
R89A
R93A
R121A
Q199ATnsB
R443ATnsB
K447ATnsB
250
130
100
70
55
35
25
15
g
bp
M
-
WT
Q199ATnsB
R443ATnsB
K447ATnsB
5000
4000
3000
2000
1500
1000
relaxed
linear
supercoiled backbone
transposon
Fraction excision product
0.2
0.1
0
WT
Q199ATnsB
R443ATnsB
K447ATnsB

#### Supplementary Fig. 8 | Transposon right end DNA binding by *Pse*TnsAB

(**a**) Schematic representation of interactions between *Pse*TnsAB and transposon right end (RE) DNA within the paired-end complex (PEC). (**b-c**) Panels show a zoomed view of the structural regions corresponding to the indicated panels in **Fig. 4**, with the atomic model shown in stick representation overlaid with cryo-EM density displayed as a transparent mesh at a contour level of 0.05 for (**b**) and 0.06 for (**c**). Residues highlighted in the corresponding panels of **Fig. 4** are labeled for reference, with consistent coloring. Panel correspondence: (**b**) **Fig. 4a**; (**c**) **Fig. 4d**. (**d**) Analysis of *Pse*TnsB-DNA interactions by electrophoretic mobility shift assays (EMSA). Each panel shows a representative EMSA for indicated DNA. Left, RE BS1 DNA comprising the native BS1 (TnsB-binding site 1) and terminal repeat (TR) sequences of the transposon right end (RE); right, scrambled DNA sequence. Cy5-labeled DNAs were incubated with increasing concentrations of TnsB. Protein-DNA complexes are marked on the side of each native PAGE gel. -, no-protein control; WT, wild-type. (**e**) Structural comparison of TR recognition in the type I-B2 *Pmc*TnsAB[2] and the type V-K *Sh*TnsB[4] strand-transfer complexes. DBD, DNA binding domain; CAT, catalytic domain. TS, transferred strand; NTS, non-transferred strand. Dashed lines represent interaction distances ≤ 3.5 Å. (**f**) SDS-PAGE of purified mutant *Pse*TnsB and *Pse*TnsAB proteins with substitutions in the DNA binding interface of TnsB. (**g**) *In vitro* excision assay with *Pse*TnsAB variants carrying mutations in TR binding residues of TnsB. Left: Representative agarose gel of resolved DNA species; -, no-protein control. Right: quantification of excision from three independent experiments. Relative amounts of plasmid backbone (~2 kb-long excision product) were determined by densitometry. Data represent mean ± s.d. (see **Methods** for details). M, marker.

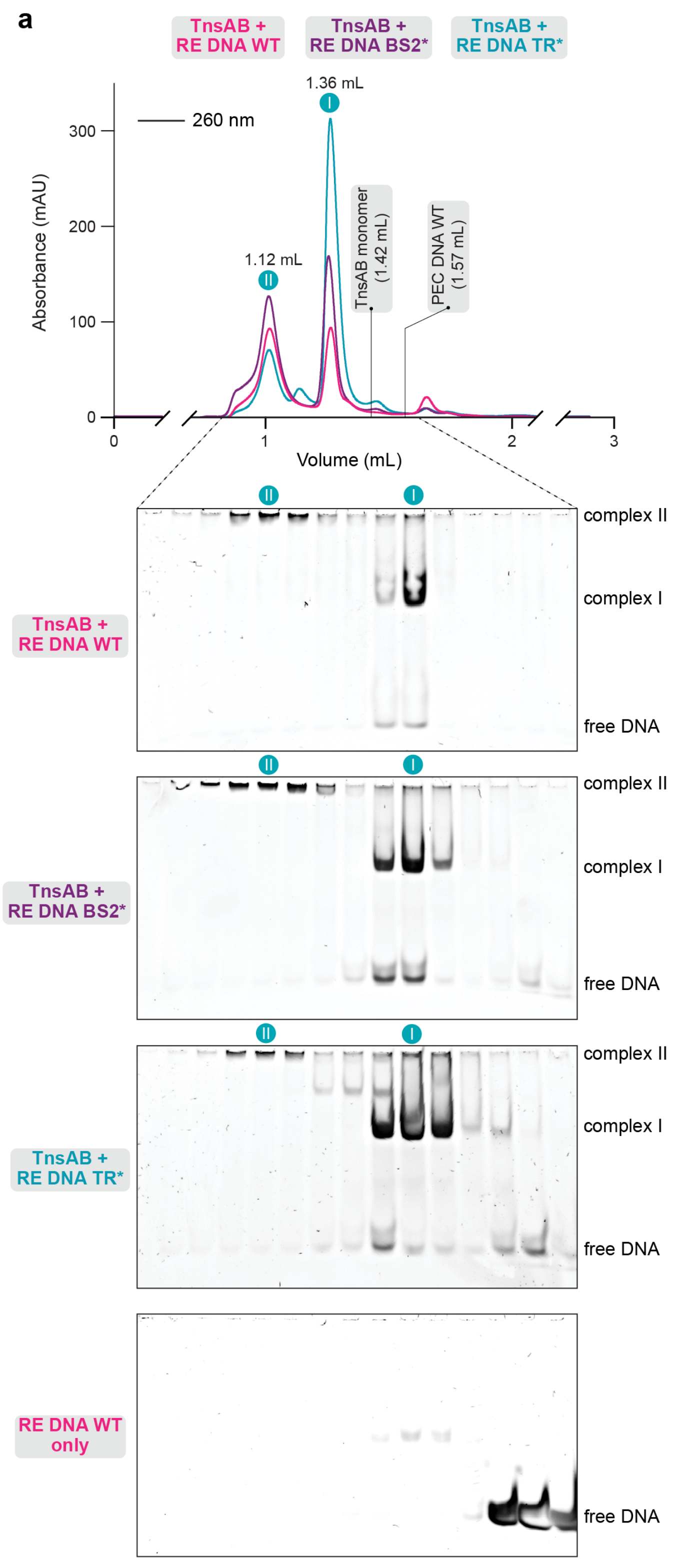


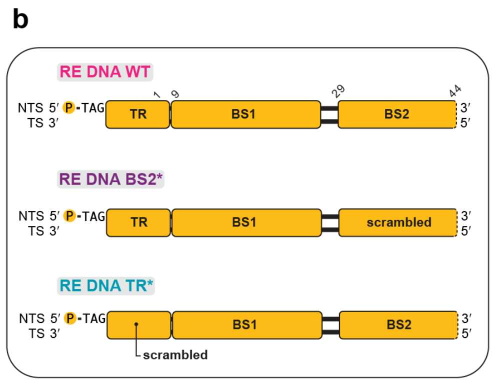


c

| TnsAB + RE DNA | mAU of peak II | mAU of peak I | Ratio mAU II / mAU I |
|---|---|---|---|
| WT | 93.1 | 94.2 | 0.96 |
| BS2* | 127.0 | 169.1 | 0.75 |
| TR* | 70.9 | 312.9 | 0.23 |

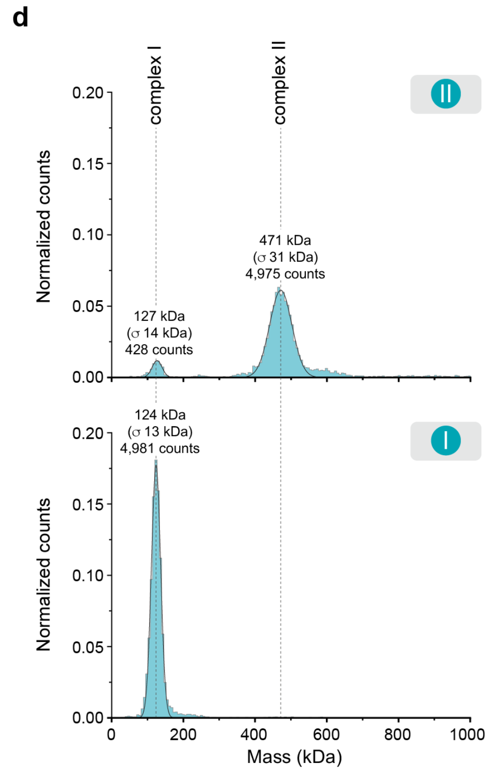


#### Supplementary Fig. 9 | DNA sequence requirements for *Pse*TnsAB complex assembly

(**a**) Size-exclusion chromatography (SEC) profiles of reconstituted *Pse*TnsAB-DNA complexes and analysis of elution fractions by native PAGE. Elution volumes for protein-only and DNA-only controls are indicated on the chromatogram for reference. (**b**) Schematic of the right end (RE) DNAs used, indicating the TnsB binding sites (BS1 and BS2) and the terminal repeat (TR) sequences of the transposon right end (RE). Scrambled DNA sequences are highlighted. NTS, non-transferred strand;

TS, transferred strand. (**c**) Table summarizing SEC analysis of complex I and complex II formation, based on absorbance at 260 nm. (**d**) Mass photometry analysis of PseTnsAB complex II (top) and complex I (bottom), with measured masses and corresponding particle population distributions indicated. The theoretical molecular weight of a TnsAB monomer bound to a single RE DNA is 125.8 kDa, and that of a TnsAB tetramer bound to two RE DNA substrates is 447.0 kDa.

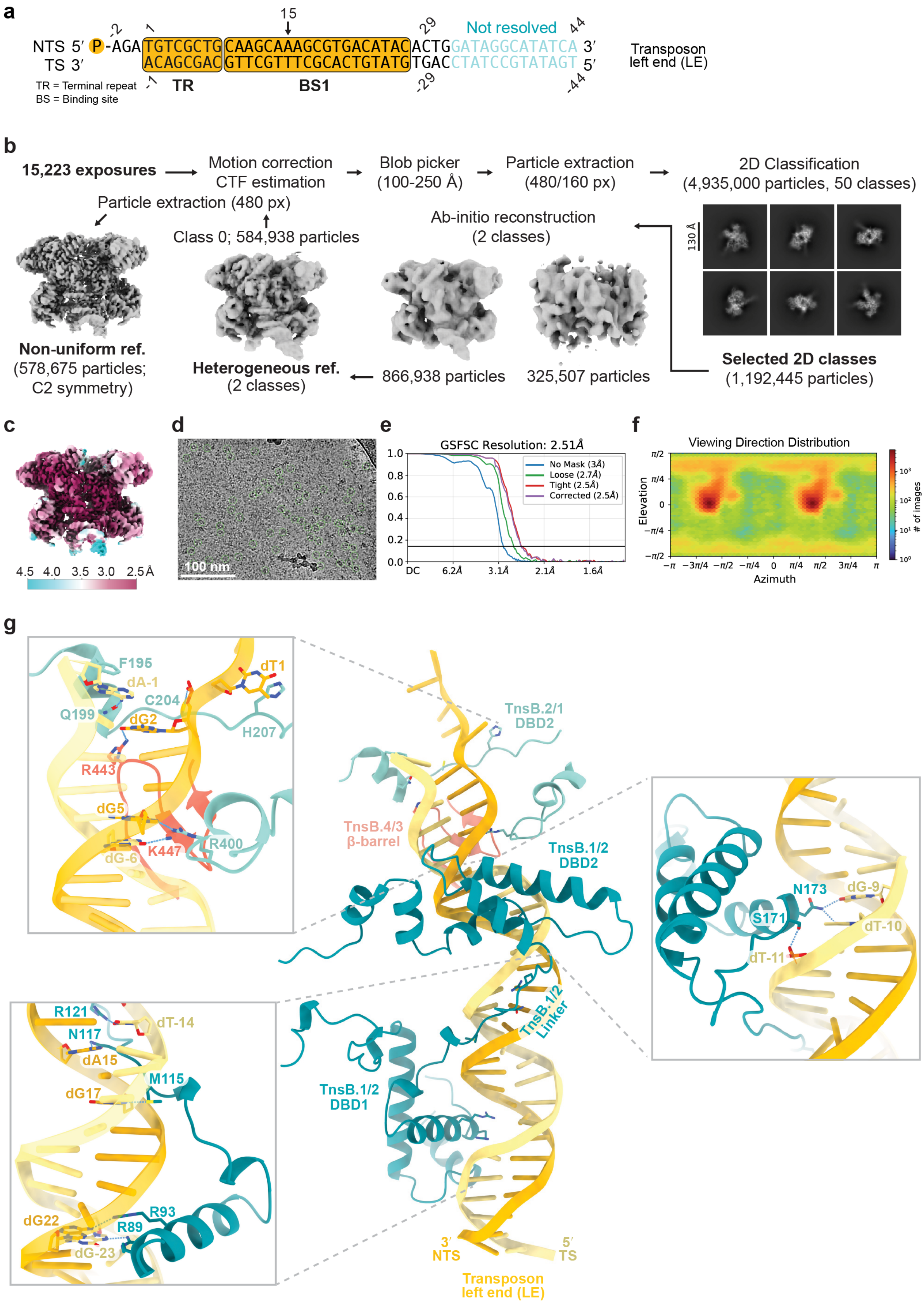
a
NTS 5′ P-AGA TGTCGCTG CAAGCAAAGCGTGACATAC ACTG GATAGGCATATCA 3′
TS 3′ ACAGCGAC GTTCGTTTCGCACTGTATG TGAC CTATCCGTATAGT 5′
Not resolved
-2
1
15
29
44
-1
-29
-44
TR
BS1
TR = Terminal repeat
BS = Binding site
Transposon left end (LE)
b
15,223 exposures
Motion correction
CTF estimation
Blob picker
(100-250 Å)
Particle extraction
(480/160 px)
2D Classification
(4,935,000 particles, 50 classes)
Particle extraction (480 px)
Class 0; 584,938 particles
Ab-initio reconstruction
(2 classes)
130 Å
Non-uniform ref.
(578,675 particles;
C2 symmetry)
Heterogeneous ref.
(2 classes)
866,938 particles
325,507 particles
Selected 2D classes
(1,192,445 particles)
c
4.5 4.0 3.5 3.0 2.5Å
d
100 nm
e
GSFSC Resolution: 2.51Å
No Mask (3Å)
Loose (2.7Å)
Tight (2.5Å)
Corrected (2.5Å)
1.0
0.8
0.6
0.4
0.2
0.0
DC
6.2Å
3.1Å
2.1Å
1.6Å
f
Viewing Direction Distribution
Elevation
Azimuth
### of images
g
F195
dA-1
dT1
C204
Q199
dG2
H207
R443
dG5
R400
K447
dG-6
TnsB.2/1
DBD2
TnsB.4/3
β-barrel
TnsB.1/2
DBD2
N173
dG-9
S171
dT-10
dT-11
R121
dT-14
N117
dA15
M115
dG17
TnsB.1/2
Linker
TnsB.1/2
DBD1
dG22
R93
R89
dG-23
3′
NTS
5′
TS
Transposon
left end (LE)

**Supplementary Fig. 10 | Cryo-EM analysis of the *Pse*TnsAB PEC assembled on left end DNA in the presence of $Mg^{2+}$**

(**a**) Schematic of the transposon left end (LE) DNA substrate used for paired-end complex (PEC) reconstitution, showing the TnsB-binding site BS1, and the terminal repeat (TR) sequences. NTS, non-transferred strand. TS, transferred strand. A15 is indicated with an arrow. (**b**) Cryo-EM image processing workflow. (**c**) Cryo-EM density maps colored by local resolution. (**d**) Representative cryo-EM micrograph. Scale bar, 100 nm. (**e**) Fourier Shell Correlation (FSC) of the reconstruction calculated from two independently refined half-maps. The gold-standard cutoff (FSC = 0.143) is marked with a horizontal black line. (**f**) Angular distribution showing the range of observed particle orientations. (**g**) Detailed views of sequence-specific protein-DNA interactions mediated by the DBD2 (top), linker (middle) and DBD1 (bottom) of the TnsB catalytic protomers on the PEC LE DNA. Dashed lines represent interaction distances ≤ 3.5 Å.

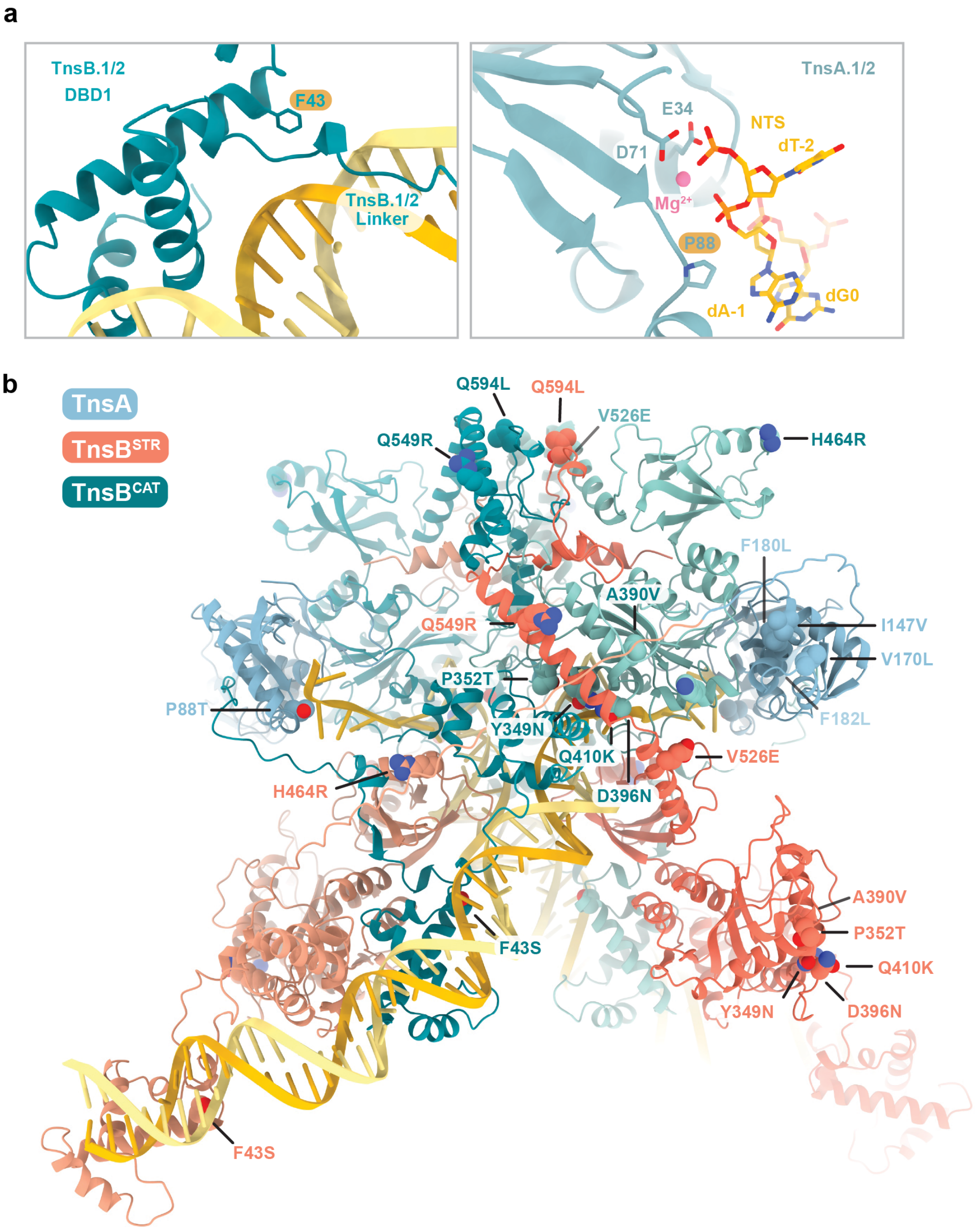


**Supplementary Fig. 11 | *In silico* mapping of amino acid substitutions in evolved *Pse*TnsAB**

(**a**) Detailed view of selected residues mutated in evolved (evo)TnsAB mapped onto the *Pse*TnsAB paired-end complex (PEC) structure. The atomic model corresponds to the PEC assembled on right end DNA in the presence of $Mg^{2+}$. Phe43$^{TnsB}$ and Pro88$^{TnsA}$ are highlighted in orange. NTS, non-transferred strand. (**b**) AlphaFold3[5] model of evoTnsAB PEC assembled on the same right end DNA substrate used for experimental structural determination. TnsA, structural (TnsB$^{STR}$) and catalytic TnsB (TnsB$^{CAT}$) protomers are highlighted in distinct colours (see legend). Residues mutated in evoTnsAB relative to TnsAB WT are shown as spheres.

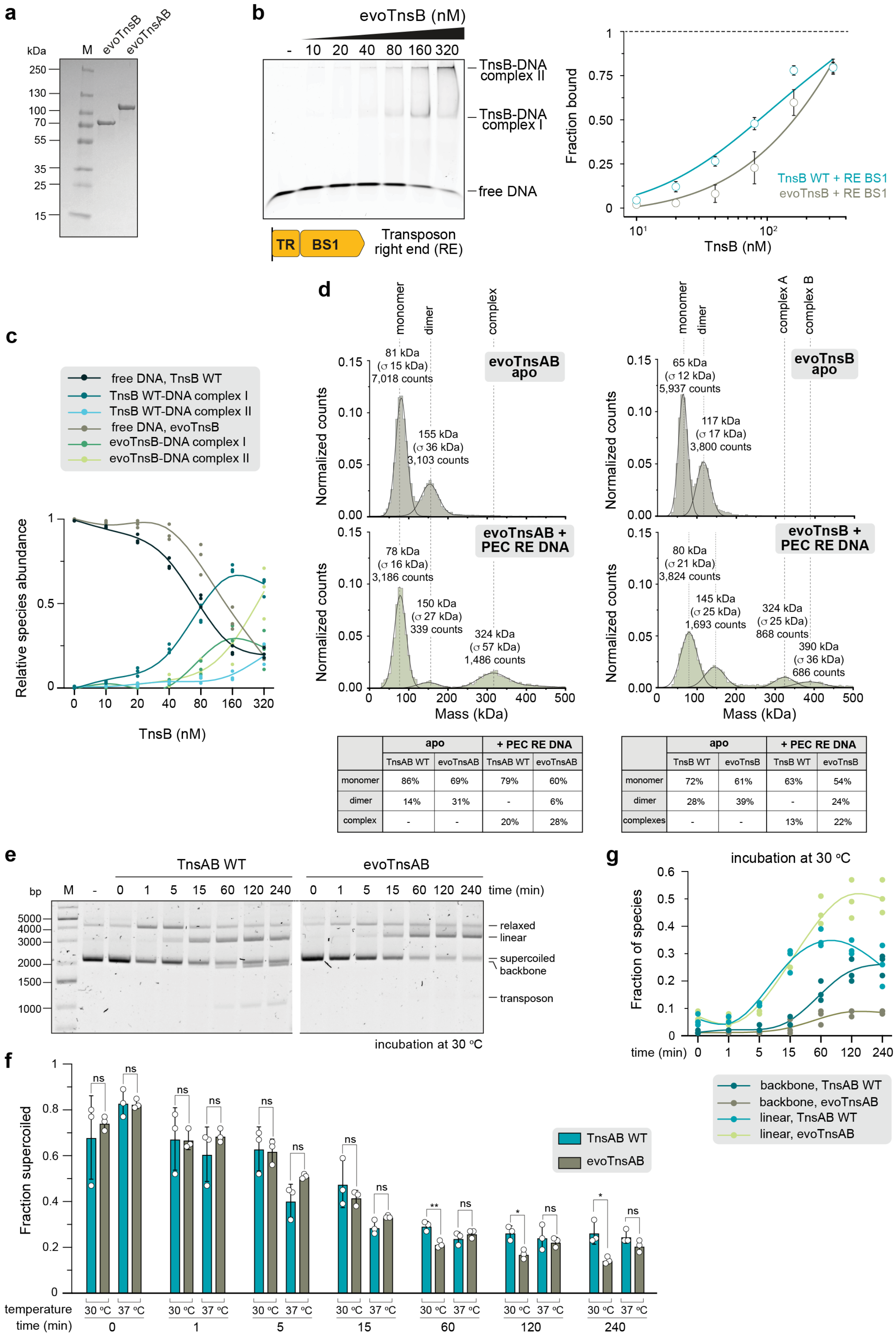


| | apo | | + PEC RE DNA | |
|---|---|---|---|---|
| | TnsAB WT | evoTnsAB | TnsAB WT | evoTnsAB |
| monomer | 86% | 69% | 79% | 60% |
| dimer | 14% | 31% | - | 6% |
| complex | - | - | 20% | 28% |

| | apo | | + PEC RE DNA | |
|---|---|---|---|---|
| | TnsB WT | evoTnsB | TnsB WT | evoTnsB |
| monomer | 72% | 61% | 63% | 54% |
| dimer | 28% | 39% | - | 24% |
| complexes | - | - | 13% | 22% |

#### Supplementary Fig. 12 | *In vitro* analysis of evolved *Pse*TnsAB

(**a**) SDS-PAGE analysis of purified evoTnsAB and evoTnsB proteins. M, marker. (**b**) EMSA analysis of evoTnsB-DNA interactions compared with wild-type (WT). The Cy5-labeled substrate was RE BS1 DNA containing the native BS1 (TnsB-binding site 1) and terminal repeat (TR) sequences of the transposon right end (RE), as in **Supplementary Fig. 8d**. Left: representative native PAGE gel for evoTnsB, with protein-DNA complexes indicated. -, no-protein control. Right: densitometric quantification showing the fraction of bound DNA versus protein concentration (mean ± s.d., $n$ = 3). (**c**) Quantification of relative species abundances from three independent EMSA experiments. (**d**) Mass photometry of evoTnsAB and evoTnsB in apo and DNA-bound states using transposon right end DNA (PEC RE DNA), with measured masses and particle population counts. A comparison with parameters for WT protein is provided in the accompanying table. (**e-g**) *In vitro* time-course excision assays of WT and evo*Pse*TnsAB variants at different temperatures. (**e**) Representative agarose gel showing resolved DNA species for assays performed at 30 °C. (**f-g**) Densitometric quantification of supercoiled plasmid (mean ± s.d.) (**f**), plasmid backbone (~2 kb-long excision product) and linearized plasmid (~3 kb-long single-end cleaved product) (**g**) from three independent experiments. Statistical analysis was conducted using unpaired two-tailed $t$-tests. P-values: *$p < 0.05$; **$p < 0.01$; ns, not significant (see **Methods** for details).

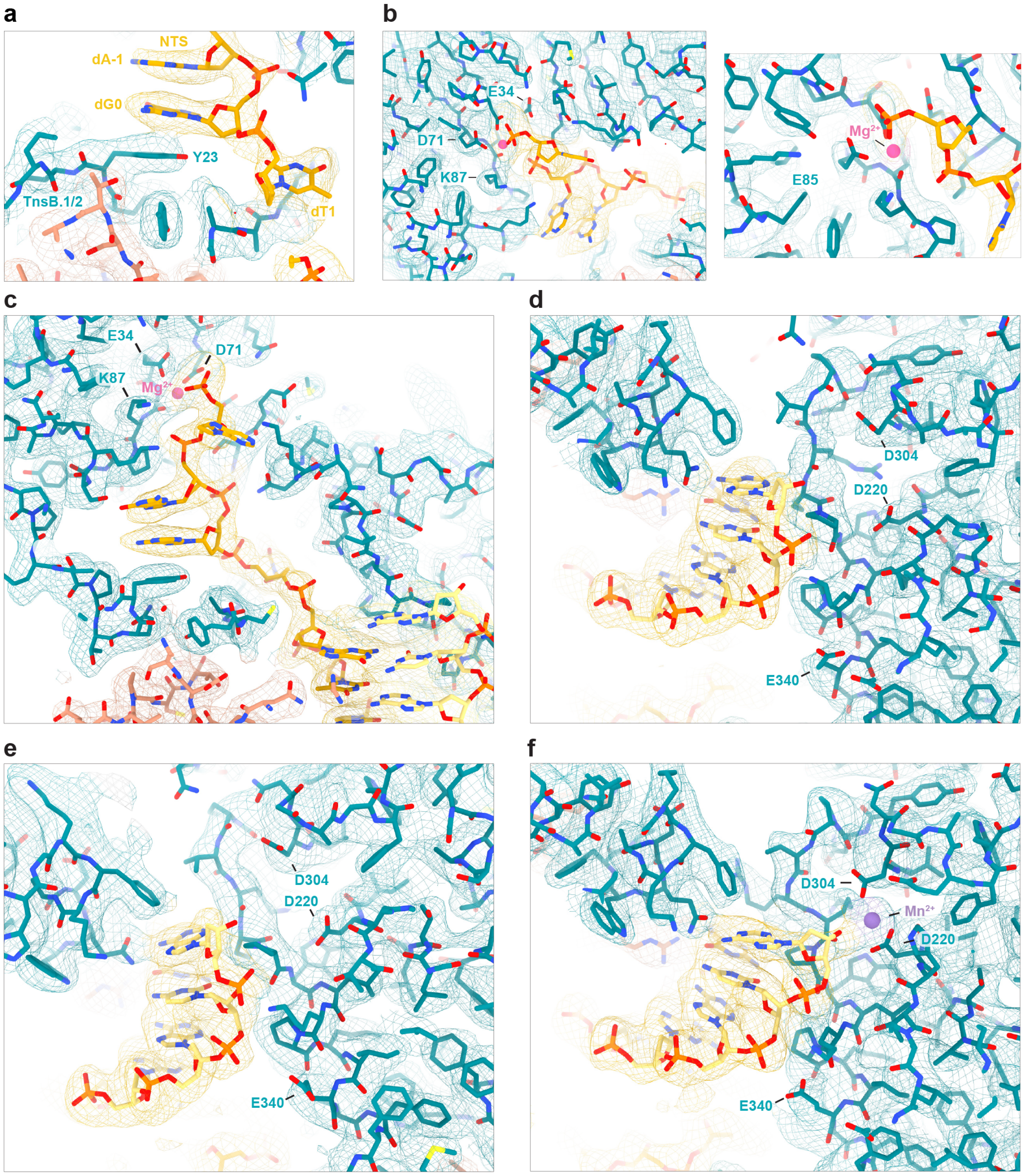
a
NTS
dA-1
dG0
Y23
TnsB.1/2
dT1
b
E34
D71
K87
Mg2+
E85
c
E34
D71
K87
Mg2+
d
D304
D220
E340
e
D304
D220
E340
f
D304
Mn2+
D220
E340

#### Supplementary Fig. 13 | Detailed views of the TnsA and TnsB active sites

(**a**) Detailed view of the TnsB N-terminal tail domain stabilizing the 3-nucleotide 5′ NTS (non-transferred strand) overhang (TAG) accommodated in the TnsA active site. The atomic model is shown in stick representation overlaid with cryo-EM density displayed as a transparent mesh at a contour level of 0.05. (**b-c**) Zoomed views of the TnsA active site in PEC structures assembled on right end DNA (**b**, corresponding to **Fig. 5b**) and left end DNA (**c**) in the presence of $Mg^{2+}$. Atomic models are shown in stick representation overlaid with cryo-EM density displayed as a transparent mesh at contour levels of 0.04 (**b**, left panel and **c**) and 0.06 (**b**, right panel). (**d-e**) Zoomed views of the TnsB active site in PEC structures assembled on right end DNA (**d**, corresponding to **Fig. 5c**) and left end DNA (**e**) in the presence of $Mg^{2+}$. Atomic models are shown in stick representation overlaid with cryo-EM density displayed as a transparent mesh at contour levels of 0.05 (**d**) and 0.04 (**e**). (**f**) Zoomed view of the TnsB active site in the PEC structure assembled on right end DNA in the presence of $Mn^{2+}$ (corresponding to **Fig. 5d**). The atomic model is shown in stick representation overlaid with cryo-EM density displayed as a transparent mesh at a contour level of 0.05. Residues highlighted in the corresponding panels of Fig. 5 are labeled for reference, with consistent coloring.

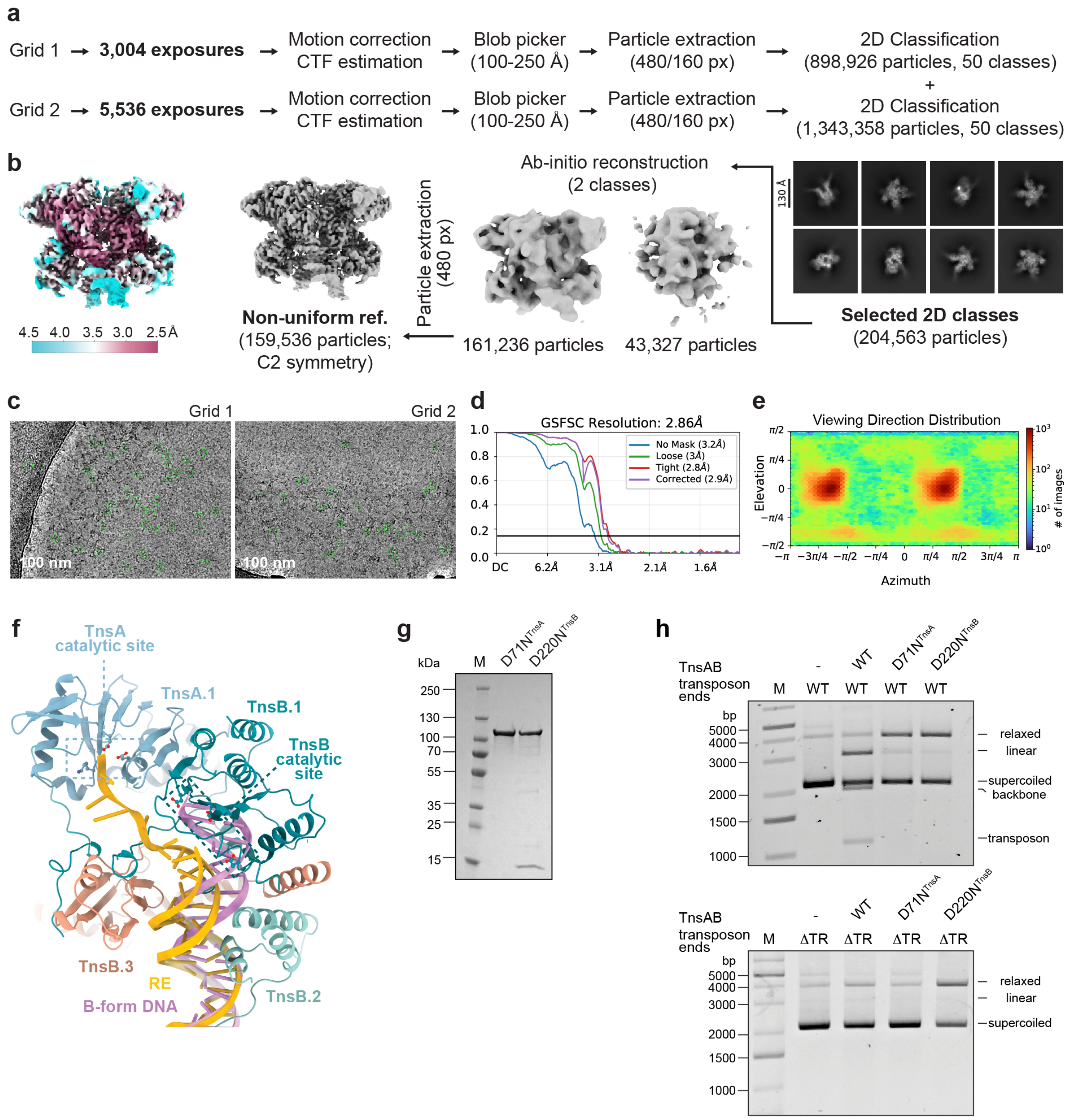


#### Supplementary Fig. 14 | DNA cleavage by *Pse*TnsAB

(**a**) Cryo-EM image processing workflow for the *Pse*TnsAB paired-end complex (PEC) assembled on transposon right end DNA in the presence of $Mn^{2+}$. (**b**) Cryo-EM density maps colored by local resolution. (**c**) Representative cryo-EM micrographs. Scale bars, 100 nm. (**d**) Fourier Shell Correlation (FSC) of the reconstruction calculated from two independently refined half-maps. The gold-standard cutoff (FSC = 0.143) is marked with a horizontal black line. (**e**) Angular distribution showing the range of observed particle orientations. (**f**) Canonical B-form DNA cannot be accommodated in both TnsA and TnsB active sites without steric clashes. The atomic model corresponds to the PEC assembled on right end DNA in the presence of $Mg^{2+}$. (**g**) SDS-PAGE analysis of purified *Pse*TnsAB proteins with mutations in the TnsA or TnsB subunit. (**h**) *In vitro* excision assay with *Pse*TnsAB wild-type (WT) protein and catalytic site mutants. Representative agarose gels of reactions with plasmids containing either wild-type transposon ends (top) or lacking terminal repeat (TR) sequences on both LE and RE (bottom); -, no-protein control; M, marker.

### Supplementary Data

```
Tn7016           1  --------------------------------MT------DFFNEFDESL---VPLKPQ---------TP   20
Tn6677           1  --------------------------------MA----KKGFSSFHRKAVS--SQDTLE----------S   22
Tn6900           1  --------------------------------MD-KHNGGLFEDEFVIPQ---PSTSTS----------P   24
Tn7000           1  --------------------------------MP----NKSFSSFHRKSAF--QQEKLE----------P   22
Tn7001           1  --------------------------------MS----RTSFSSFHGTPTS--DSENAS----------S   22
Tn7002           1  --------------------------------MP-SEGSKRFGELFPSTNF-DNESKPK----------E   26
Tn7003           1  -------------------------------MSDDSENLYAFGSFFPEKHS--NTSVPK----------T   27
Tn7004           1  -------------------------------MPSDSNNIFGFFDEFEASE---GESQQL----------P   26
Tn7005           1  -------------------------------MPPDSNSIFGFFDEFEASE---EESQLL----------P   26
Tn7006           1  --------------------------------MS-NDALFGFRDESSSFDNDVADDKTL----------P   27
Tn7008           1  --------------------------------MS-----FPFNDEFQEIIE--VSGEQN----------K   21
Tn7011           1  --------------------------------MF---NDGLFDDEFNQPL---PKVETK-----------   21
Tn7012           1  --------------------------------MN-SKSLGLFDDEFQHPI---PQVEAP-----------   23
Tn7013           1  ---------------------------------------MPFDDEFESIN---DDTQAEYDSTSEAKLVR   28
Tn7014           1  --------------------------------MS----FGPFEDEFGSIT---NDVQQQYEASPEAKLSR   31
Tn7015           1  --------------------------------MI------EFKDEFTEST---SVKKPD---------TP   20
Tn7017           1  MRASIEPGMEYRFRNRVYSIVSVGDSEMIVMDRYDEEEIHISKDEFNKCW---IKNEIK----------L   57
Consensus_aa:       .........................................F.sb@.p......p.p.............
Consensus_ss:                                                 hhhhhh

                                    |............... DBD1 .................   ............
Tn7016          21  TQYVK--LDDANLIQRDLDTFSDTFKNQALQRYKLISTIDKKLSRGWTQRNLDPI----LDELFKGGDVV   84
Tn6677          23  IELVSSANCLESVTYQDISAFPETIAVEINFRLSILRFLARKCE-TIVAKSIEPH----RVELQQNYSRK   87
Tn6900          25  IDA------IQAVLPATVDSFPYVLKVEALHRRDYILWVEKNLAGGWTEKNLTPL----LADAALVLPPP   84
Tn7000          23  NDRVVDDNDVDEATYQDISAFPDNIATQITFRLSILRYLASKCE-RIIPKTIEPH----RVGLQRLHDRK   87
Tn7001          23  KLTFIE-NDDL----QDLTSFSKKAQNEAKLRLSILRNILNKSS-EINLVLIERL----RADLQQQGERS   82
Tn7002          27  TLSLPL-WKDVDLTFKDVFSVDEDRRNEAIHRYNILEFLIDQYGTDFSKKQIDES----LTALIDKFGSS   91
Tn7003          28  SKGTRF-GIELQESYQDLFSFDEKRRDEAIHRYNILDYLIELHGPSLTLKKISGS----MKGLADKFHPN   92
Tn7004          27  IEL----IQEPIEIFSTIDTQSPKVQQEVIRRIKIIDFVERRLKGGWTEKNLNPI----LSLIEPELKLT   88
Tn7005          27  KEL----ILEPVEISSTIDSLPAKIQEEVLRRIKVITFVEKRLKGGWTEKNLNPI----LSLVESELQLT   88
Tn7006          28  TEF------LAEDENLELAFGDLPHKETALYRLDLIRYLERHVQGGWTPKNLDKL----LEEYALLKKTS   87
Tn7008          22  IIH----NEETNSLLLSLDSYSHNIKMEVLRRISFIKWIKPRLKGGWTQKNLNPL----LDDAATQLKMI   83
Tn7011          22  ---------LPQNYAKDLQALPEKIKNTTFAKLKYIQWLEANIQGGWTQKNLEPL----LKVMPEVEGEK   78
Tn7012          24  ---------KSTKAEKDLSAYPQEIQQTTYARLKYIRWLKERIVGGWTEKNLAPL----INEMPKVEGAD   80
Tn7013          29  KQYLP--LDSVTIHERDLSSFSEEQKNKALERYKLISAVAKEISGGWTPKNINPL----IDKYGLNLSIK   92
Tn7014          32  LKYSP--LETTKVIERDLSSFPEEQKLKALERYKLISLIAKEINGGWTPKNLIPL----IDKHIETLNIP   95
Tn7015          21  GQYIK--LDDAEILKRDLDTFPDFLKEKAFDKYKLISFIEQENSGGWTQKKLDPI----LDKLFEGNRDK   84
Tn7017          58  VSGRMA-NSVKNKLKTPVSTLSEEQKKVAVRKEEYIKKFLDKKDALRKSETDLDYSKIIVEIAELKDPY  126
Consensus_aa:       ...........p.h.pslsshs.p.p.phh.+hphlphl.pp..s.hs.+pl..h....h..h.......
Consensus_ss:       hh         hhh        hhhhhhhhhhhhhhhhhhhhh     hhhhhhh    hhhhhhhhhh

                    ............................| |........ ........ DBD2 ................
Tn7016          85  RPNWRTVARWRKKYIESNGDIASLADKNHKMGNRTNRIK-GDDKFFDKALERFLDAKRPTIATAYQYYKD  153
Tn6677          88  IPSAITIYRWWLAFRKSDYNPISLAPNIKDRGNRETKVSTVVDSIMEQAVERVISGRKVNVSSAYKRVRR  157
Tn6900          85  TPNWRTLARWRKIYIQHGRKLVSLIPKHQAKGNARSRLPPSDELFFEQAVHRYLVGEQPSIASAFQLYSD  154
Tn7000          88  IPSAISIYRWWLVFRASGCNPVSLAPKNKNKGNSKAKVPTFVDALMEQAVDRVISGRKVRIRSAYRRVRR  157
Tn7001          83  IPSAITIYRWWLTYKKSNFSILSLVPKTKQKGNRNTKVPKTISVYIDQAVEQVISANKINVSTAFRRVRR  152
Tn7002          92  TPSAISIYRYWRDFKKSNFQLNSLIPKVSAGNTKRNTPD-ELEPLIQGAIRNYFSEECPTATSAYVTLEV  160
Tn7003          93  VPSAPSIYRYWTTFKKSGFVLSSLIPGVTRGNTKQRKTL-ELEEYIERAIKSYFSAESPTIQQAFTLLET  161
Tn7004          89  PPSWRTLATWKKSYFEAGKDPCALIPKHTFKGNRQKEMD-SQSLIDEAVQNVYLTRERLSVAEAYRYYKS  157
Tn7005          89  PPSWRTVATWKKSYAEAGREASALIPKHTFKGNRQKEMD-SQSLIDEAIQNVYLTRERLSVAEAYRYYKS  157
Tn7006          88  VPSSRTVADWKKLYYESGKDVASLVPGHSKKGNRILKND-SSDLVTEAIQTKFLTKERVSVNTAYEYYKY  156
Tn7008          84  APNWRTLAEWHKNYHSSGNDISSLIPKHSSKGNIQQHTD-SDFLIMKAINEKYLTLTRCSISQTFKYYCD  152
Tn7011          79  KPSWRTAARWYSAYTNADKNIMALIPSHQKKGNRERDTA-TDK-FFEKALERYLVKEKPSVASAYKYYAD  146
Tn7012          81  KPNWRTAARWFKSYSESDESIFALIPKHQKKGKRTQRTD-TDK-FFEKALERFLVKERPSVSSAYQYYKD  148
Tn7013          93  RPSYKSVIRWYKSFCGSDGNIVCLVDHNHSKGNRTKRII-DDESFFVEATERFLDAKRPNYSQAYQFYCD  161
Tn7014          96  KPSDRTVKRWYKAFCESDGDIKSLVDSHHLKGNRQPRIE-DDEPFFIEAVERFLDAVRPSYSKAYQVYCD  164
Tn7015          85  RPNWRTVVRWRKSYIDSNGDLASLVVKRHKMGNRKKRVE-GDEVFFERALSRFLDAKRPKVTTAYQYYKD  153
Tn7017         127  PPSLRTLMRWKSDYNAANGSIRSLINGFSNSGRKSNLNKCQKELIEKAIDNVYLQPEKGTVAQTHSELGV  196
Consensus_aa:       .Psh.olhcW.ps@..ts.sh.tLls..p..Gs+ppc.....p.hhc.h.pp@ls.c+sslspA@p.h..
Consensus_ss:          hhhhhhhhhhhhh     hh                 hhhhhhhhhhhhhh      hhhhhhhhhh

                    .............   ....................................||................
Tn7016         154  LIVIENESIVEGK---IPIISYNAFNKRIKAIPPYAVAVARHGKFKADQWFAYCAAHVPPTRILERVEID  220
Tn6677         158  KVRQYNLTHGTK----YTYPKYESVRKRVKKKTPFELLAAGKGERVAKREFRRMGKKILTSSVLERVEID  223
Tn6900         155  SIRIENLGVVENP---IKTISYMAFYNRIKKLPAYQVMKSRKGSYIADVEFKAIASHKPPSRIMERVEID  221
Tn7000         158  KLRQHNLNNGTK----YKYPTYESLRKRVKKKTPFELLAAKKGERVAKREFRRMGKKILTSYVLERVEID  223
Tn7001         153  KVRQYNLTHKTK----HTYPSYEAIRKRVKKITPYEKLAASKGDRIAKREFRRMGKKIQTEYLLERVEVD  218
Tn7002         161  EIDRHNEINGTQ----FEPFSIQAFRKRLRKHSEYEKILLRKGKSVADSTFRKVGERPATTRVLERVEAD  226
Tn7003         162  EIDRHNECNDTQ-----LSFEYESFRKRIVKKTDYERLLIKKGKKAADTYYKKVGQRPETTRVLQRVEAD  226
Tn7004         158  RVIQINRGIVEGK---IKPISDRSFYNRIHDLPPYEVAIARFGKRYADREFRSVGQQVAATKPMEFVEID  224
Tn7005         158  RVIQMNRGIVEGK---IKPIAERSFYNRINELPPYEVAIARFGKRYADREYRSVGQQVVATKPMEFVEID  224
Tn7006         157  RVIEENRQLTQIK---IATISQRTFYNRVNALPPYEVALARYGKRYADNKFKTVGSIIPATRPMECVEID  223
Tn7008         153  LVIIENRSLPEKK---IKSVSQRTFYNRINALPKYDVTLKQYGKRYADINYRTVDRMVNVTRPLERVEID  219
Tn7011         147  LVIIENDSVVGSV---LKPLTYKAFKNRIDNLPQYDVMVSRYGKRLADIAFNKVEGHTRPTRVLEKVEID  213
Tn7012         149  LVVIENEKVIGED---LRPLTYKAFKNRIDKLPHYEVLRARYGKRIADLAYNKVESHKRPTRVLERVEVD  215
Tn7013         162  RIEIENSNIISGQ---ISKVSYQAFKERLKKLPPYEVALKRFGQNYANKLFNYYQSSVPTTRILERVELD  228
Tn7014         165  RIEIENSTIVSGK---IAKVSYEAFKKRLKKLPPYTVALKRHGKYYADKLFNYYEAVKMPTRILERVEID  231
Tn7015         154  AITIENETIVDGE---IPIISYTAFNQRIKSLPPYPIAVARHGKFKADQWFAYCSSHIPPTRILERVEID  220
Tn7017         197  LVEIHNRENKDNESMIIKCPSRFQVQKAIENLDYLTKLEKREGKREAARDVRTSKHNYRPERILEIGETD  266
Consensus_aa:       .l.b.N....p.....h...obpsh.pRlpphs.@..so+.hE.VEhD
Consensus_ss:       hhhhhhhhhhh          hhhhhhhhhh     eee                          eeeee
```

```
                                          RNAseH-like fold (CAT)
Tn7016          221  HTPLDLILLDDELLIPIGRPYLTLLIDVFSGCVLGFHLSYKSPSYVSAAKAITHAIKPKSLDA-LNIELQ  289
Tn6677          224  HTVVDLFAVHEEYRIPLGRPWLTQLVDCYSKAVIGFYLGFEPPSYVSVSLALKNAIQRKDDLISSYESIE  293
Tn6900          222  HTPLDLLLLDDDLLVPLGRPSLTLLIDAYSHCVVGFNLNFNQPSYESVRNALLSSISKKDYVKNKYPSIE  291
Tn7000          224  HTVVDLFAVHEEHRVPLGRPWLTQLVDCYSKAVIGFYLGFEPPSYVSVSLALKNAILRKDDLLSSFDSVE  293
Tn7001          219  HTWLDLFAVHEEHRVPIGRPYLTQLVDCHSKSVIGFYLGFEPPSYMSVALALKNAIKRKDNLLQNYPSIQ  288
Tn7002          227  HTRLDLFIIDEKNGIPLGRPWLTVLYDTHTKSIVGFYLGFEPPSYLSVSLALVNAILPKEHVKELYPSVE  296
Tn7003          227  HTRLDLFVIDDARKLPLGRPWLTLLFDTHTKSVVGFYLGFEPPGYLSVSLALENAILPKYYVKELYPEVK  296
Tn7004          225  HTPVPVILIDDELEIPLGRPYLTMLYDRFSKCIVGCSINFREPSFDSVRKALLNSLLDKSWVKQRYPSID  294
Tn7005          225  HTPVPVILIDDELDIPLGRPYLTMLYDRFSKCIVGCSINFREPSFDSVRKALLNSLLDKSWLKAKYPSIE  294
Tn7006          224  HTTAPVILLDDDLDLPLGRPHLTILYDRYSTCIVGLSVNYRDPSYETVRAAFLNSVLKKDWVKEKYPSIK  293
Tn7008          220  HTPLDLILLDDNLEIPLGRPYLTILIDSYSKCIVGYNLSFRPPSFESIRHAFCNACLDKSSITQQYPHLQ  289
Tn7011          214  HTPLDLILLDDELHIPLGRPTLTMLVDVYSHCIVGFYFSFSEPSYDAVRRAMLNAMKPKSDVAKLYPDTI  283
Tn7012          216  HTPLDLILLDDELQIPLGRPTLTILIDVYSHCIVGFYLGFNDPSYDAIRRAMLNAMKPKNWIKEQYPDIT  285
Tn7013          229  HTPLDLILLDDDLLIPLGRAYLTLLVDVFSGCIVGFHLGFNPPSYVSVAKAIIHSVKSKDYVHDLNIELT  298
Tn7014          232  HTPLDLILLDDELLVPLGRAYLTLLVDVFSGCIIGFHLGFKAPSYTAVSKAIIHSVKSKEYVNELPIGLS  301
Tn7015          221  HTPLDLILLDDELQLPLGRPYLTLIVDVFSNCVLGFHLSYKAPSYVSAAKAIVHAIKPKTLGI-VGIELQ  289
Tn7017          267  HTLLDYYVIDDDLLMPLGRPWMTNIMELYSRIPLGSEITYENGSFITIANALKMSILPKSNFYEMYPDIK  336
Consensus_aa:        HT.lDlhllD-ch.lPLGRP@LTbLhD.@Sptl lGh.ls@p.PS@.ol..Ah.ptlb.Ks.h.p...plp
Consensus_ss:        eeeeeeeee          eeeeeeee    eeeeeee             hhhhhhhhhhhhhhhhhhhhhhhhhh

Tn7016          290  NDWPCFGKFENLVVDNGAEFWSKNLEHACQSAGINIQYNPVRKPWLKPFIERFFGVMNEYFLPELPGKTF  359
Tn6677          294  NEWLCYGIPDLLVTDNGKEFLSKAFDQACESLLINVHQNKVETPDNKPHVERNYGTINTSLLDDLPGKSF  363
Tn6900          292  HEWPCYGKPETLVVDNGVEFWSASLAQSCLELGINIQYNPVRKPWLKPMIERMFGIINRKLLEPIPGKTF  361
Tn7000          294  NEWLCYGIPDLLVTDNGKEFLSKAFDKACESLLINVHQNRVETPDDKPHVERSYGTINTSLLDDLPGKAF  363
Tn7001          289  NEWLCYGIPDLLVTDNGKEFLSKDFTLACDSLLINIHQNRVETPDNKPHTERQYGTTNTELLNDLPGKTF  358
Tn7002          297  GCWPCYGLPEHLIVDNGTEFNSSDFKIACKDLKIKVKKNPTKKPWLKGSVERYFRTINNRFLSRMPGKSF  366
Tn7003          297  GEWPCYGLPEHLIVDNGAEFNSKDFVTACKNLRIKVKKNPVKKPWLKGSVERYFRTINNKLLSGIPGKSF  366
Tn7004          295  NEWPCHGKIDCLVVDNGAEFWSKSLEDSLRPLVSDIQYSQAAKPWRKSGIEKLFDQMNKGLVNALPGKTF  364
Tn7005          295  NEWPCHGKIDCLVVDNGAEFWSQSLEDSLRPLVSDIQYSQAAKPWRKSGIEKLFDQMNKGLVNALPGKTF  364
Tn7006          294  SEWPCYGNITNLIVDNGAEFWSDSLESALKPLVTDIQYNQRGKPWRKAGVEKSFDTFYKKLFSRFPGKTF  363
Tn7008          290  QDWPISGKIENLVVDNGAEFWSNSLEDSLRPFATNILFNKVGKPWMKPLVEKFFDVLNKGLVHSLPGTTR  359
Tn7011          284  NEWKCAGKIETLVVDNGAEFWSNSLELACEEIGINTQYNPVAKPWLKPFVERMFGTINTELLDPVPGKTF  353
Tn7012          286  HDWHCCGKIETLVVDNGVEFWSNSFESACAQIGISLQYNPVGKPWLKPFVERIFGVINTQLLDPIPGKTF  355
Tn7013          299  NDWLCHGKMETLVVDNGAEFWSKSLDQACMEAGIHYEYCKVGQPWEKPRVERKFLEIIQGIVGWVPGKTF  368
Tn7014          302  NQWICHGKIENLVVDNGAEFWSKSLDQACIEAGINIIYNKVRKPWLKPFVERKFGELIQGIVGWVPGRTF  371
Tn7015          290  NDWPCYGKFETLVVDNGAEFWSKSLDHACKEAGINIQYNPVRKPWLKPFVERFFGMINQYFLTELPGKTF  359
Tn7017          337  NTTNYSGILESLRTDNGADFISNDLDDAALEIGMILHHVQVTKSWQKGAIEGFFRKQNVMLLDKIKGKTF  406
Consensus_aa:        spW.ChGb.-.LlhDNGhEFhSpshcpth..h.hplpbs.h.pPW.Ks.lE+bF..hNp.lls.lPGKoF
Consensus_ss:        hhhh      eeee       hhhhhhhhhh   eeee          ee   hhhhhhhhhhhh

Tn7016          360  SNILEKEEYKPEKDAIMRFSTFVEEFHRWIADVYHQD---SNSRETRIPIKRWQQGFDAYPPLTMNEEEE  426
Tn6677          364  SQYLQREGYDSVGEATLTLNEIREIYLIWLVDIYHKK---PNQRGTNCPNVAWKKGCQEWEP-EEFSGSK  429
Tn6900          362  SNIQEKGDYDPQKDAVMRFSTFLEIFHHWVIDVYHYE---PDSRYRYIPIISWQHGNKDAPPAPIIGDDL  428
Tn7000          364  SQYLQREGYDSVGEATLTLDEIKEIYLIWLVDIYHPK---SNQRGTNCPNVAWKRGCQEWEP-EEFNGSK  429
Tn7001          359  SNYLQREGYNSSDEASLTLSEIKQIYLTWLVDIFHPR---PNSRGTNCPNRVWKHAEKNWPP-NEFLGTD  424
Tn7002          367  TNIFERKDYDPLKNAVIDNSLLQEMVHIWIVDVYQNG---KNGLENNIPNLSWVDAVNSSIPPRPFKGSR  433
Tn7003          367  SNIFARGDYNPQKNAIITRSDLMKVIHVWLIDIYQSS---PNGLENNIPNLSWADAMRSAFPPRSFNGSI  433
Tn7004          365  TNPTQLQDYNPKKDAVVRVSVFLELLHKWIVDYYHMA---PDSRERDIPYHKWNQSEWTPSY--YSGVEK  429
Tn7005          365  TNPTQLQDYNPKKDAVVRVSVFLELLHKWIVDYYHMA---PDSREREIPYHKWHQSKWTPSY--YDGAEK  429
Tn7006          364  TNPTQLKDYNPKRDAVINVSDFLELLHKWLIDVYHKK---ADTRYKRVPYQKWTESQGKIIF--CEGPEA  428
Tn7008          360  SRIEQLKGYNPKKDAAITFSLFLELFHTWIIDIYHMS---SDTRETAVPYFKWQEGVTALPPLTYTDEEA  426
Tn7011          354  SNILQKHEYNPKKDAIMRFTTFMQLFHKWVVDVYHQD---ADSRFKYIPSQLWEQGFNTLPPTVLSNADL  420
Tn7012          356  SNMMKKEGYDPNKDAIMRFSAFMEVFHHWIIDVYHQE---ADSRFSYIPALEWDRGYEQLPPAPLSSDDI  422
Tn7013          369  SNILEKDRYDPQKDAVMRFSSFVEELHRWIIDVHNAS---PDSRNTKIPNYHWKKSEEALPPAALSDRDE  435
Tn7014          372  SNVLEKEDYDPQKDAVMRFSVFVEELHRWIIDVHNAS---ADSRHTRIPNYHWQKSEEVLPPPALTERDE  438
Tn7015          360  SNILEKEDYKPEKDAIMRFSVFVEEFHRWIVDIYHQD---SDSRDTRIPIKQWQHGFDVYPPLQMSVEDE  426
Tn7017          407  SNILERAKYDPKKQSLIRLSDFKNILYKWLYDVYPYNEVRVEHGKKIIPHKLWEKNIKDHPQ---DFIEK  473
Consensus_aa:        oNhhp...YsPp+pAhhphS.h.pbh@.WllDlYp.p...ssp..p.lP...Wpct.p.h........p.
Consensus_ss:                  hhhhhh  hhhhhhhhhhhhhhhh              hhhhhhhhhhh          h

                                                                      β-barrel
Tn7016          427  TRFSMLMRISDSRTLTRNGFKYQE-LMYDSTALADYRKHYPQTK-ETVKKLIKVDPDDISKIYVYLEELE  494
Tn6677          430  DELDFKFAIVDYKQLTKVGITVYKELSYSNDRLAEYRGKKG-----NHKVQFKYNPECMAVIWVLDEDMN  494
Tn6900          429  TKLEVILSLSLHCTHRRGGIQRYH-LRYDSDELASYRMNYPDQTRGKRKVLVKLNPRDISYVYVFLEDLG  497
Tn7000          430  DELDFKFAIVDQKQLTKAGVTVYKELTYSSERLAEYRGKKG-----NHKVQFKYNPECMAVIWVLDEDLN  494
Tn7001          425  EELDFYFTKLDKRMLRKEGIGLATELFYSSERLAEYRGKKG-----NHKVAIKYNRENMGFIWVLDEDTE  489
Tn7002          434  DELKFNLGKNREVSLDKNGVRLGTTIRYSSVRLARYFGHKTCKDMKSVRVRIKYDPSCLGRVYVLDETKE  503
Tn7003          434  DELRFNLGKHVEISLDRNGIRLKKTLRYTSSYLAQYFGKHTYDG-KSIKVKIKYNPICMGNIYVLDEDKH  502
Tn7004          430  EQLRVELGLLRHRTIGVAGIRLHN-LHYQSSELIEYRKYWASNNGKKLFVKTKTDPSDISSIHVYLESEK  498
Tn7005          430  EQLRVELGLLRHRTIGVAGIRLHN-LRYQSAELIEYRKYCTPNNGKQLFVKTKTDPSDISYIHVYLESEK  498
Tn7006          429  EQLKIELGVVNHRTIRRGAIELHS-LKYQSDELEEYGKQYSSRARKSAYVKTKTDPNDISSIYVYLEEEK  497
Tn7008          427  QQLRIELGILNTRTVRIGGIFLYG-LRYESEELNEYRKIWGAIDKNNLTLKTKTDPSDISNIFVYLTNES  495
Tn7011          421  QQLDVVLSISNHRVLRKGGIRLEN-LSYDSTELANYRKQFSHKV--SQEVLIKLNPDDISYIYVYLDKLE  487
Tn7012          423  QRLEIVLGISTWRHIRKGGIHFEN-LRYDSNELADYRKRFSPKA--SVKLMLKVNPEDLSRVYAFLPELD  489
Tn7013          436  KQFRIIMGVIHEGVVTTKGIKYKH-LMYDNVALEQYRKQYPQTK-KSRKKTIKIDPDDLSSIFVYLEEIG  503
Tn7014          439  IQFRVIMGMVHKGALTSKGIKFKH-LMYDNVALEHYRKQYPQSK-DSRIKTVKIDPDDLSRIFVFLEEKK  506
Tn7015          427  KRFNVLMGITDERTLTRNGFKFEE-LMYDSTALADYRKHYPQTK-DTIKKLIKIDPDDLSNIHVYLEELE  494
Tn7017          474  SNLDIIFGKTGSRKARDKGIEINN-LKYDGIDLTKFRASHGN----NETVKVKFDPSDLGFIHFFDKFEK  538
Consensus_aa:        pphch.htb..p..lpp.Glph.p.L.Ypt..L.pY..p.s.p...s....hKhsPpsht.I@Vh.cpbp
Consensus_ss:        hhhhhh   eeeeee    eee   eeee hhhhhhh           eeeeee      eeeeee
```

```
                       ..............||.......... ................C-helical............... ..........................................................  ..|
Tn7016           495   SYLEVPCTDPT-GYTDGLSIYEHKTIKKINREVIRE-SKDSLGLAKARMAIHERVKQEQEVFI-ESKTK-   560
Tn6677           495   EYFTVNAIDY--EYASRVSLWQHKYNMKYQAELNSA-EYDEDKEIDAEIKIEEIADRSIVKTN-KIRARR   560
Tn6900           498   SYIRVPCIDPI-GYTKGLSLQEHQINVKLHRDFINE-QMDVVSLSKARIYLNDRIKNELIEVR-RNIRQ-   563
Tn7000           495   EYFTVHAIDY--EYASRVSLWQHKYNMKYQAELNSA-EYDEDKEIDAEIKIEEIADRSIVKTK-KIRARR   560
Tn7001           490   SYFKVPAIDY--EYASTVTLWMHKKNLKIKQELNIT-EHNLDSEIDAELRIDEIVDQSIQKKKTTITNKR   556
Tn7002           504   EFFAVDAIDA--DYAYSVSEWLHKTCCNYARRHIRK-NYNHRDVVKAWRKILDIIDEAFEKTGQNKKQNS   570
Tn7003           503   EFFAVESVDP--DYAYSVSEWLHKVCCDYARNHIRN-NYRHNDVIKAWRVIYDIIDEALHLSG-NGKQAN   568
Tn7004           499   RYIKVPAVDNT-GYTKGLSLFEHERIQKVRRLNTKQ-LTDDEALADTFLYMKKRIQEETDRFR-GAKSNK   565
Tn7005           499   KYIKVPAVDNS-GYTNGLSLFEHQRIQKVRRLNTKD-LADDEALADTFLYMKKRIHEETDRFR-RVKSSK   565
Tn7006           498   KYIKVPAVDHT-GYTKGRSLYEHQRINSLRRLKVRL-GEQDESLADASLYLDRAMDEAIERMS-RSKSKK   564
Tn7008           496   RYIKVPCITDI-SYTSGLTLFQHQTAQKLQRTKTRL-QIEHEKLADSRIYVENRIAEEVEKIK-SSKK--   560
Tn7011           488   HYIKVPCIDPN-GYTQNLSLNQHKINIRIHRDFISG-SIDNVGLAKARMFIHNKIQNEFEELK-NAPKH-   553
Tn7012           490   HYIEVLCIDPN-GYTAGLSLHQHKVNVRLQRDFIGS-HMDVAALARARMLIHEKIQNEVELLA-NSTKK-   555
Tn7013           504   GYIEVPCKYDPLGYTKNLSLSEHVRITKIHRDFIKG-QVDALSLAKARQALHERIKTEQEHLS-LMSVES   571
Tn7014           507   GYIEVPCKYDPLGYTKKLSLCEHLRTVKVHRDFIKG-QVDSLSLAKARQALHERIKQEHENLR-QMSLPH   574
Tn7015           495   GYLKVPCTDTT-GYANGLSLHEHKVIKKINREIIRE-SKDNLGLAKARMAIHARVQQEQELFN-ESKTK-   560
Tn7017           539   RFIRLNAVDS--EYANKLTLYQHGIIKKYVKEDMRRSEEDIKVLMDGKIQLRDMINDLLENKP-SKLASK   605
Consensus_aa:          p@hpVsthD....Yhp.loLhbHb...ch.+p.hp..p.s..slhct.b.lcc.lpp.hp....p...p.
Consensus_ss:          eeeeee                hhhhhhhhhhhhhhhhh hh hhhhhhhhhhhhhhhhhhhhhhh hhhhhh

Tn7016           561   ---AKITAVKKQAQIADVSNTGTSTIKVSE------ESA-APVQ-KHISN--DNSDDWD---DDLEAFE-   613
Tn6677           561   RGARHQEN------SARAKSISNA------------NPA-SIQK-HEDEIVSADNDDWDI--DYV-----   603
Tn6900           564   ---RNVKGVNKIAKYRNVGSHAETSIVHELNHPATNEVISKMES-ASQPEHCDDWDNFT---SGLEPY--   624
Tn7000           561   RGARHQEN------SARAKSISDA------------KPV-PSQK-HEDEIVIVDNEDWDI--DYV-----   603
Tn7001           557   RAARYQENEQM---AANARQPKEK------------KSA-NNQKTSTKNIEKEDSSGWDI--DYV-----   603
Tn7002           571   LGTTTTSR------VQRIKEHSKRATSEAKTQQVDYVKS-ELDD-ELDWDIEVNTNGWSI--K-------   623
Tn7003           569   VGIRQASK------LERVREHAERTKSHQKPE----LHM-SSND-DIDWDVEVNTDGWKI--DSVRGTNK   624
Tn7004           566   ASLPKTGNTSKLAKFNDVGSEGPSSINVIPVT----HKS-EVV-----SDASEYLDDDDF--EDIEGY--   621
Tn7005           566   PNLPKTGNTSRLAKFNDVGSEGPNSINVTPVR----LKS-EVV-----SDASEYLDDDDF--EDIEGY--   621
Tn7006           565   SALPKTTHASKIAKQRGVGSEGPSTIVITSPK----PIS-EVP----KEVIDIGTASDDL--SDIEGY--   621
Tn7008           561   -RTTKTTHASKIARHQDIGSHTQKSIQIPNEQ----SDT-KNE--------HDVLNGWDEQFDDLEGF--   614
Tn7011           554   ---SKVKGGKALAKHQNVSSDSQKSIAQSEPL----EPK-KVTP--KEQP-TDSWDDFI---SDLDGF--   607
Tn7012           556   ---PKARGGKALAKYQNVASDNLVSVKPSSPTKKPPEKV-ETEE---AQN-KGDWDDFV---SDLEGF--   612
Tn7013           572   -RAKKAKHGKKMAALSGISNEQPMSIQNALEN----KNK-PLD-----DNFD--EPTPV---DNLKSLWN   625
Tn7014           575   -RAKKAKNGKKMAELAGVNSDSPKSI-----T----TDY-PIE-----DTIQLHESTPV---DDLQSLWN   625
Tn7015           561   ---AKISAVKKQAQLADISNTGQGTIRLENSD----TLS-DITN-KPESNISDILDNWD---DNIEGFE-   617
Tn7017           606   RGAQLRQGMTNS--IADAGENQRDSIPRLEEN----QIS-------SVSDVSVESTACDSIISDSKGT-E   661
Consensus_aa:          ....p.p..p....h.ph.pp...ol.....p..................hp...ss.s...sslcth..
Consensus_ss:           hhhhhhhhhhhhhh

Tn7016                 ------------------
Tn6677                 ------------------
Tn6900                 ------------------
Tn7000                 ------------------
Tn7001                 ------------------
Tn7002                 ------------------
Tn7003                 ------------------
Tn7004                 ------------------
Tn7005                 ------------------
Tn7006                 ------------------
Tn7008                 ------------------
Tn7011                 ------------------
Tn7012                 ------------------
Tn7013           626   KRKAMKRSKE--------   635
Tn7014           626   KRRALRKSGK--------   635
Tn7015                 ------------------
Tn7017           662   ESKKNTSYEKWTDDDIEW   679
Consensus_aa:          ..................
Consensus_ss:
```

#### Supplementary Data 1 | Multiple sequence alignment of TnsB proteins from type I-F CASTs

Output from Promals3D[6]; Tn*7016* (*Pseudoalteromonas* sp. strain S983), WP_029772100.1; Tn*6677* (*Vibrio cholerae* strain HE-45), WP_001074608.1; Tn*6900* (*Aeromonas salmonicida*), WP_088821968.1; Tn*7000* (*Vibrio cholerae* strain 4874), WP_119464978.1; Tn*7001* (*Photobacterium iliopiscarium*), WP_107274795.1; Tn*7002* (*Vibrio* sp. strain F12), WP_136985358.1; Tn*7003* (*Vibrio parahaemolyticus*), WP_015313514.1; Tn*7004* (*Vibrio* sp. strain 16), WP_005472985.1; Tn*7005* (*Vibrio cholerae* strain M1517), WP_001134621.1; Tn*7006* (*Vibrio splendidus*), WP_054541713.1; Tn*7008* (*Aliivibrio* sp.), WP_065591601.1; Tn*7011* (*Pseudalteromonas* sp. strain P1-25), WP_055018246.1; Tn*7012* (*Pseudoalteromonas ruthenica*), WP_138590899.1; Tn*7013* (*Vibrio cholerae* strain OYP7G04), WP_114778930.1; Tn*7014* (*Vibrio diazotrophicus*), WP_102952857.1; Tn*7015* (*Shewanella* sp.), WP_076538387.1; Tn*7017* (*Endozoicomonas ascidiicola*), WP_095210281.1[7]. Domain boundaries of *Pse*TnsB are indicated above the alignment; residues described in this study are highlighted in yellow.

```
Tn7016             1   --------------------MYIRNLRKPSPNKNVFKFASTKVSSVVMCESSLEFDACFHHEYNDLIESF   50
Tn6677             1   MTSLPTPSAITTSALEYAFHTPARNLT-KSRGKNIHRYVSVKMSKRITVESTLECDACYHFDFEPSIVRF   69
Tn6900             1   --------------------MYRRHLK-HSRVKNLFKFVSAKMNTVFTVESALEFDTCFHLEYSPSVKFY   49
Tn7000             1   MSALPSPSTTTLIALESAFVTPARNLT-KSRGKNIHRYVSAKMGKRVTVESFLECVACYHFDFEPSIVRF   69
Tn7001             1   MQPSPATTYQTDSPLEFAFTQPARKLT-KSRGKNIHRYVSIKMNTIISVESTLEFDACFHFDFNKNISRF   69
Tn7002             1   --------------------MKKRILR-NSSVKNISRFVSLKTNSIHTVESDLEFDACFHFEFSPQIITF   49
Tn7003             1   --------------------MKKRIIK-NSKVKNISRFVSLKTDSVQTTESDLEFDACFHFEFASHVKSF   49
Tn7004             1   --------------------MFDQTKK-SSHVHNICKFMSLKNDAVVRTLSILEFDFCFHLEYNADIKTF   49
Tn7005             1   --------------------MFDQTKK-SSHVHNICKFMSLKNDAVVRTLSILEFDFCFHLEYNPNIKSF   49
Tn7006             1   --------------------MYDQTKK-SSAVHNICKFMSLKNDSVVRTMSMLEYDFCFHAEYNPQIVRY   49
Tn7008             1   --------------------MYVRTLK-QSQVKNISKFMSLKNDSIIRTESMLEFDMCFHLEYSPDVVSF   49
Tn7011             1   --------------------MYRRKLK-HSRVKNLHKFASQKNKSTCLVESSLEFDACFHFEFSPSIAAF   49
Tn7012             1   --------------------MYRRNPK-SSRVKHQFKFASQKSKNTCFAESALEFDACFHFEYSSAIVAF   49
Tn7013             1   --------------------MYVRNLRKPSATKNVYKFASSKNRSVILCESSLERDCCYHLEYSKDVFSF   50
Tn7014             1   --------------------MYVRNLRKPSANKNVYKFVSVKNGCNIMCESSLEYDCCYYLEYSDDVVRY   50
Tn7015             1   --------------------MYIRNLRKPSPNKNVFKFASAKVSETIMCESTLEFDACFHHEYNETIETF   50
Tn7017             1   MTPL----------LSQY--VGSRKLT---PGRHRYKYPSRKMEAHVITESPNECNFCEFLEFDSRVKGY   55
Consensus_aa:          ....................h..Rplp..S.s+Nlh+@hS.K.pshh.hES.LEhDhC@HhE@sspl..@
Consensus_ss:                                           eeeeeee      eeeee hhhhhhhhhh      eeee

                       ...............................nuclease...................................
Tn7016            51   GSQPEGFKYEFMGKSLPYTPDALISYTDKTQKYHEYKPYSKIASPLFRAEFAAKRAASLK-LGIDLVLVT  119
Tn6677            70   CAQPIRFLYYLNGQSHSYVPDFLVQFDTNEFVLYEVKSAYAKNKPDFDVEWEAKVKAATE-LGLELELVE  138
Tn6900            50   EAQPEGFYYEFAGRQCPYTPDFRLVDQNDSVSFLEIKPSDKVADPDFLHRFPLKQQRAIE-LSSPLKLVT  118
Tn7000            70   CSQPIRFSYGLNGKTHTYVPDFLVQFDTGEFKLYEVKSDMESSKEEFQCEWEAKVQGAFE-LGLELELVT  138
Tn7001            70   CSQPIRYSYVIDGIIRTYVPDFLCEFYCGEMVLYEVKAPNAVNSKRFTKEFEAKRQYARKLFDVDLELIE  139
Tn7002            50   EAQPIGFEYFIEGKVHRYTPDFLVTYKDSYQSFYEIKPKHIAEKDEFKEKFRAQKEQALG-DGKDLHVLT  118
Tn7003            50   ETQPLGFEYRLNGRLRRYTPDMLCYFNDGYATYYEVKPKWVTERDEFKKKFDAQKQQAIA-NGYDLLVLT  118
Tn7004            50   TSQPFGFHYQFNNRKCRYTPDFLATDHDEHSTFFEVKHSSQILKPDFRERFKEKQRVAFNEFNRRLVLVT  119
Tn7005            50   TSQPFGFHYLFNNRKCRYTPDFLAIGHNEQSTFFEVKHSSQIPKPDFRERFEEKQRVALSEFNRRLVLVT  119
Tn7006            50   ESQPHGFEYYFNGRYCRYTPDFQLFDSIDTPSLIEVKHSSQILKPDFRARFKEKQLVAQAEYGKKLILVT  119
Tn7008            50   ESQPQGFHYEYQGKRLPYTPDFLITHSSGQQQLLEVKPLSKTQRPDFQSKFIQKQQAAQK-LNLSLILIT  118
Tn7011            50   EAQPLGYEYEFDNRICRYTPDFLLTHTDGTQKFIEVKPQSKIADEDFRARFIEKQTIAKQ-DGRDLILVT  118
Tn7012            50   ETQPLGFKYDFEGRTCPYTPDFLLTHSDGTQKYIEIKPVKELAKDEFRQRFEQKQLASKQ-LGIELILVT  118
Tn7013            51   QSQPEGFYYSSGNKRCPYTPDFLVRNQDGSEYYLEVKPLAKTFSEDFKRSFALKRIAAQH-QGKPLVLVT  119
Tn7014            51   QSQPKGYRFPYQGKEHPYTPDFLVHKKDGTSYLLEVKPLSKTFSSEFQDVFRQKQIMASE-LGAPLLLVT  119
Tn7015            51   GSQPKGFYYCFEGKRLPYTPDALLHYIDGTTKFHEYKPYSKTFDPIFRAKFVAKKEAAQA-LGTELILVT  119
Tn7017            56   ISQPKTIIYDLYQCGHDYTADFFVDLYLSKPAFYEIKPKDYEKSEEEEDKFYEVEKVFYD-MGFNFYLVT  124
Consensus_aa:          .tQPbs@.Y.h.sp.h.YTPDFhh...ssp..hhElKs..b..p.-Fp.cF..ppb.tb..bs.pL.LlT
Consensus_ss:          ee   eeeeee  eeeeeeeeeeeeee   eeeeee  hhhh  hhhhhhhhhhhhhhhh     eeeee

                       ........||.......................helical....................      ...................................
Tn7016           120   DRQIRVNPILNNLKLLHRYSGVYGISGIQKELLSFIHKSGVI-----KLNDISSQVGIPIGETRSFLFGL  184
Tn6677           139   ESDIRDTVVLNNLKRMHRYASKDELNNVHNSLLKIIKYNGAQ-----SARCLGEQLGLKGRTVLPILCDL  203
Tn6900           119   EKQIRIAPILGNLKLLHRYSGFQSFTPLHMQLLGLVQKLGRV-----SLLRLSDSIDAPPEEVLASALSL  183
Tn7000           139   EEEILDEVIFSNLKLLHRYASRDNLNHFHQTLLATFKLNGTQ-----TAKSLGHHLGLNGRKILPFLCDL  203
Tn7001           140   EHDIRHIPLLKNLKRMHRYASHSELTYEQTTILKYLNKHGST-----KLETIIDHFSNS-KNIIPMIYDL  203
Tn7002           119   DDDIQIYPLLDNLRIIHRYACDERLNSIQRNILNLFKKYGEL-----RIEQVVQYSSVHSSQLLPSLYDL  183
Tn7003           119   EDDIQTYPLLDNLKIIHRYACSDSLDNVQVRILKLFQNYGEM-----RISQVINASQGQSASILPALYDL  183
Tn7004           120   EKQIRMGPTLDNFKLLHRYSGLRTVTEFQKLVLAFIQRKQMV-----KLQEVSLYFGLSEQDTLISTLPW  184
Tn7005           120   EKQIRMGPTLDNFKLLHRYSGLRTVTEFQKRVLAFIQRKQMV-----KLQEVSLYFGLSEQDTLISTLPW  184
Tn7006           120   EKQIRTGFLLSNLKLLHGYSGIRTITDIQKHVLQFVQANRSV-----TLHYLSHQLKISPDETLTAALCW  184
Tn7008           119   EKQIRTGHLLNNFKLLHRYSGLHSISVTQKAIIHLIQKVNKI-----QINQIANSLNISNGEALTGVLSW  183
Tn7011           119   DKQIRVYPTLNNLKLLHRYSGFQSLTELQASVLELVKQYGSI-----KVGQLVNFLKVTAGELLATVLRL  183
Tn7012           119   DKQIRVFPVLDNLKLLHRYSGFQVLTELHTVVVGLVKSTGLV-----KVAQLVNYLKVSAGEVLSIVARL  183
Tn7013           120   DKQIRNGVYLENLNLIHRYSGLVDFSLSSTKIVEELSTAGRM-----CIRSLADNLKLSIGEVIAVVFRL  184
Tn7014           120   DRQIRNDVHLNNLKLVHRYSGCIGNSSHLESVWSAVNQSSSI-----CIKALSAILNLTIGEVFASVLRL  184
Tn7015           120   DKQIRVNPILNNLKLLHRYSGIYGVTDIQRELLQLIRHSGKI-----QLDDVADEYELSVGETRSFLYSL  184
Tn7017           125   GDYIEKGNRLINFKKLRRFVGSLPERKIREFIERLVFESNEGISFGDLIAKLRNQLDVEERDAFQNIYQC  194
Consensus_aa:          -cpIc..shLpNhKblHRYtt..shs.hpp.ll.hlpp.s........l.pl.p.h.lp..phh..lhph
Consensus_ss:          hhh  hhhhhhhhhhhhhh       hhhhhhhhhhhhh             hhhhhhh     hhhhhhhhhhh

                       .......|
Tn7016           185   MHKGLVKADLGCDDLTNNPTLWATP--------------   209
Tn6677           204   LSRCLLDTRLDK-PLSLESRFELASYG------------   229
Tn6900           184   IARGIMQSDLTVQKIGISSFVWAGGHSGIDHG-------   215
Tn7000           204   LSRNLLQTSLET-PLSLESEFELGCYA------------   229
Tn7001           204   LSKYLINTDLRV-LLNNQTELKTTYV-------------   228
Tn7002           184   IARQLLSIDMHQ-PIGWQSPIWSS---------------   206
Tn7003           184   IAKKILEFDWHC-PISHDSLVWRVS--------------   207
Tn7004           185   ISSGQVKTNLNNIGFGLETYVWC----------------   207
Tn7005           185   ISSGHVKTDLNTIGFGLETCVWC----------------   207
Tn7006           185   LSSGEIQTDFNQKKFDLESSVWC----------------   207
Tn7008           184   LSKGALQTDYSNGVINGNSYVWL----------------   206
Tn7011           184   LSLGQLFADLTTNEISIETAIWSNNV-------------   209
Tn7012           184   LCIGQLATDLTTDALSLDSVIWASDEQ------------   210
Tn7013           185   IGLGRVNVPLDS-AINEMSVISVN---------------   207
Tn7014           185   IGLGKAKTKLDV-LLDENSLISVA---------------   207
Tn7015           185   INKGLLEADLTQDDLSCNPFVWCNA--------------   209
Tn7017           195   IANKIIGFDIDRYKLNQSVMVGVWNESFDRAWNGVPIQE   233
Consensus_aa:          lt..bl.hchp...ls.ps.l.hs...............
Consensus_ss:          hh  eeeeee        eeeee
```

#### Supplementary Data 2 | Multiple sequence alignment of TnsA proteins from type I-F CASTs

Output from Promals3D[6]; Tn*7016* (*Pseudoalteromonas* sp. strain S983), WP_029772101.1; Tn*6677* (*Vibrio cholerae* strain HE-45), WP_000202732.1; Tn*6900* (*Aeromonas salmonicida*), WP_088821967.1; Tn*7000* (*Vibrio cholerae* strain 4874), WP_119464977.1; Tn*7001* (*Photobacterium iliopiscarium*), WP_107274794.1; Tn*7002* (*Vibrio* sp. strain F12), WP_136985357.1; Tn*7003* (*Vibrio parahaemolyticus*), WP_015313515.1; Tn*7004* (*Vibrio* sp. strain 16), WP_005472937.1; Tn*7005* (*Vibrio cholerae* strain M1517), WP_000460251.1; Tn*7006* (*Vibrio splendidus*), WP_050622132.1; Tn*7008* (*Aliivibrio* sp.), WP_065591602.1; Tn*7011* (*Pseudalteromonas* sp. strain P1-25), WP_036981971.1; Tn*7012* (*Pseudoalteromonas ruthenica*), WP_138590901.1; Tn*7013* (*Vibrio cholerae* strain OYP7G04), WP_114778920.1; Tn*7014* (*Vibrio diazotrophicus*), WP_102952856.1; Tn*7015* (*Shewanella* sp.), WP_076538389.1; Tn*7017* (*Endozoicomonas ascidiicola*), WP_157673482.1[7]. Domain boundaries of *Pse*TnsA are indicated above the alignment; residues described in this study are highlighted in yellow.

### Supplementary Tables

#### Supplementary Table 1 | Cryo-EM data collection, refinement and validation statistics

| ID, EMDB and PDB codes | Cryo-EM structure of the *Pse*TnsAB paired-end complex (right end) in the presence of Mg (EMDB-55638) (PDB-9T7L) | Cryo-EM structure of the *Pse*TnsAB paired-end complex (left end) in the presence of Mg (EMDB-57834) (PDB-30JW) | Cryo-EM structure of the *Pse*TnsAB paired-end complex (right end) in the presence of Mn (EMDB-57833) (PDB-30JV) |
|---|---|---|---|
| **Data collection and processing** | | | |
| Magnification | 130,000 | 130,000 | 130,000 |
| Voltage (kV) | 300 | 300 | 300 |
| Electron exposure ($e^-$/Å$^2$) | 62.130 | 64.201 | 64.201 |
| Defocus range (μm) | -1.0 to -2.4 (0.2 steps) | -0.8 to -2.6 (0.3 steps) | -0.8 to -2.6 (0.3 steps) |
| Pixel size (Å) | 0.325 | 0.325 | 0.325 |
| Symmetry imposed | C2 | C2 | C2 |
| Initial particle images (no.) | 1,095,968 | 4,935,000 | 2,242,284 |
| Final particle images (no.) | 203,418 | 575,675 | 159,536 |
| Map resolution (Å) | 2.78 | 2.51 | 2.86 |
| FSC threshold | 0.143 | 0.143 | 0.143 |
| Map resolution range (Å) | 2.3-5.0 | 2.0-4.0 | 2.5-5.0 |
| **Refinement** | | | |
| Initial model used (PDB code) | AlphaFold3 | 9T7L | 9T7L |
| Model resolution (Å) | 2.8 | 2.5 | 2.8 |
| FSC threshold | 0.143 | 0.143 | 0.143 |
| Model resolution range (Å) | 2.7-3.0 | 2.5-2.7 | 2.8-3.1 |
| Map sharpening *B* factor (Å$^2$) | Unsharpened map | Unsharpened map | Unsharpened map |
| Model composition | | | |
| Non-hydrogen atoms | 17938 | 17,940 | 17,940 |
| Protein residues | 1,888 | 1,888 | 1,888 |
| Nucleotide residues | 124 | 124 | 124 |
| Ligands | MG: 2 | MG: 2 | MN: 4 |
| *B* factors (Å$^2$) min/max/mean | | | |
| Protein | 30.00/207.02/132.93 | 59.35/209.03/124.87 | 64.27/216.45/138.28 |
| Nucleotide | 80.15/258.18/140.39 | 61.36/255.48/129.51 | 69.14/273.80/145.19 |
| Ligand | 163.15/163.15/163.15 | 161.53/161.53/161.53 | 160.44/165.24/162.84 |
| R.m.s. deviations | | | |
| Bond lengths (Å) | 0.004(0) | 0.003(0) | 0.003(0) |
| Bond angles (°) | 0.625(0) | 0.602(0) | 0.563(0) |
| Validation | | | |
| MolProbity score | 1.66 | 1.55 | 1.68 |
| Clashscore | 6.35 | 7.73 | 8.08 |
| Poor rotamers (%) | 2.59 | 1.47 | 1.94 |
| Ramachandran plot | | | |
| Favored (%) | 98.55 | 98.66 | 97.90 |
| Allowed (%) | 1.45 | 1.34 | 1.99 |
| Disallowed (%) | 0.00 | 0.00 | 0.11 |

#### Supplementary Table 2 | Oligonucleotides and plasmids used in this study
